# Specialized structure of neural population codes in parietal cortex outputs

**DOI:** 10.1101/2023.08.24.554635

**Authors:** Houman Safaai, Alice Y. Wang, Shinichiro Kira, Simone Blanco Malerba, Stefano Panzeri, Christopher D. Harvey

## Abstract

Do cortical neurons that send axonal projections to the same target area form specialized population codes for transmitting information? We used calcium imaging in mouse posterior parietal cortex (PPC), retrograde labeling, and statistical multivariate models to address this question during a delayed match-to-sample task. We found that PPC broadcasts sensory, choice, and locomotion signals widely, but sensory information is enriched in the output to anterior cingulate cortex. Neurons projecting to the same area have elevated pairwise activity correlations. These correlations are structured as information-limiting and information-enhancing interaction networks that collectively enhance information levels. This network structure is unique to sub-populations projecting to the same target and strikingly absent in surrounding neural populations with unidentified projections. Furthermore, this structure is only present when mice make correct, but not incorrect, behavioral choices. Therefore, cortical neurons comprising an output pathway form uniquely structured population codes that enhance information transmission to guide accurate behavior.

## Introduction

The processing of sensory stimuli, computations for cognitive functions, and the generation of behavioral outputs all require the communication between densely interconnected brain areas to transmit information and maintain coherent functionality^1,2^. A fundamental component of neural computation is therefore how populations of neurons encode and transmit information to specific downstream target areas. Each cortical area communicates with many other areas and contains a heterogeneous population of neurons that project to distinct downstream targets^3–6^. For the transmission of information between brain areas, the relevant neural codes are likely formed by populations of neurons that communicate with the same downstream target area so that their activity can be read out as a group. However, in most studies of neural population codes, populations have been analyzed without knowledge of whether the cells project to the same target. It therefore is an open question of what principles underlie coding in populations of neurons that project to the same target area.

The information encoded in a population of neurons is strongly determined by correlations between the activity of different neurons. Much experimental and theoretical work has demonstrated how the correlations in activity between pairs of neurons can either enhance the population’s information, due to synergistic neuron-neuron correlations, or increase redundancy between neurons, which might establish robust transmission but limit the information encoded^7^. Most of this understanding arises from considerations of typical or average pairwise correlation values in populations. However, recent work has reported that pairwise correlations in large populations can take on additional network structures that could include hubs of redundant or synergistic interactions^8–10^. Yet, little is understood about how a network-level structure of pairwise correlations may contribute to the information in a neural population. Importantly, how this network-level structure influences the transmission of information between brain areas has not been studied.

We studied the population codes for transmitting information between cortical areas with a focus on the posterior parietal cortex (PPC). Much work from multiple species has identified PPC as a sensory-motor interface during decision-making tasks, including during navigation in rodents^11–17^. PPC has a heterogeneous set of activity profiles, including cells encoding various sensory modalities, locomotor movements, and cognitive signals, such as spatial and choice information^11,18–21^. PPC is densely interconnected with cortical and subcortical regions, in particular in a network containing retrosplenial cortex (RSC) and anterior cingulate cortex (ACC)^22^. In addition, population codes in PPC contain correlations between neurons that benefit behavior^23–25^. We study PPC in the context of a flexible navigation-based decision-making task because navigation decisions require the coordination of multiple brain areas to integrate signals across areas and because PPC activity is necessary for mice to solve navigation decision tasks^11,26–28^.

Here we developed statistical multivariate modeling methods to investigate the different components of population codes in cells sending axonal projections to the same target. We find that PPC distributes its sensory, choice, and locomotor information broadly to its outputs but with higher transmission of sensory information to ACC. Further, we discovered that, in PPC neurons projecting to the same target, pairwise correlations are stronger and arranged into a specialized network structure of interactions. This structure consists of pools of neurons with enriched within-pool and reduced across-pool information-enhancing interactions, with respect to a randomly structured network. This structure enhances the amount of information about the mouse’s choice encoded by the population, with proportionally larger contribution for larger population sizes. Remarkably, this information-enhancing structure is only present in populations of cells projecting to the same target, and not in neighboring populations with unidentified outputs. Such structure is present when mice make correct choices, but not when they make incorrect choices. Together, we propose that specialized network structures in PPC populations that comprise an output pathway increase signal propagation in a manner that may aid accurate decision-making.

## Results

### A delayed match-to-sample task that isolates components of flexible navigation decisions

We developed a delayed match-to-sample task using navigation in a virtual reality T-maze (Figure 1A)^26^. The beginning of the T-stem contained a black or white sample cue followed by a delay maze segment that had identical visual patterns on every trial. When mice passed a fixed maze location, a test cue was revealed as a white tower in the left T-arm and a black tower in the right T-arm, or vice versa. The sample cue and test cue were chosen randomly and independently of one another in each trial, and the two types of each cue defined four trial types (Figure 1B). The mouse received a reward when it turned in the direction of the T-arm whose color matched the sample cue. Thus, the mouse combined a memory of the sample cue with the test cue identity to choose a turn direction at the T-intersection. This process is equivalent to an exclusive OR (xor) computation for the association between the sample cue-test cue combination and the turn direction (choice) that results in a reward. After weeks of training, mice performed this task around 80% correct, on average (Figure S1A).

**Figure 1.**
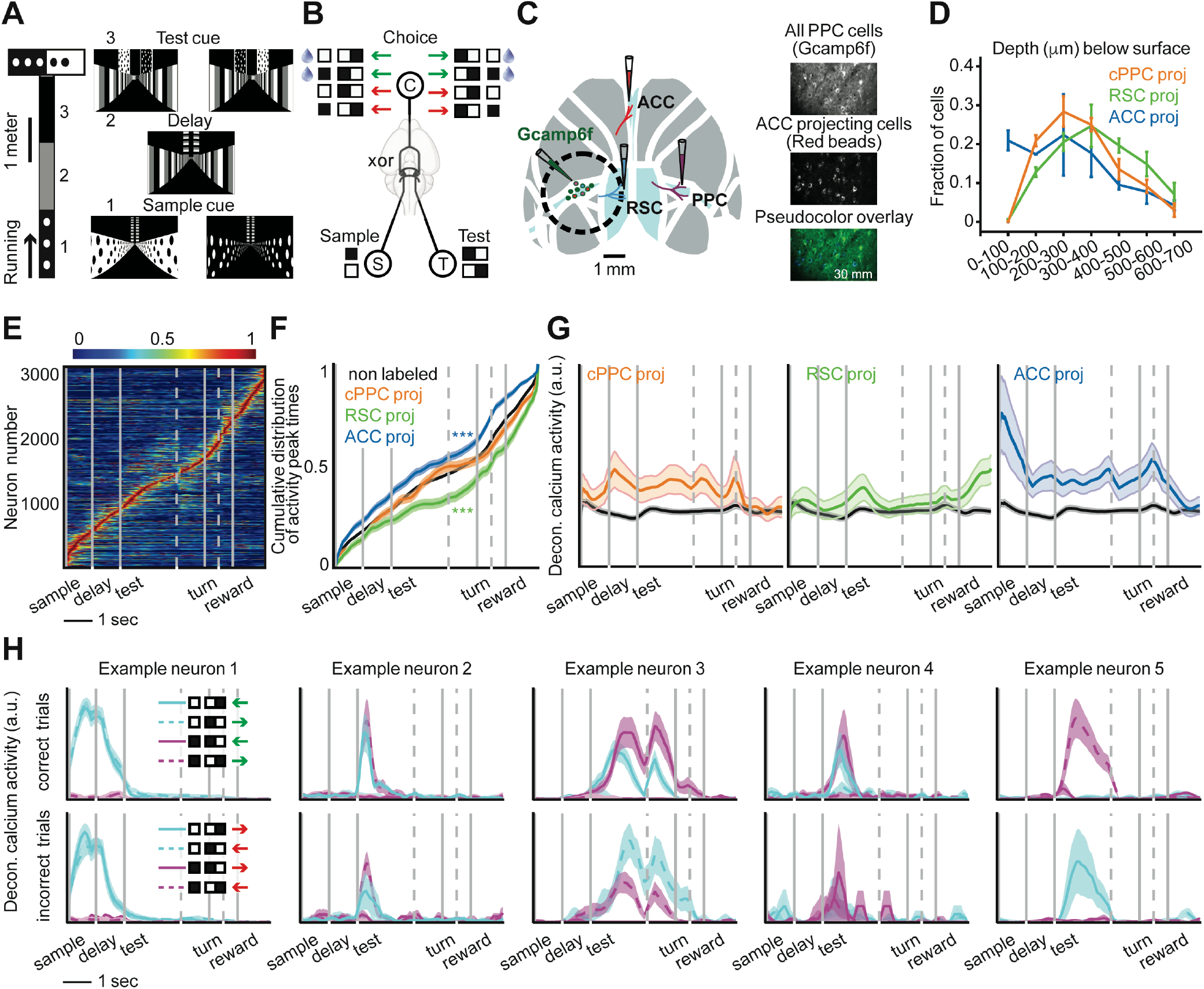
Activity differences of neurons projecting to distinct cortical targets. (**A**) Schematic of a delayed match-to-sample task in virtual reality. (**B**) The xor combination of a sample cue and test cue dictates the rewarded direction (green and red arrows represent correct and incorrect decisions respectively). (**C**) Retrograde virus injections to label PPC neurons projecting to ACC, RSC, and contralateral PPC. (**D**) Depth distribution of PPC neurons projecting to different areas. Error bars indicate mean ± SEM over mice. (**E**) Normalized mean deconvolved calcium traces of PPC neurons sorted based on the cross-validated peak time. Vertical gray lines represent onsets of sample cue, delay, test cue, turn into T-arms, and reward. Gray dashed lines correspond to one second before turn and 0.5 second before reward. (**F**) Cumulative distribution of the peak activity times for neurons. Compared to non-labeled population: p < 0.001 for ACC-projecting and RSC-projecting neurons, two-sample KS-test. (**G**) Mean ± SEM deconvolved calcium activity of different populations. Non-labeled neurons are shown in black. (**H**) Mean ± SEM deconvolved calcium activity of example neurons with different encoding properties are shown in different trial conditions. Each trace corresponds to a trial type with a given sample cue and test cue in correct (green choice arrow) or incorrect (red choice arrow) trials. Neurons 1 to 4 encode the sample cue, the test cue, the left-right turn direction (choice), and the reward direction, respectively. Neuron 5 is active on only one of the four trial conditions in correct and incorrect trials and thus encodes multiple task variables.

During the task, we used two-photon calcium imaging to measure the activity of hundreds of neurons simultaneously in layer 2/3 of PPC. We injected retrograde tracers conjugated to fluorescent dyes of different colors to identify neurons with axonal projections to ACC, RSC, and contralateral PPC (Figure 1C, Figures S1B-C). These target regions were chosen because they are major recipients of projections from layer 2/3 PPC neurons, whereas other targets, in particular subcortical areas, receive projections from deeper layers^4^. Furthermore, the ACC-RSC-PPC network has dense interconnectivity and is important for navigation-based decision tasks^28,29^. We did not have the bandwidth to examine other projection targets of PPC. The PPC neurons projecting to ACC, RSC, and contralateral PPC were largely intermingled, except the most superficial part of layer 2/3 had an enrichment of ACC-projecting neurons (Figure 1D). We did not observe cells labeled with multiple retrograde tracers.

Individual layer 2/3 neurons were transiently active during task trials with different neurons active at different time points, and the activity of the population tiled the full trial duration (Figure 1E, Figure S1D)^11^. ACC-projecting cells had higher activity early in the trial while RSC-projecting cells had higher activity later in the trial. Contralateral PPC-projecting neurons had more uniform activity across the trial (Figures 1F,G). These patterns were apparent from the distribution of cells’ peak activity times during the trial (Figure 1F) and from the time-course of mean activity of each projection type (Figure 1G).

These differences in average activity levels across the trial suggest that neurons projecting to different targets could contribute to different stages of information processing in the task (Figure S3). They could encode: the sample cue (neuron 1, Figure 1H), the test cue (neuron 2, Figure 1H), the left-right turn direction (choice) (neuron 3, Figure 1H), and the xor combination of the sample cue and test cue that indicates the reward direction (neuron 4, Figure 1H). Note that the reward direction (xor of the sample and test cues) and choice are identical on correct trials and opposite on incorrect trials. In addition to cells that encode these individual variables, we identified neurons that were active on only one of the four trial types in correct and/or incorrect trials and thus encode multiple task variables (neuron 5, Figure 1H).

### Vine copula models to analyze encoding in multivariate neural and behavioral data

To quantify the selectivity of neurons for different task variables, we aimed to isolate the contribution of a task variable to a neuron’s activity while controlling for other variables that might also contribute. This was important because neural activity is modulated by movements of the mouse^18,19,30^, and a mouse’s movements correlate with task variables (Figures S2A,C).

We adapted methods called nonparametric vine copula (NPvC) models to estimate the multivariate dependence between a neuron’s activity, task variables, and movement variables (Figure 2A). This method expresses the multivariate probability densities as the product of a copula, which quantifies the statistical dependencies among all these variables, and of the marginal distributions^31–33^. The mutual information between two variables depends only on the copula and not on the marginal distributions, which simplifies the information estimation^34^. Using a specific sequential probabilistic graphical model called the vine copula^31,33^, we broke down the complex and data-hungry estimation of the full multivariate dependencies into a sequence of simpler and data-robust estimations of bivariate dependencies (Figure 2A). Using a non-parametric Kernel-based estimator for each bivariate copula^34^, we took into account correlations between all the variables in the multivariate probability, and we avoided strong assumptions about the nature of the dependencies. The NPvC does not make assumptions about the form of the marginal distributions of the variables and their dependencies since it is based on kernel methods to estimate these probabilities. Thus, the NPvC is a convenient method to quantify information in the presence of multivariate dependencies between variables while minimizing the risk of potentially imposing invalid assumptions on the data. Also, because it models the whole dependency structure, the NPvC is a powerful tool to discount possible collinearities when studying information encoding.

**Figure 2.**
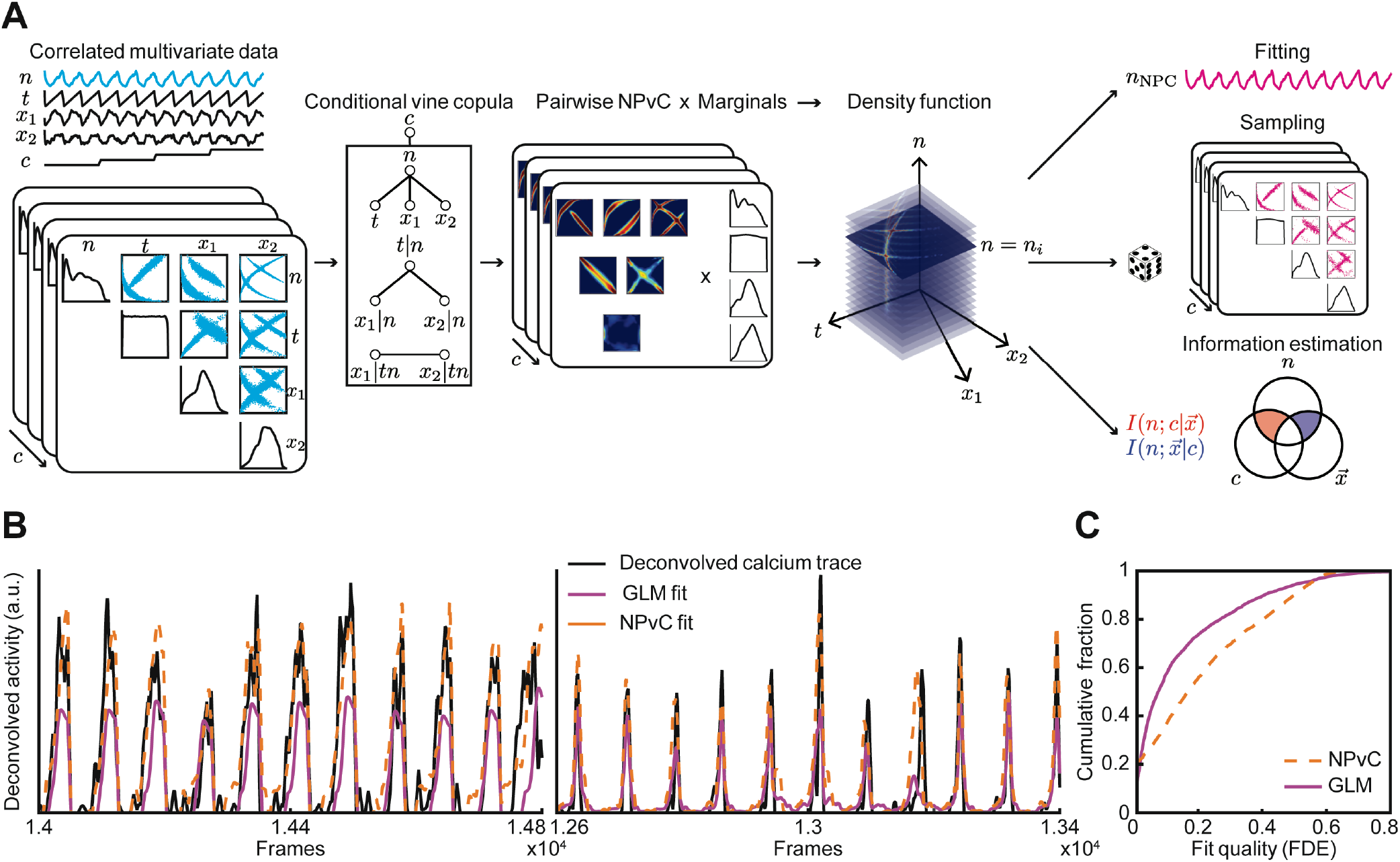
Vine-copula modelling of neural activity. (**A**) Schematic of nonparametric vine copula model (NPvC) of neural activity (*n*) as a function of time (*t*), a vector of movement variables 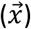 with components (*x*_1_, …, *x*_*N*_), and task variable (*c*). Conditional vine copulas are built between neural activity and all the other variables for each task variable (*c*). Mixing the vine copula and marginal distributions gives the conditional density function of neural activity and other variables. The vine copula model can be used either to estimate the value of neural activity conditioned over all the other variables (which is the copula fit *n*_*NPC*_), or to generate samples, or to estimate various conditional entropy and mutual information values. (**B**) Deconvolved calcium activity of two example neurons (black) and the fits of the vine copula model fit (orange, dashed line) and GLM (pink, solid line). Cumulative distribution of fraction of deviance explained (FDE) across neurons for the GLM and the NPvC model.

Using our NPvC model, we estimated the expected activity of a neuron for any given value of task and movement variables and at any time in the trial (Figure 2B). We then validated the model’s performance by quantifying the fraction of deviance explained on held-out test data. The NPvC fitted frame-by-frame neural activity better than a generalized linear model (GLM), which is commonly used for these types of analyses (Figure 2C)^23,35,36^.

We used the NPvC model to estimate the mutual information between a neuron’s activity and each task variable at each time point. By using the NPvC model, we accounted for potential covariations between the measured task and movement variables in the estimates of information. Further, in our calculation of the information carried by neural activity about each variable, we conditioned on the values of all other measured variables to remove the effect of the correlations between task and movement variables^18,23^.

### Preferential, but widespread, routing of information

The population of PPC neurons contained information about each of the task variables, even after conditioning on the movement variables (Figure 3A). Sample cue information was high in the sample, delay, and test segments. Both sample cue and test cue information were appreciable in the early part of the test segment when the cues needed to be combined to inform a choice (Figure 3A, left). PPC neurons carried information about the reward direction (xor combination of the sample and test cues) and the choice, but the choice information was larger, indicating that PPC activity was more related to the turn direction selected by the mouse than the reward direction defined by the cues (Figure 3A, middle). In addition, PPC contained information about the movements of the mouse throughout the trial (Figure 3A right).

**Figure 3.**
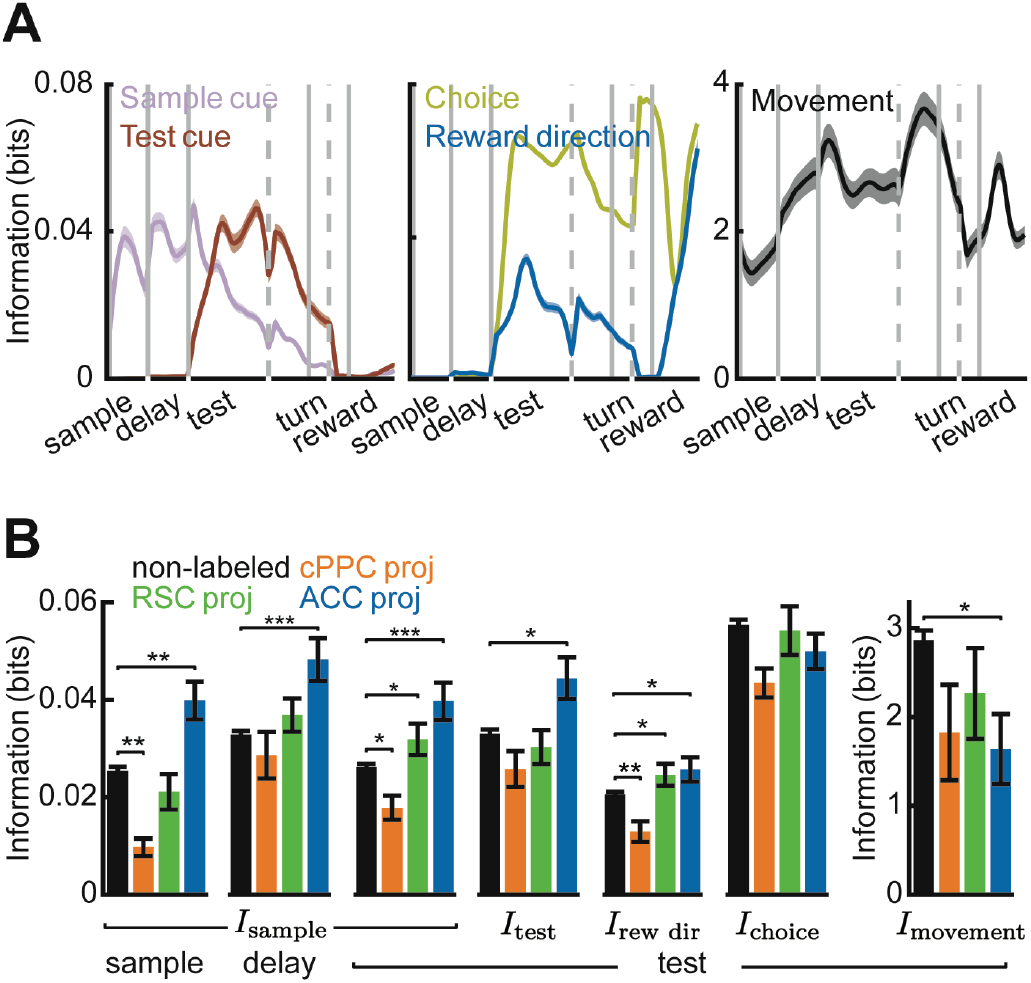
Single-neuron information in labeled projection neurons and non-labeled cells. (**A**) Time-course of different information components in all PPC neurons. Shading indicates mean ± SEM. (**B**) Average single-neuron information in different populations about different task variables during the first two seconds after sample cue onset, delay onset, or test cue onset (non-labeled n = 2185 cells, cPPC proj: n = 76 cells, RSC proj: n = 108 cells, ACC proj: n = 84 cells). Error bars indicate mean ± SEM across cells. * indicates p < 0.05, ** indicates p < 0.01, and *** indicates p < 0.001, t-test with Holm-Bonferroni correction for statistical multiple comparisons.

To determine if different aspects of task and movement information are transmitted to distinct targets, we compared neurons with different identified projections and the unlabeled cells with unidentified projections. Information for the sensory-related task variables – sample cue, test cue, and their xor combination – was enriched in ACC-projecting neurons and lowest in contralateral PPC-projecting cells (Figure 3B). Thus, PPC preferentially transmits sensory information to ACC. In contrast to sample cue and test cue information, information about the choice and movements was similar across the projection types, indicating that this information is more uniformly transmitted (Figure 3B). However, all three projection types had lower information about the movements of the mouse than the unlabeled cells, suggesting that the movement information is enriched in neurons projecting to areas other than those studied here or in interneurons (Figure 3B, right). In addition, cells projecting to contralateral PPC often had less information about each variable than the unlabeled cells, indicating that across-hemisphere communication may be less critical for encoding specific task and movement events. On the other hand, RSC-projecting neurons carried the information typical of the PPC population, as shown by similar levels of information to the unlabeled neurons. Thus, neurons projecting to different target areas differ in their encoding, revealing a specialized routing of signals from the PPC to its targets. However, each projection class contains a significant level of information about each variable, showing that PPC also broadcasts its information widely.

### Enriched information-enhancing pairwise interactions in neurons projecting to the same target

Beyond single-neuron coding, the structure of correlated activity patterns in populations of neurons can impact the transmission and reading out of information^7,37^. We used the NPvC model to calculate pairwise noise correlations^7,37^, defined as the correlations in activity for a pair of neurons for a fixed trial type (Supplemental Information, Figure 4A). We focused on the first two seconds after the test cue onset. Remarkably, noise correlations were significantly larger in pairs of neurons projecting to the same target than in unlabeled neurons with unidentified projection patterns (Figure 4B left; Figure S4A), suggesting that correlations aid transmission of information between areas. For all groups of neurons, noise correlations were higher on correct trials than incorrect trials, consistent with the possibility that correlations aid the transmission and reading out of information to guide behavior^24^ (Figure 4B; Figure S4A). We also considered that behavioral variability within a given trial type, such as differences in running velocities, could contribute to trial-to-trial variability and thus potentially to noise correlations. After conditioning on movement variability using the single-neuron NPvC models, noise correlations were lower, confirming that movement variability contributed to traditional noise correlation measures (Figure 4B right, Figure S4A). However, even in this case, noise correlations were higher in neurons projecting to the same target than in pairs of unlabeled neurons and higher in correct trials (Figure 4B right, Figure S4A).

**Figure 4.**
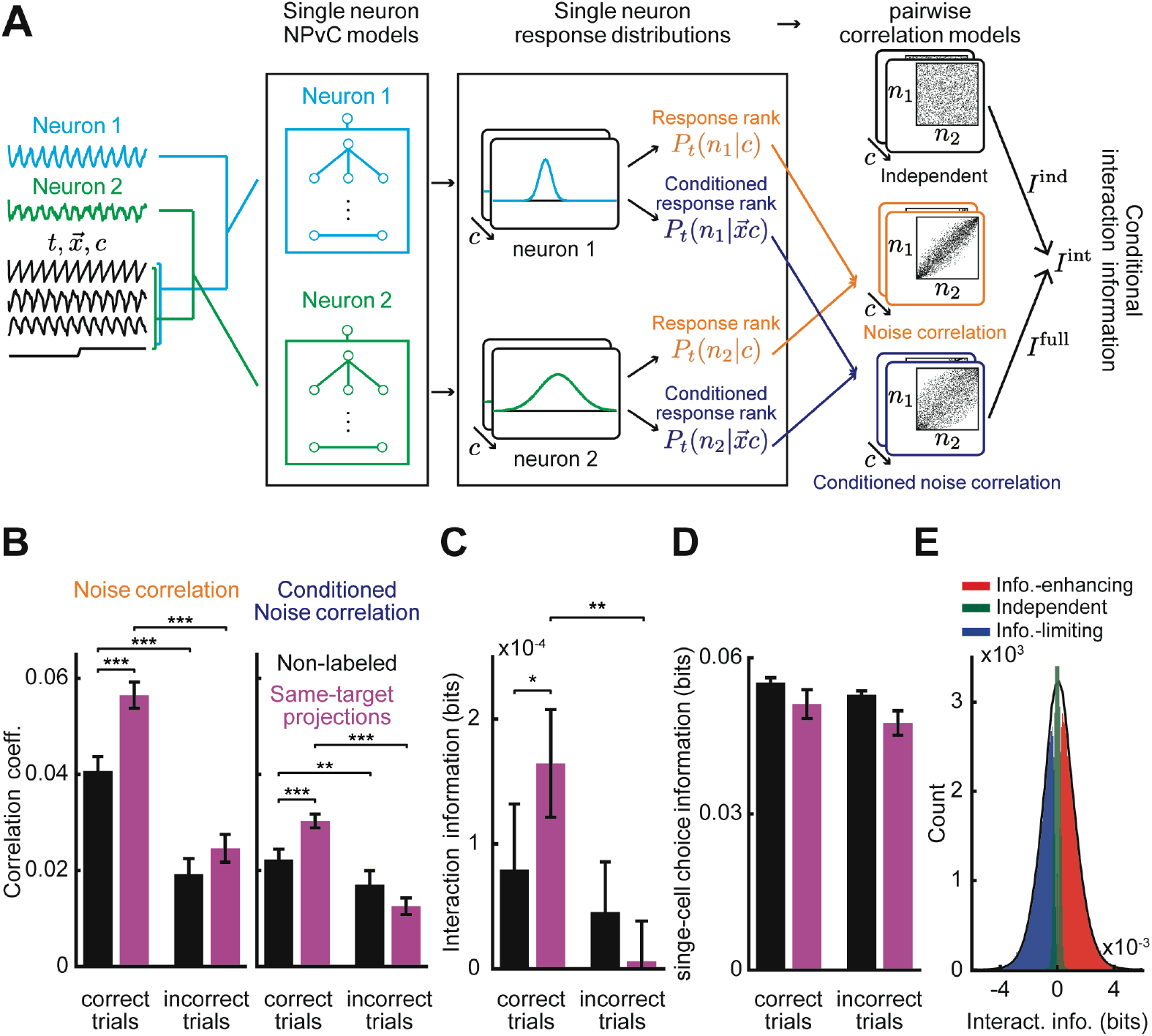
Pairwise interactions between pairs of non-labeled neurons and pairs of neurons projecting to the same target. (**A**) Schematic of the models to compute pairwise joint probability density functions and conditional joint probability density functions of two neurons with correlated activities (*n*_1_, *n*_2_) as a function of time (*t*). A vector of movement variables 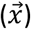 with components (*x*_1_, …, *x*_*N*_) is represented. Using single neuron NPvC model outputs, we build different types of pairwise correlation models with and without conditioning over the movement variables. The joint pairwise model is then used to estimate noise correlations or interaction information. (**B**) Left: Noise correlations computed for pairs of non-labeled neurons and pairs of neurons projecting to the same area for correct and incorrect trials. Right: Same except for noise correlations conditioned on movement variables. (**C**) Similar to panel (B) but for interaction information. (**D**) Average single-neuron choice information in different populations during the first two seconds after the test onset for correct and incorrect trials. (**E**) Histogram of interaction information divided into pairs of information-enhancing (red), information-limiting (blue), and independent pairs (green). In panels (B-D), error bars indicate mean ± SEM across all pairs of neurons. * indicates p < 0.05, ** indicates p < 0.01, and *** indicates p < 0.001, t-test with Holm-Bonferroni correction for statistical multiple comparisons.

Depending on well characterized relationships between the structure of signal and noise correlations, noise correlations can either limit or enhance the information in neural populations^7,37^. To quantify the effect of noise correlations on information, for each pair of neurons, we decomposed the mutual information between the pair’s activity and a task variable into independent information, defined as the information that the pair would carry if their noise correlations were absent, and interaction information, defined as the difference between the actual information carried by the pair (which includes the effect of noise correlations) and the independent information (Figure 4A). The interaction information thus quantifies how much a neuron pair’s noise correlations increase (information-enhancing) or decrease (information-limiting) information about a task variable. We focused on choice information because it is relevant for guiding behavioral actions and in PPC is larger than sample cue and test cue information.

The average value of interaction information across pairs of neurons was positive, and thus noise correlations were on average information-enhancing (Figure 4C). Remarkably, interaction information on correct trials was larger in pairs of neurons projecting to the same area compared to pairs of unlabeled cells (Figure 4C). Furthermore, interaction information was higher on correct trials than on incorrect trials. For pairs of neurons projecting to the same target, interaction information even dropped close to zero on incorrect trials (Figure 4C). In contrast, in single neurons, the choice information was on average similar in value between correct and incorrect trials (Figure 4D), suggesting a specific role of neuron-to-neuron interactions for generating correct behavioral choices. These results indicate that the pairwise interactions differ in populations projecting to the same target relative to surrounding neurons, that these interactions enhance the information that is transmitted, and that they may be beneficial for accurate decisions.

Interestingly, there was a wide range of interaction information values across pairs of neurons, with extended and almost symmetric positive and negative tails (Figure 4E). We identified pairs of neurons with information-enhancing (significantly positive), information-limiting (significantly negative), and independent (not significantly different from zero) interaction information. Most pairs (∼90%) were classified as information-enhancing or information-limiting (Figure S4E). Confirming previous studies^37–39^, the most important determinant of whether a pair had information-enhancing or information-limiting interaction information was the sign of noise correlations relative to the sign of signal correlations (i.e., the similarity of choice selectivity of the two neurons forming a pair). Pairs with the same sign for signal and noise correlations had information-limiting interactions, whereas pairs with opposite signs for these correlations had information-enhancing interactions (Figures S4H,I). Thus, the PPC population has a rich distribution of information-enhancing and information-limiting pairs.

We also computed the interaction information for individual projection targets and about the sample cue and test cue (Figures S4B,D). RSC-projecting pairs had similar interaction information to unlabeled pairs, consistent with RSC-projecting neurons being most similar to the unlabeled population. For test cue information, we observed similar but weaker results to those reported above for choice. For sample cue information, we did not find differences in interaction information between cells projecting to the same target and unlabeled pairs. Thus, there exists possible specificity of interactions between these neurons with respect to different task variables and projection pathways.

### The network structure of pairwise interactions

Two networks that have the same set of values of information-limiting and information-enhancing interactions can differ in how these interactions are organized within the network (Figures 5A,B). For example, the same set of interaction pairs can be distributed either randomly within a group of neurons (Figure 5A, top) or structured as clusters containing enriched information-enhancing or information-limiting interactions (Figure 5A, middle and bottom).

**Figure 5.**
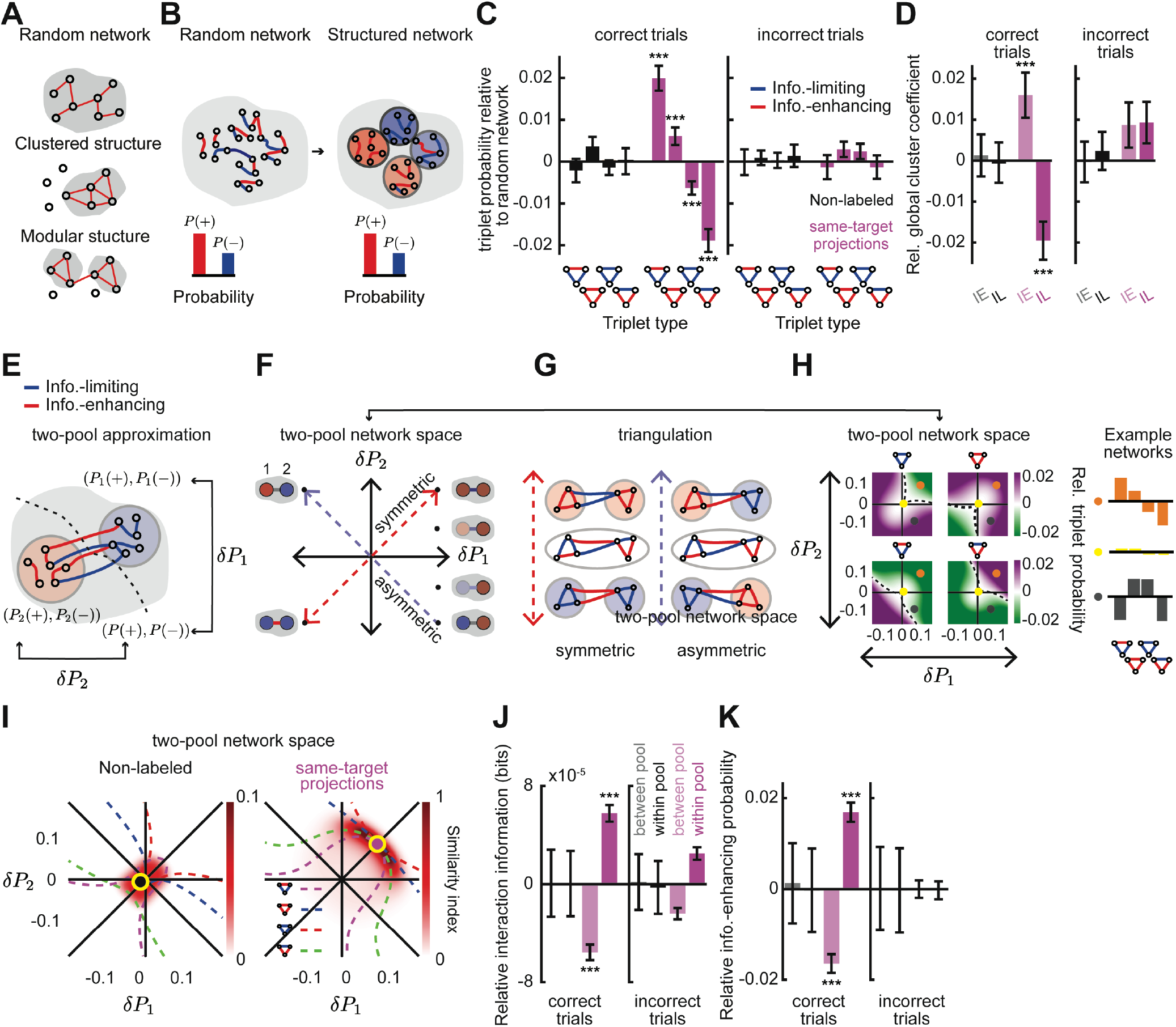
The presence of network structure of interaction information in populations projecting to the same target. (**A**) Schematics of a random network (top), a network with clustered interactions (middle), and a network with modular structure (bottom). (**B**) Sketch of how networks with the identical probability of information-enhancing and information-limiting interactions can be organized randomly or in clusters of information-enhancing or information-limiting pairs. Red and (+) indicates information-enhancing pairs. Blue and (-) indicates information-limiting pairs. (**C**) Relative triplet probability with respect to a random network for non-labeled and same-target populations during correct (left) and incorrect (right) trials. The random network has the same distribution of pairwise interactions, except shuffled between neurons. (**D**) Global cluster coefficient of information-enhancing (IE) and information-limiting (IL) sub-networks relative to a random network. (**E**) Schematic of a two-pool network model. (**F**) Schematic of the space of two-pool networks is quantified in terms of the probability of information-enhancing pairs in pool 1 (x-axis) and pool 2 (y-axis) minus the probability of these pairs in a random network. Red indicates information-enhancing pools, and blue indicates information-limiting pools. (**G**) Schematic of some examples of symmetric and asymmetric networks corresponding to different points in the two-pool network space along the diagonal (red dashed line) or antidiagonal (blue dashed line) axes. (**H**) Left: The probabilities of different triplets computed for different two-pool networks minus the triplet probabilities in a random network, computed analytically as derived in Supplemental Information: Analytical calculation of triplet probabilities in the two-pool model of network structure for two-pool networks. Right: The triplet probabilities for three example model networks sampled from the space of two-pool networks. At each point in the 2D space of two-pool networks, comparison of the triplet probabilities for the model network and the empirical data. The similarity index ranges from 1 for similar networks to 0 for completely different networks. Yellow circles correspond to networks that are more similar to data for non-labeled (left) and same-target projection networks (right). The four dashed contours correspond to two-pool networks with similar value of each of the four triplet probabilities to the one obtained from data (from Figure 5C for correct trials). (**J**) Using pools of neurons defined based on their choice selectivity to left or right choices, the average interaction information within the pool and between the pools, relative to a random network. Pink indicates same-target projections. Black indicates non-labeled neurons. (**K**) Similar to (**J**) but for the probability of information-enhancing pairs within or between pools, relative to a random network. Error bars indicate mean ± SEM estimated using bootstrapping over all triplets. *** indicates p < 0.001, t-test with Holm-Bonferroni correction for statistical multiple comparisons.

Graph-theoretic measures can be used to identify structured arrangements of pairwise links in a network. The simplest measures are defined based on the frequency of interaction triplets, which are the simplest motifs in a graph beyond pairs^40,41^. For example, a network with information-limiting clusters will have a larger number of ‘-,-,-’ triplets compared to a random network, and a network with information-enhancing clusters will have more ‘+,+,+’ triplets compared to a random network (Figures 5A,B), where ‘-’ and ‘+’ correspond to information-limiting and information-enhancing pairwise interaction links, respectively. For interactions for choice information, we identified the network structure by computing the difference in the probabilities of ‘-,-,-’, ‘-,-,+’, ‘-,+,+’, and ‘+,+,+’ triplets between our data and an unstructured network obtained by randomly shuffling the position of pairwise interactions within the network, without changing the set of values of interactions within the network. In the unlabeled population, triplet probabilities were like those in a random network, indicating that the pairwise interactions are not structured (Figure 5C). However, in populations of neurons projecting to the same target, there was an enrichment of ‘+,+,+’ and ‘-,-,+’ triplets and a relative lack of ‘-,-,-’ and ‘-,+,+’ triplets compared to a random network. Notably, this structure was present only on correct trials and was absent when mice made incorrect choices (Figure 5C). The network structure was present in ACC-, RSC-, and contralateral PPC-projecting populations for choice information but was less apparent for sample cue and test cue information (Figure S5A,B).

We next used a graph global clustering coefficient^40,41^ to measure information-limiting or information-enhancing clusters. This coefficient measures the ratio between the number of specific types of closed triplets and the number of all triplets, normalized to the same quantity computed from shuffled networks. It thus measures the excess of closed triplets in real data compared to the shuffled network. In populations of neurons on correct trials, the clustering coefficient obtained from data was larger compared to a shuffled network for information-enhancing interactions, meaning that these interactions were clustered together. In contrast, for information-limiting interactions, the clustering coefficient was smaller than for a shuffled network, indicating that they were preferentially set apart (Figure 5D). This clustering was mostly absent in unlabeled neurons and when mice made incorrect choices (Figure 5D). These results thus reveal the presence of pairwise interaction clusters in the network of neurons projecting to the same target.

To exemplify how the network’s topology relates to the triplet distributions, we considered a simple two-pool network model in which the network structure can be varied parametrically, while keeping the overall set of values for information-enhancing and information-limiting interactions constant (Figure 5E). Each pool was parameterized by the difference in probability of information-enhancing pairwise interactions within the pool relative to a random network. Thus, the range of possible network models resides in a two-dimensional space consisting of the enrichment of information-enhancing interactions in pool 1 along one axis and in pool 2 along the second axis (Figure 5F). In this parameterization, the diagonal corresponds to symmetric networks, in which both pools have more information-enhancing or more information-limiting interactions than expected in a random network. The antidiagonal corresponds to asymmetric networks, with one pool having more information-enhancing interactions and the other having more information-limiting interactions. Networks near the origin are similar to a random network.

For every point in the two-dimensional space of network models, we analytically computed the triplet probabilities compared to a random network (Figure 5H, left). We then used these model predictions of triplet probabilities to map the triplet probabilities estimated in our empirical data back into the parametric model (Figure 5I). Populations of cells projecting to the same target mapped to a symmetric network with both pools having enriched within-pool information-enhancing interactions and elevated across-pool information-limiting interactions (Figure 5I, right). In contrast, the population of unlabeled neurons mapped to a point close to the origin, corresponding to a randomly structured network (Figure 5I, left).

What factors define the neurons that participate in the pools with enriched within-pool information-enhancing interactions and elevated across-pool information-limiting interactions? One commonly studied factor is the selectivity of neurons. We divided neurons into two pools based on their choice selectivity (i.e., left vs. right choice selectivity) and compared these pools to a network with shuffled pairwise interactions. For unlabeled neurons, the interaction information and proportion of information-enhancing interactions were similar to the random network both within pools of neurons with similar choice preferences and across pools with different preferences (Figures 5J,K). In contrast, for neurons with the same projection target, pools of cells with the same choice preference had higher interaction information and a larger proportion of information-enhancing interactions than the random network (Figures 5J,K). Further, in these same-target projection populations, cell pairs with opposite choice preferences had lower interaction information and an enrichment of information-limiting interactions. This structure was strong on correct trials and largely absent when mice made incorrect choices. Thus, the network structure we described above was present in pools of neurons defined by choice preference.

Together, these results reveal rich structure in the pairwise interaction information that is approximated by a network of symmetric pools with enriched within-pool information-enhancing interactions and across-pool information-limiting interactions. This structure can be in part conceptualized in terms of organization according to the neuron’s choice selectivity. Strikingly, this structure was only present in neurons projecting to the same target, not in neighboring unlabeled neurons, and only when mice made correct choices, suggesting this structure could be significant for the propagation of information to downstream targets in a manner that is important for accurate behavior.

### The contribution of the network structure of pairwise interactions to population information

What are the consequences of this network-level structure of pairwise interactions on the information encoded in a neural population? To address this question quantitatively, we analytically expanded the information encoded by the population about a given task variable as a sum of contributions of interaction graph motifs. Under the assumption that single-pair interaction information values are small compared to single-neuron information (as is the case in our empirical PPC data; compare Figures 4C,D), the population information can be approximated as a sum of, in order of increasing complexity (Figure 6B), contributions of nodes (single neuron information), links (pairwise interaction information), and triplets (triplet-wise arrangements of pairwise interaction information). In Figures 6B-D, we illustrate this expansion for the simple case of a symmetric network, which describes well our empirical data. However, for calculating information, we used a complete and general expansion of population information that is valid also for non-symmetric networks (Supplementary Methods and Figure S7D).

**Figure 6.**
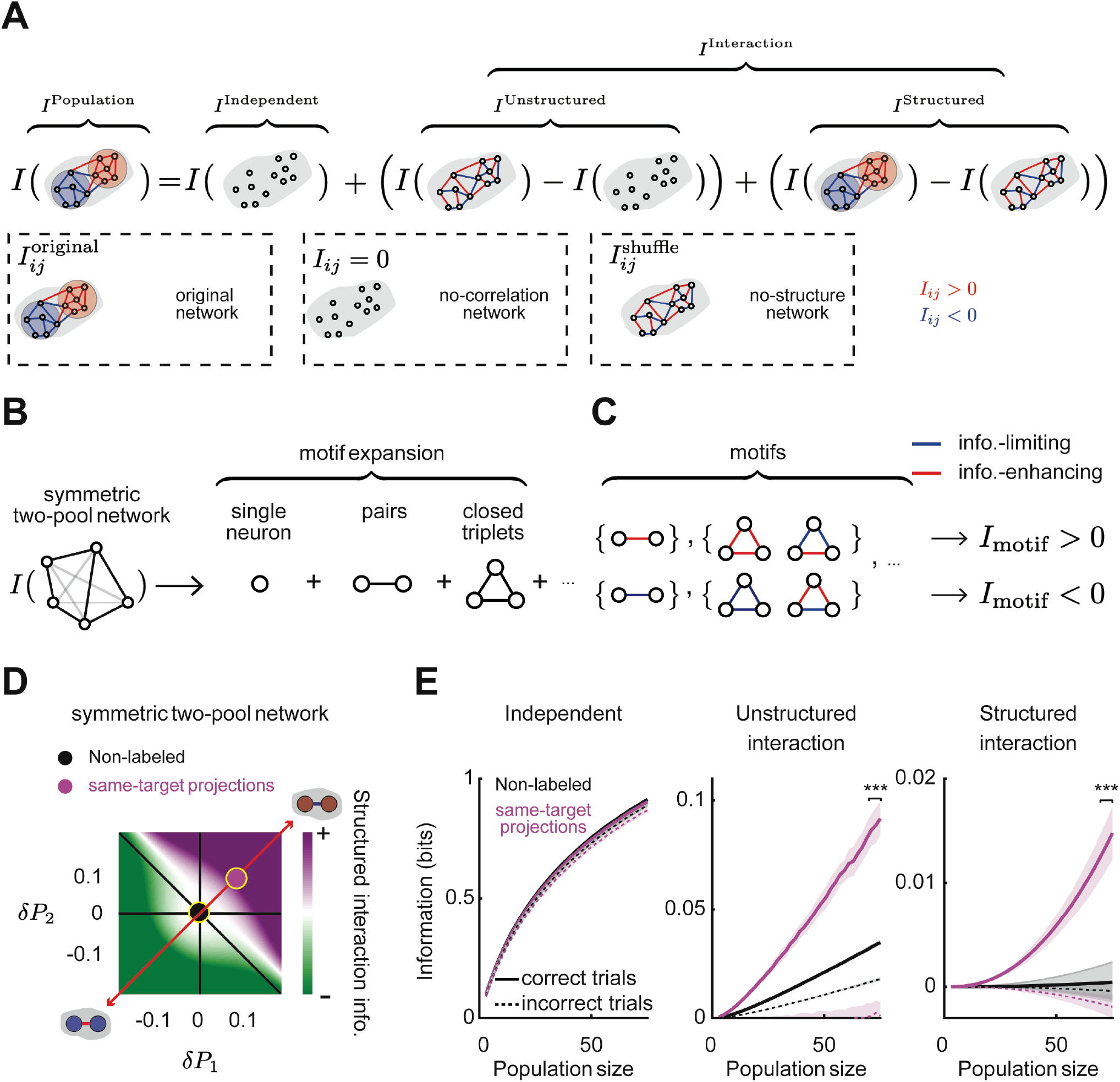
The contribution of network structure of interaction information to the population information. (**A**) Schematic of how the population information can be decomposed into three components: independent information, unstructured interaction information, and structured interaction information. (**B**) The population information can be expanded as a sum of contributions from network motifs with increasing complexity. For a symmetric network, the dominant contributions are from single neurons, interaction pairs, and interaction triplets. (**C**) In symmetric networks, triplets contribute to structured interaction information with a positive or negative sign if they have an even or odd number of information-limiting interactions, respectively. (**D**) Structured interaction information in the space of symmetric two-pool networks. The structured interaction information can be information-enhancing or information-limiting depending on whether the pools have enriched information-enhancing or information-limiting pairs. Pink and black points correspond to the networks of same-target projection and non-labeled neurons, respectively. (**E**) Independent, unstructured interaction, and structured interaction information for non-labeled (black) and same-target projection (pink) neurons during correct and incorrect trials for increasing population size. Shading indicates mean ± SEM estimated from bootstrapping over triplets. *** indicates p < 0.001, t-test with Holm-Bonferroni correction for statistical multiple comparisons at n = 75 population size.

We then used the results of this expansion to break down the total information about the task variable carried by the population into three components (Figure 6A), quantifying how pairwise correlations and its structure shape information encoding in the population. The independent information is the information that would be encoded if the population had the same single-neuron properties but zero pairwise noise correlations ^7,38,42,43^. The interaction information, which is the difference between the total population information and the independent information, was further broken down into two components. The unstructured interaction information component is the information in a population with the same values of pairwise interactions as the data, but randomly rearranged across neurons by shuffling (thus no network structure), minus the independent information. This component quantifies the contribution of the values of pairwise interactions to the population information. The structured interaction information component is defined as the difference between the total population information of the original network and the population information in an unstructured network (Figure 6A). This component isolates the contribution of the structure of pairwise interactions in the network.

The structured interaction information depends on the triplet-wise arrangements of pairwise interaction information because triplets recapitulate the network-level organization of pairwise interactions. Based on analytic calculations presented in the Supplemental Information, each triplet contributes to the structured interaction information with a sign that depends on the product of the signs of its pairwise interactions (Figure 6C). There is a positive contribution for {‘+,+,+’, ‘-,-,+’} triplets and a negative contribution for {‘-,-,-’, ‘-,+,+’} triplets. The sign of the structured interaction information component therefore depends on the difference in the probability of triplets in the original network relative to a shuffled network lacking structure. The magnitude of the structured interaction information depends on these triplet probabilities as well as the single neuron information and pairwise interaction values contained within the triplets.

To visualize how structured interaction information changes with the topology of the network, we computed the value of the structured interaction information component in the simple two-pool network defined in the previous section, making the additional assumption that the values of single neuron information and the absolute values of interaction information per triplet are homogeneous across the network (Figure 6D). In the case of a symmetric network, networks with enriched information-enhancing pools (compared to a randomly structured network) have structured interaction information that is information-enhancing (Figure 6D). In contrast, symmetric networks with enriched information-limiting pools (compared to a randomly structured network) have negative structured interaction information (Figure 6D).

Then, using the single neuron information, pairwise interactions, and triplet probabilities estimated from our PPC data, we computed the independent, the unstructured interaction, and the structured interaction components of the population information for choice (for a symmetric network (Figure 6E) and for a general non-symmetric network (Figure S7D)). The independent information was the largest component and was similar between neurons projecting to the same target and unlabeled populations (Figure 6E, left). The unstructured and structured interaction information contributed less to population information, as expected from the findings reported above that pairwise interaction information values were smaller than single-neuron information values (c.f., Figures 4C,D). Strikingly, the contributions of both structured and unstructured interaction information were markedly larger for neurons projecting to the same target than for unlabeled neurons (Figure 6E, middle, right). The larger unstructured interaction information for populations projecting to the same target is a consequence of their higher pairwise interaction values. The higher structured interaction information for populations projecting to the same target is instead a consequence of the network structure that they exhibit (Figure S7E). The structured interaction information contributed close to zero in the unlabeled population, consistent with the lack of network structure in the unlabeled population. The structured interaction information in the cells projecting to the same target was positive and information-enhancing, consistent with this population having a symmetric network structure with enriched within-pool information-enhancing interactions and across-pool information-limiting interactions compared to a randomly structured network.

Importantly, both mathematical calculations (see Supplementary Information) and numerical evaluations (Figure 6E) show that the contribution of structured interaction information scales faster with increasing population size than the two other components, indicating that this structured interaction information could be an important factor in populations of hundreds of neurons, as are likely involved in biological computations (Figure 6E). In addition, the structured interaction information was mostly absent on incorrect trials, whereas independent information was largely unchanged, suggesting that the structure of network interactions may play a role in guiding accurate behavior.

We found similar trends for network structure for choice information for the populations of neurons projecting to contralateral PPC, RSC, and ACC (Figure S6C). Consistent with the differences in the triplet probabilities for sample cue and test cue information with respect to the choice information (Figure S5A), we found that the network information is either information-enhancing or information-limiting when computed for sample cue or test cue information (Figure S6B), suggesting specificity of network structure with respect to the information content.

Together, these results reveal a rich structure in the pairwise interactions within a population, providing significant insight into the population code compared to models that only consider the mean values of pairwise correlations. Strikingly, this network structure enhanced information levels and was only present in cells projecting to the same target and when mice made correct choices. Thus, populations of PPC cells projecting to the same target are organized in a specialized manner to enhance the propagation of information to downstream targets, potentially impacting accurate decision-making.

## Discussion

The understanding of how the properties of population codes emerge from the interactions among neurons has been built on studies of information coding in populations of all neurons recorded from a given location, without identifying those that may be read out as a group. Because populations of nearby cortical neurons project to a wide diversity of targets, these previous studies have probably mixed neurons that are read out by different downstream networks. Instead, by studying population codes from the perspective of neurons that project to the same target area, we discovered properties that are unique to these populations and absent when considering all recorded neurons in PPC, including stronger pairwise correlations and a structured manner in how pairwise correlations are organized within the population. Thus, neurons comprising an output pathway in PPC form specialized population codes.

A first specialized feature of codes of populations projecting to the same target area is an elevated strength of pairwise correlations, with respect to neighboring neurons. Because correlated activity in PPC and other areas supports reliable propagation of information^23,24,44^ and because correlations are stronger in correct trials, elevated correlation levels may be present to support behavior by enhancing the efficacy of transmission of information to downstream areas. A second specialized feature of codes of populations projecting to the same target area is the presence of a network-level structure of pairwise interactions that enhances the population’s information about the mouse’s choice, even though it is made up of a diverse mix of information-enhancing and information-limiting pairwise interactions. While the information provided by the network structure is small relative to single neuron information values in small populations of neurons, we estimate that it grows rapidly with population size and provides a substantial fraction of the information carried by the whole population projecting to a specific area. In addition, the network-level structure is present only when mice make correct choices during the task and is absent when mice make incorrect choices, suggesting that the information carried by this structure is key to correct behavior. Whereas some previous work has suggested that populations must trade off the disadvantage of correlations for limiting information levels with the benefit for robust signal propagation^7,24,26,44–47^, the output pathways we studied might use this specialized interaction structure in population codes to simulatenously optimize multiple goals, including information encoding and information transmission. It will be of interest to understand how these specialized features of population codes arise from circuit connectivity. It is possible that structured local connections between cells in the same output pathway play a critical role^48^.

Previous work has focused on how pairwise and higher-order correlations shape the information encoded in a population, including establishing principles for how signal and noise correlations combine to give rise to pairwise information-limiting or information-enhancing interactions^7,37,49–53^. However, these effects have mainly been studied without taking into account how these interactions are arranged as a network. Networks with identical sets of values of pairwise interactions can have these interactions arranged in a random manner or with various types of structures. Recent work has moved in this direction finding that network-level structure exists, with redundant and synergistic interactions tending to form separate hubs in visual cortex^8^. Other work has investigated the structure of functional connectivity with graph theory or network science and revealed how principles of network organization (e.g., rich-club structure) relate to aspects of network dynamics, such as speed or robustness^54^. In many cases, it remains unstudied if and how these principles of network organization enhance or limit the encoding of information, especially in the context of decision-making tasks. Yet other work has begun to develop methods to look at the communication between brain areas based on larger scale population activity^2,55,56^. However, connections between the population geometries revealed with these approaches, the structure of pairwise correlations, and information encoding remain to be investigated. Our results build on these previous lines of research by directly identifying neurons comprising an output pathway and demonstrating how the existence of structured task-relevant information interactions in these populations impacts information encoding. In addition, we established a general framework to link network-level structures to its information-enhancing or information-limiting effect for population codes. It will be of interest to apply this approach to other communication pathways and types of tasks and behavioral information. Here, we found that the network-level structure was present across projection pathways for choice information but was less strongly apparent for other types of information.

Our analysis was facilitated by the development of NPvC models that can estimate the relationships between variables in a large multivariate setting even if the form of these relationships is unknown or complex (Figure S4G). Because these models described the neural activity better than simpler models such as GLMs, we could more effectively discount potential covariations between task and movement variables in the estimates of population information. This facilitates the removal of the covariations due to the common tuning to movement variables of many neurons^18,19,30,57^, which may induce large covariation-based redundancies masking the structure of the underlying information-enhancing and information-limiting interactions. Further development and application of copula-based methods to neuroscience will aid the analysis of relationships in high dimensional behavioral and neural data, including the structure and information content of neural correlations in larger scale neural recordings^58^.

PPC broadcasts information about the task variables to all the targets we studied here. While there was large overlap in the encoding of ACC-, RSC-, and contralateral PPC-projecting neurons, we observed some specificity in the types of information transmitted in these pathways. Most notably, sensory-related information was strongest in the ACC-projecting neurons, consistent with streamlined sensory and motor communication pathways between parietal and frontal cortical areas^59^. In contrast, contralateral PPC projections tended to carry less information about the task, implying that cross-hemisphere interactions may serve a role other than computing specific task quantities, such as choice, and instead may be more involved in maintaining consistent activity patterns^60^. All three intracortical projection pathways had lower information about the movements of the mouse, indicating that cells projecting to other targets, perhaps subcortical areas, might be more related to the biasing of movements. This finding could be consistent with corticostriatal projections from PPC carrying action selection biases^61^. Collectively, these findings support the notion of specificity in the information flow in cortex^62–65^. Simultaneously, the presence of information about each task and movement variable in each output pathway fits with many findings of highly distributed representations across cortex and with work that has failed to identify categories of neurons in PPC based on activity profiles^18,19,21,29,30,65–67^.

Together, our findings demonstrate that specialized network interaction structures in PPC aid the transmission of choice signals to downstream areas, and that these structures are potentially important for guiding accurate behavior. In addition, our results suggest that the organization principles of neural population codes can be better understood in terms of optimizing transmission of encoded information to target projection areas rather than in terms of encoding information locally.

## Acknowledgements

We thank members of the Harvey lab and Veronika Koren for helpful discussions. This work was supported by NIH grants DP1 MH125776, R01 NS089521, R01 NS108410, and the Foundation Bertarelli.

## Author contributions

HS, AYW, SP, CDH conceived of the study. AYW performed all experiments and processed the data. HS performed all data analysis and developed the analysis methods, information expansions, and models. SBM contributed to computing information expansions. CDH oversaw the experiments. SP and CDH oversaw data analyses and development of analysis and modeling methods. SK provided input on data processing and analysis. HS, SP, CDH wrote the paper with input from SK and SBM.

## Supplemental Figures

**Figure S1.**
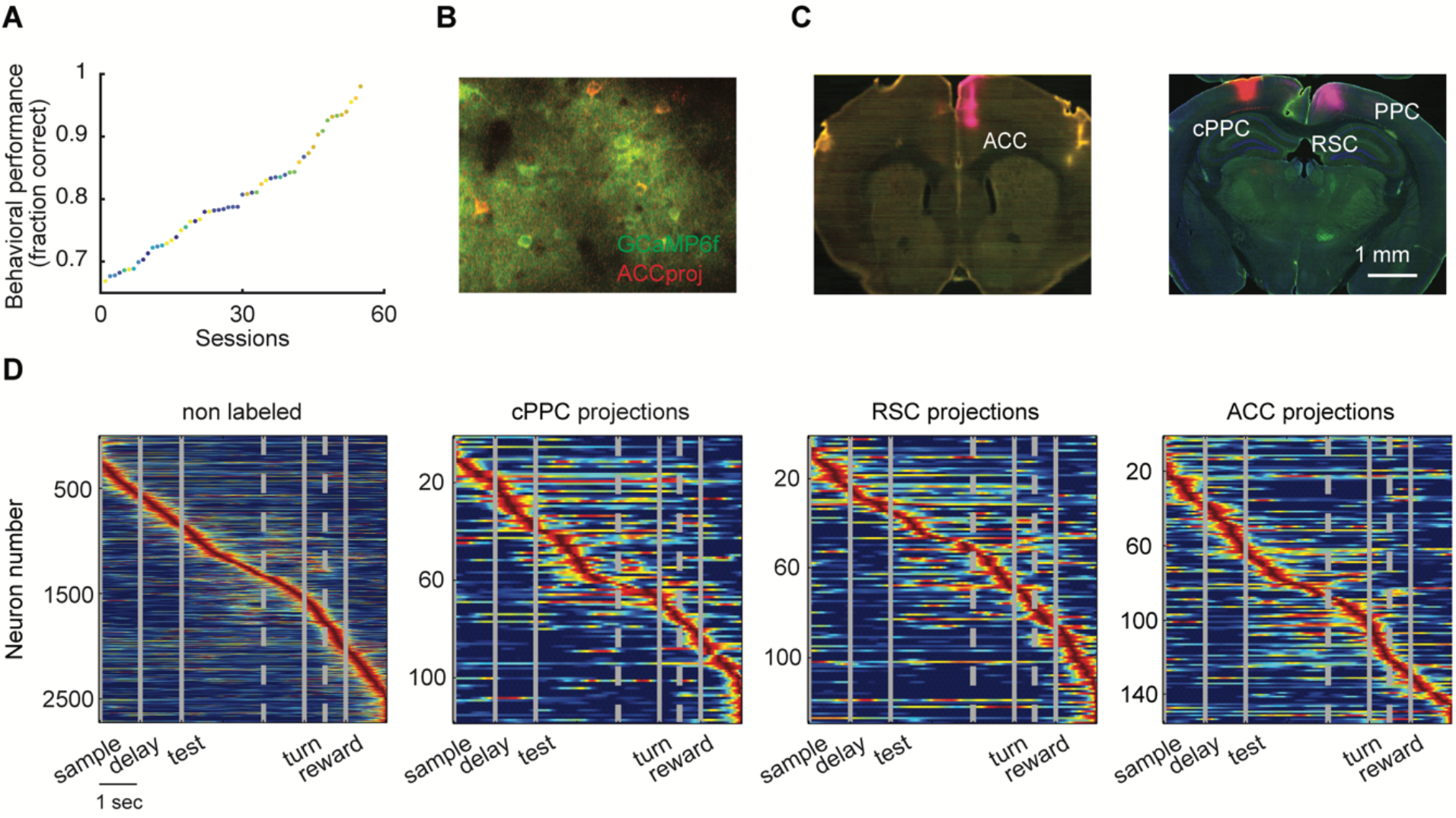
Retrograde labelling and PPC activity sequences. **(A)** Sorted behavioral performance in all sessions. Each color corresponds to one mouse. (**B**) Example two-photon imaging field of view. (**C**) Injection sites for 600 retrograde tracers in three different target areas to label PPC projection cells. (**D**) Similar to Figure 1E, normalized 601 deconvolved calcium activity of PPC neurons sorted based on the cross-validated peak time, for populations of 602 non-labeled cells and cells projecting to contralateral PPC, RSC, or ACC. Vertical gray lines represent onsets of 603 sample cue, delay, test cue, turn to arms, and reward. Gray dashed lines correspond to one second before turn 604 and 0.5 second before reward.

**Figure S2.**
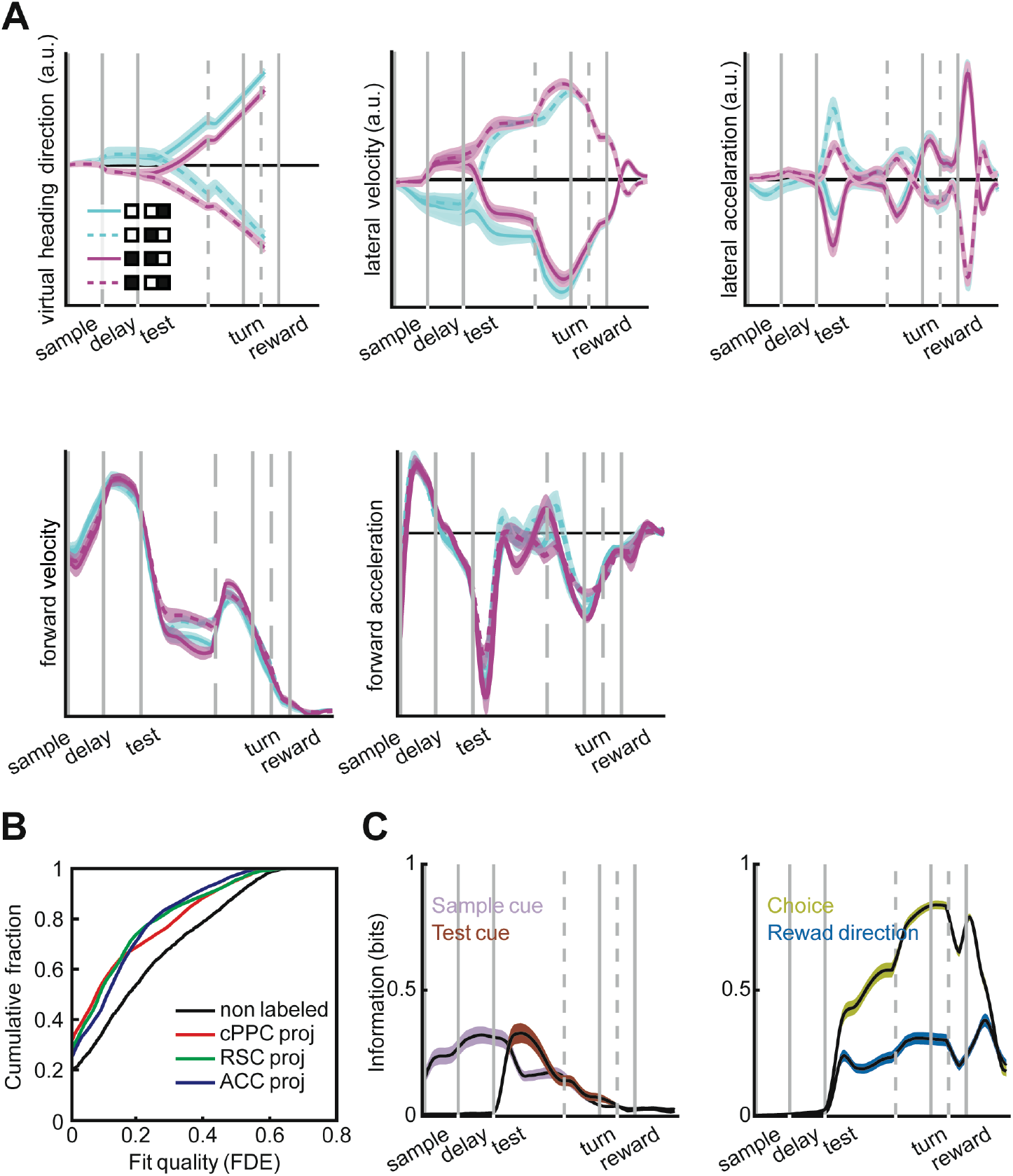
NPvC goodness of fit for different cell types. (**A**) Distribution of virtual heading direction, lateral velocity, lateral acceleration, forward velocity, and acceleration during the trial for different trial types. (**B**) The cumulative distribution of goodness of fit (fraction of deviance explained) for NPvC model for different populations. (**C**) Mutual Information between task variables (sample cue, test cue, choice, and reward direction (xor)) and behavioral variables (virtual heading direction, lateral velocity and acceleration, and forward velocity and acceleration). In Panels A,C, lines and shaded areas plot mean and SEM over experimental sessions.

**Figure S3.**
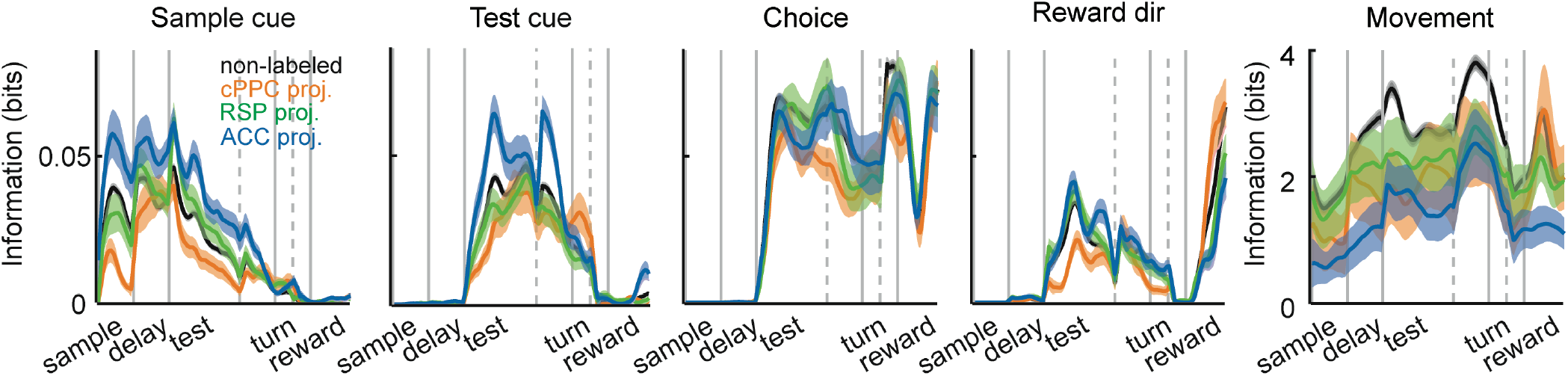
Single-neuron information in PPC neurons. Time course of different information components in PPC 616 neurons, similar to Figure 3A, divided into non-labeled neurons and neurons projecting to cPPC, RSC, or ACC. 617 Lines and shaded areas plot mean and SEM over neurons.

**Figure S4.**
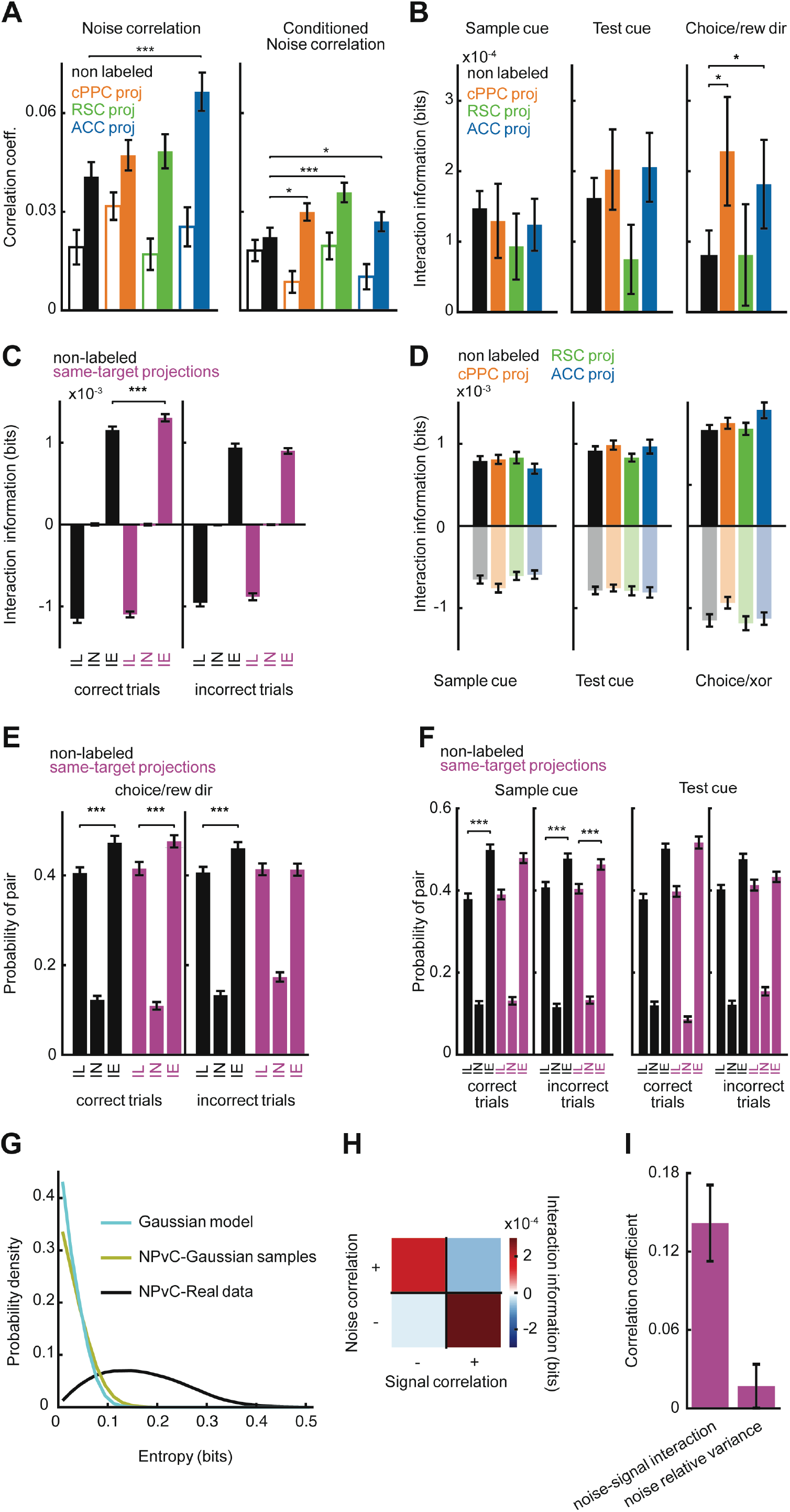
Pairwise noise correlation and interaction information for distinct projection targets. (**A**) Noise correlations (left) and conditional noise correlations (right) computed for pairs of neurons in correct (filled bars) and incorrect trials (open bars). Similar to Figure 4B, for different projection targets. (**B**) Mean interaction information of non-labeled and same-target projection pairs for sample cue, test cue, and choice/reward direction in correct trials. (**C**) Average interaction information for information-enhancing (IE), information-limiting (IL), and independent (IN) pairs. In all panels, means are from all neurons, and error bars are the standard error of mean estimated by bootstrapping. (**D**) Mean interaction information for information-enhancing (opaque bars) and information-limiting pairs (transparent bars) for sample cue, test cue, and choice/reward direction in correct trials. (**E**) The probabilities of information-enhancing (IE), information-limiting (IL) and independent (IN) pairs of neurons for choice/reward direction using correct trials. (**F**) The probabilities of information-enhancing pairs (IE), information-limiting pairs (IL), and independent pairs (IN) for non-labeled and same-target projection neurons for sample cue and test cue information using correct trials. (**G**) The distributions of entropy extracted from the correlation structures between pairs of neurons, computed either using the NPvC model (black), the analytic formula for a Gaussian distribution with linear correlation computed from each pair (cyan), or from samples generated from the Gaussian distribution with the same correlation value but computed using the nonparametric copula (green) for control. The result shows a significant positive shift in the value of the entropy extracted using the nonparametric copula, compared to a Gaussian distribution capturing only the linear correlation structure of the data. This result suggests that there are neural correlations that have significant non-linearity. Thus, using the copula can help to capture these correlations between pairs of neurons beyond just the linear component. This suggests that by using the copula to quantify the correlations and interactions between neurons, we potentially include part of entropy of pairwise interaction that would have been ignored if we used a simple Gaussian quantification for the pairwise correlation. This component of the entropy can potentially contribute to the interaction information for a neural pair. This result suggests the importance of considering correlation components beyond the Gaussian linear correlation. (**H**) The mean interaction information for each combination of signal correlation and noise correlation signs for neurons projecting to the same target during correct trials. The four quadrants correspond to the combinations of positively signed (+) and negatively signed (-) noise and signal correlations. (**I**) The correlation coefficient between the pairwise interaction information and two features of noise correlations. The first feature is the signal-noise correlation similarity coefficient (left), defined as *J* = −*nc*. sgn(*sc*) (where *nc* is the noise correlation and sgn(*sc*) is the sign of the signal correlation which is +1 for neurons with similar choice selectivity and -1 for pairs with different choice selectivity). This coefficient captures how well aligned signal and noise correlations are. The second feature is the normalized variance across stimuli of the noise correlation (right) defined as 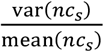 where *nc*_*s*_ is the noise correlation for stimulus *s*. This coefficient captures the strength of the stimulus-dependence of noise correlations. Both signal-noise similarity and stimulus dependence of noise correlations are factors that can create information-enhancing correlations. These results show that in our PPC dataset, the signal-noise similarity is the main factor creating information-enhancing correlations. In all the panels, * indicates p < 0.05, ** indicates p < 0.01, *** indicates p < 0.001, t-test with Holm-Bonferroni correction. In Panels A-F,I, dots and errorbars represent mean and SEM over all pairs of neurons.

**Figure S5.**
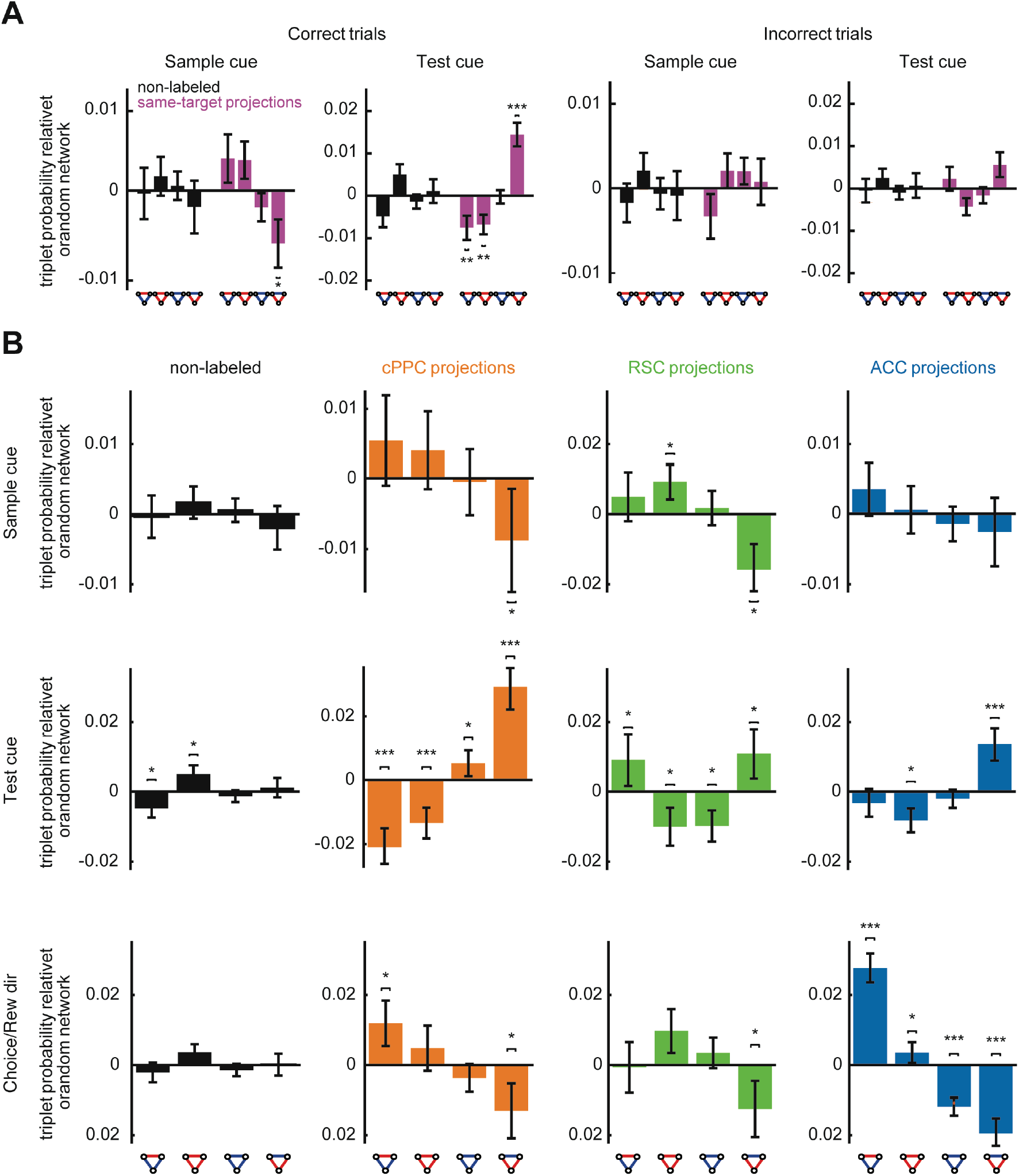
Expanded quantification of triplet probabilities. (**A**) Relative triplet probability with respect to a random network (obtained by shuffling the organization of pairs) for non-labeled and same-target neurons during correct (left two panels) and incorrect (right two panels) trials for sample cue and test cue information. (**B**) Similar to (**A**) and Figure 5C for neurons projecting to different targets. In all panels, * indicates p < 0.05, ** indicates p < 0.01, *** indicates p < 0.001, t-test with Holm-Bonferroni correction for statistical multiple comparisons. Dots and errorbars plot mean and SEM is computed by bootstrapping over all triplets of neurons.

**Figure S6.**
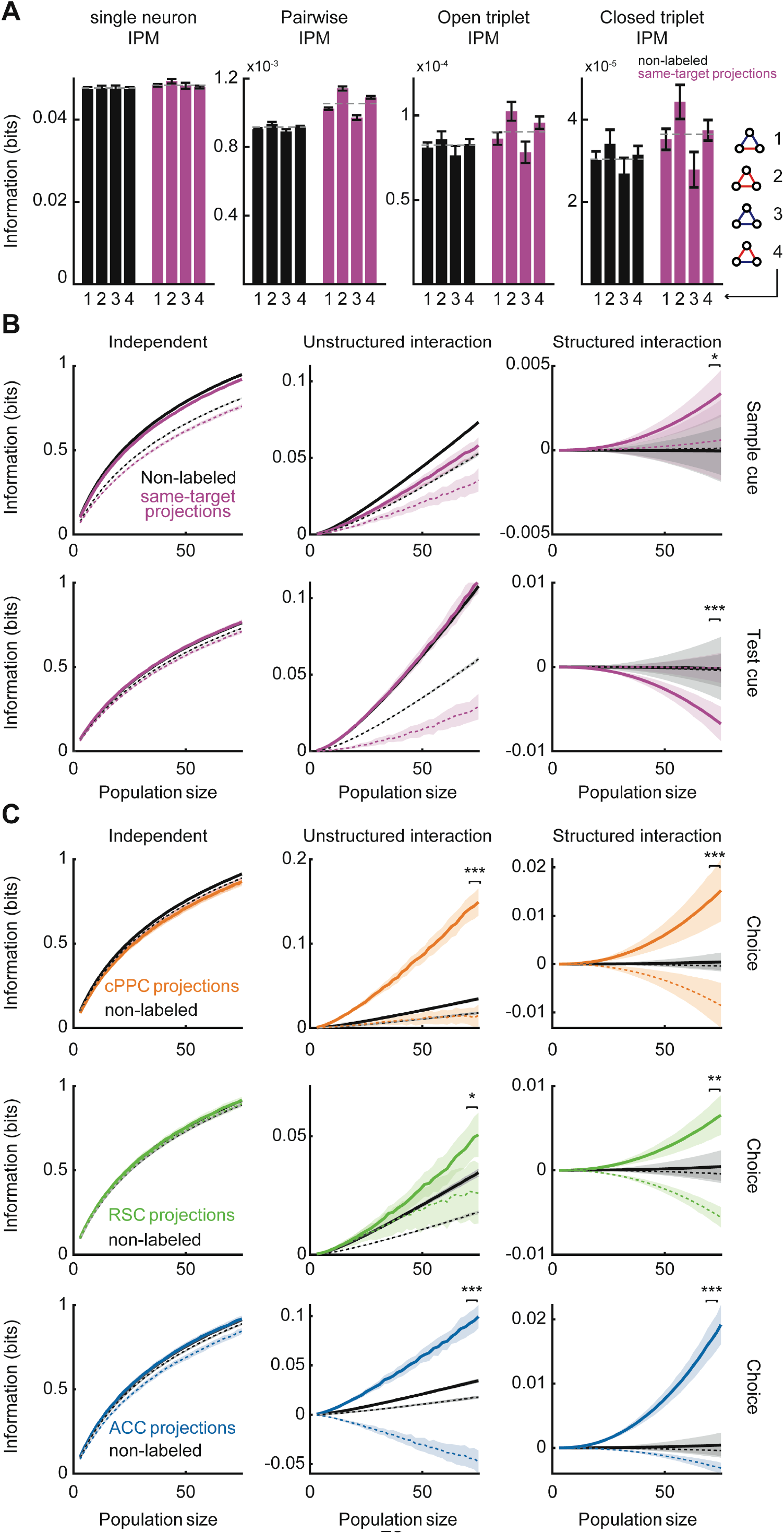
Structured interaction information for different task variables and projection targets. (**A**) The mean single-neuron information per motif, pairwise interaction information per motif, and information per motif from open and closed triplets computed for each triplet type using the non-labeled and projecting to the same target PPC neurons. (**B**) Independent, unstructured interaction, and structured interaction information for non-labeled and same-target projection PPC neurons during correct and incorrect trials for increasing population size and for sample cue and test cue information. (**C**) Independent, unstructured interaction, and structured interaction information for non-labeled and same-target projection neurons during correct and incorrect trials for increasing population size, similar to Figure 6E. In Panel A, dots and errorbars plot mean and SEM across triplets. In Panels B,C, Lines (filled for correct trials, dashed for incorrect trials) and shades represent Mean and SEM estimated from bootstrapping over triplets. In all panels, * indicates p < 0.05, ** indicates p < 0.01, *** indicates p < 0.001, t-test with Holm-Bonferroni correction for statistical multiple comparisons.

**Figure S7.**
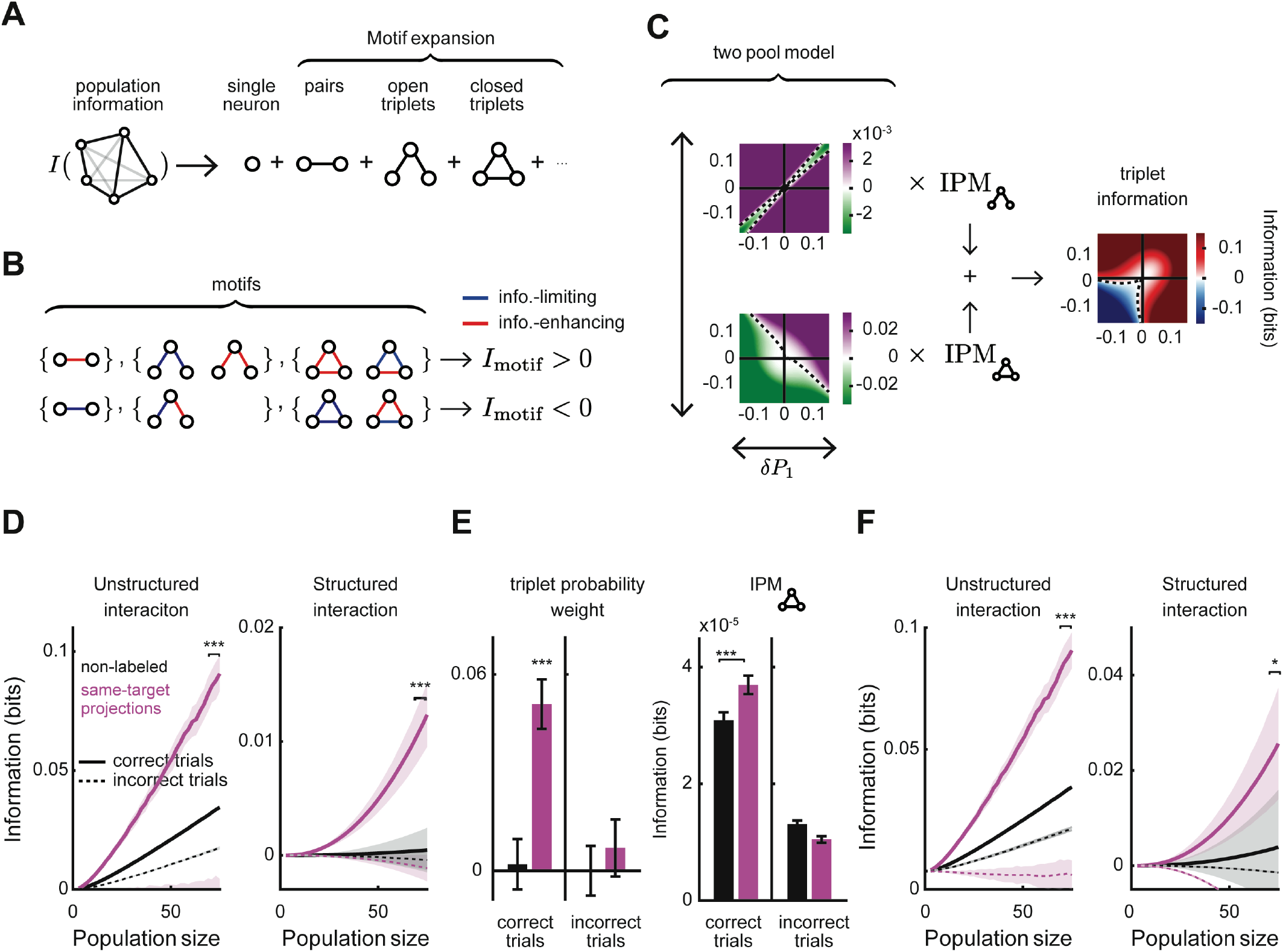
Structured interaction information for non-symmetric two-pool network. **(A,B)** Schematics representing how for a non-symmetric network the population information can be expanded as a sum of contributions from network motifs with increasing complexity similar to the case of the symmetric networks shown in Figures 6B,C. In this case, more motifs contribute, such as open triplets. **(C)** In the case in which IPMs are equal for different motif types in a two-pool network, structured interaction information is proportional to the motif probability weights computed for open and closed triplets and the IPM (which is a positive definite number). In the two left plots, the colormap represent the analytic value of the triplet probability weight for open (top) and closed (bottom) triplet motifs for any two-pool network after using the results presented in Supplemental Information: Simplifying the structured interaction information. Depending on the structure of the two-pool network, the structured interaction information can be either information-enhancing (corresponding to areas with purple colors) or information-limiting (corresponding to areas with green colors). The structured interaction information will be a weighted sum of the open and closed triplet terms weighted by their counts and their IPM’s. A simulated example of structured interaction information over the two-pool network space computed using n=75 neurons and the IPM’s IPM_∧_ = 0.001 and IPM_△_ = 0.0001 is shown in the right-hand panel. All panels are computed using the analytic formula presented in Eq. SM44, SM45, and SM47 for a range of two-pool networks. **(D)** Independent, unstructured interaction, and structured interaction information computed for non-labeled (black) and same-target projection (pink) neurons during correct and incorrect trials for increasing population size. Similar to Figure 6E but without assuming that the network is symmetric. The statistical comparisons were made for values corresponding to n = 75 population size. (**E**) The triplet probability weight (left) and interaction information per triplet (right) for non-labeled and same-target projection neurons during correct and incorrect trials. The statistical comparison in the left panel is to check whether the values are different from zero and in the right panel is to test differences between non-labeled and same-target projection distributions. (**F**) Similar to (**D**) but assuming that the interaction per motif (IPM) weights are the same for all the triplet types and using Eq. SM44, SM45, and SM47 for a symmetric network. The statistical comparisons were made for values corresponding to n = 75 population size. In all panels * indicates p < 0.05, *** indicates p < 0.001, t-test with Holm-Bonferroni correction. In all panels mean and SEM computed by bootstrapping over all triplets of neurons are shown.

**Figure S8.**
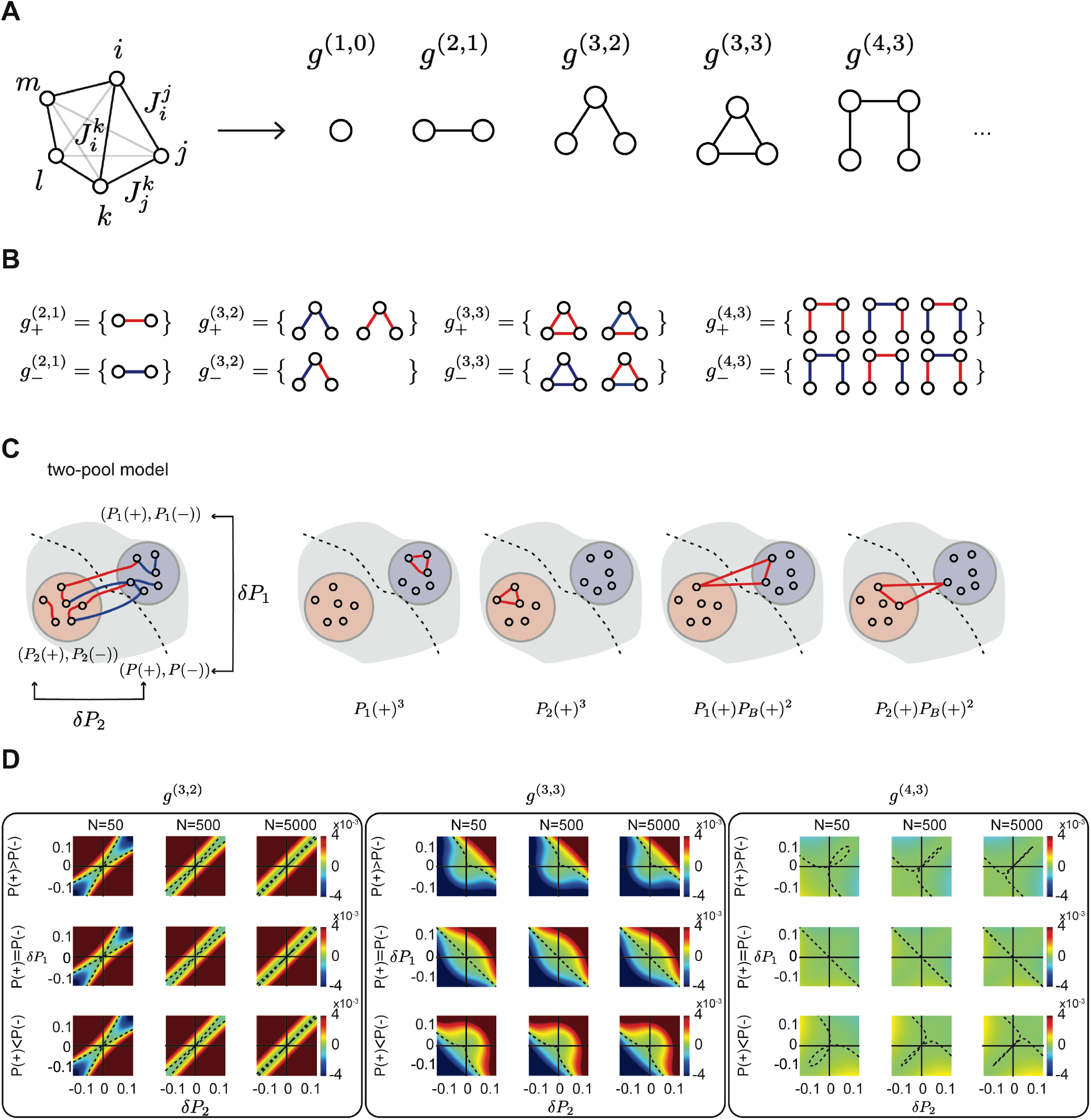
Analytical expansion of the population information. **(A)** Breakdown of interaction information in individual graph motifs. **(B)** Subgraphs with positive and negative contribution to the information for each motif. Red and blue edges in each graph correspond to information-enhancing and information-limiting pairwise interactions, respectively. **(C)** Left: In the two-pool model, the network consists of two different pools of interactions each of which have a different distribution of information-enhancing and information-limiting. Right: Three possible ways of building a ‘+,+,+’ triplet, with their probabilities are shown. **(D)** Probability weights presented in Eq. SM47 for open triplets (left box), closed triplets (middle box), and quadruplets (right box) for different network sizes (each column is one network size) and different δ*P* = *P*(+) − *P*(−) (upper row: *P*(+) − *P*(−) = 0.1, middle row: *P*(+) − *P*(−) = 0, lower row: *P*(+) − *P*(−) = −0.1).

**Figure S9.**
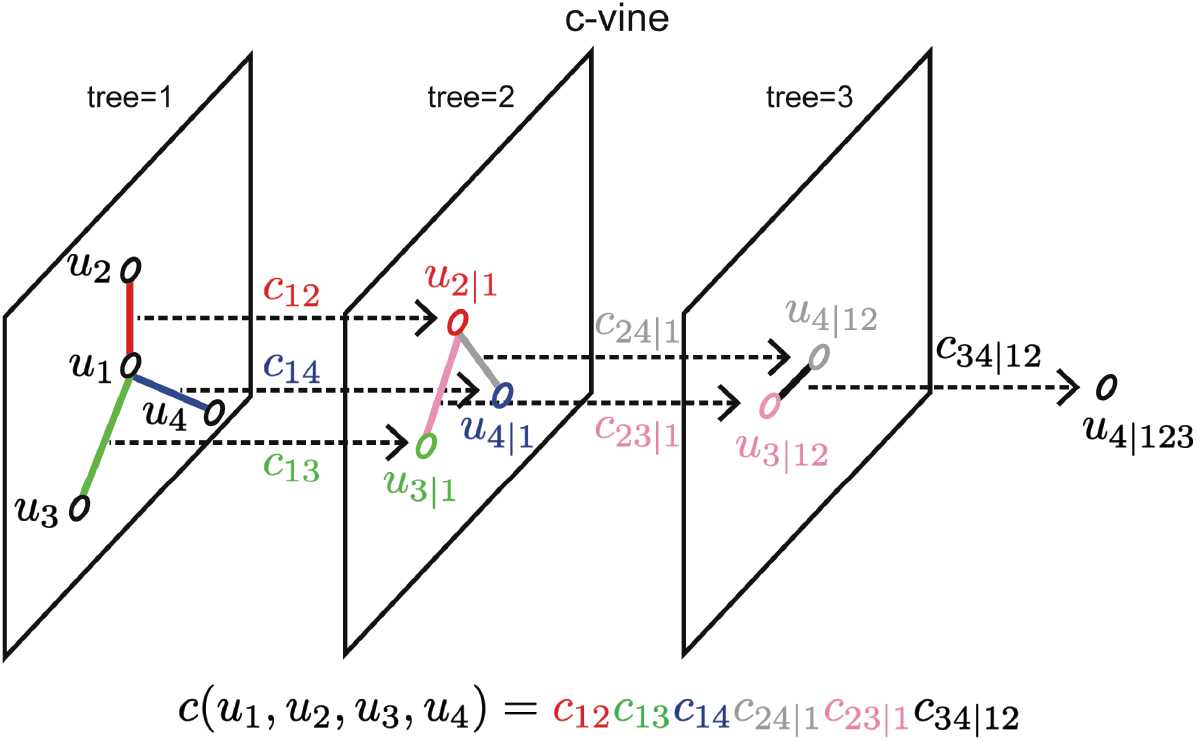
Vine copula graphical model. Schematic of the c-vine copula sequential computation of four-dimensional data. The multivariate copula density function is computed from the product of a series of sequential bivariate copulas which are being computed as it is explained in Supplemental Information: Vine copula modelling of neural responses.

## Supplemental table

**Table S1.** Interaction weight per motif for single network subgraphs represented in terms of single neuron 729 information and pairwise interaction information values of the subgraph.

| Motifs | $g^{(2,1)}$ | $g^{(3,2)}$ | $g^{(3,3)}$ | $g^{(4,3)}$ |
| --- | --- | --- | --- | --- |
| $W_{g^{(n,e)}}$ | Eq.SM26 | $e^{-2I^{\text{ind}}} \frac{I_{ik}I_{kj}}{I_k}$ | $e^{-2I^{\text{ind}}} \frac{I_{ik}I_{kj}I_{ji}}{I_iI_k}$ | $\frac{e^{-2I^{\text{ind}}}}{I_kI_l} I_{ik}I_{kl}I_{lj}$<br>$i \neq j$ |

## Methods

### Mice

All experimental procedures were approved by the Harvard Medical School Institutional Animal Care and Use Committee and were performed in compliance with the Guide for Animal Care and Use of Laboratory Animals. Imaging data were collected from 10 male C57BL/6J mice that were eight weeks old at the initiation of behavior task training (stock no. 000664, Jackson Labs).

### Virtual reality system

The virtual reality system has been described previously^25^. Head-restrained mice ran on an 8-inch diameter spherical treadmill. A PicoP micro-projector was used to back-project the virtual world onto a half-cylindrical screen with a diameter of 24 inches. Forward/backward translation was controlled by treadmill changes in pitch (relative to the mouse’s body), and rotation in the virtual environment (virtual heading direction) was controlled by the treadmill roll (relative to the mouse’s body). Movements of the treadmill were detected by an optical sensor positioned beneath the air-supported treadmill. Mazes were constructed using VirMEn in MATLAB^68^.

### Behavioral training

Prior to behavioral training, surgery was performed in 8-week-old mice to attach a titanium headplate to the skull using dental cement. At least one day after implantation, mice began a water schedule, receiving at least 1 ml of water per day. Body weights were monitored daily to ensure they were greater than 80% of the original weight. Mice were trained to perform the delayed match-to-sample task using a series of progressively complex mazes. Mice were rewarded with 4 μl of water on correct trials. First, naive mice were trained to run down a straight virtual corridor of increasing length for water rewards. In the second stage, mice learned to run in a T-shaped maze, into either a left or right choice arm. The correct choice arm was signaled by the presence of a tall tower at the choice arm. After the mouse was able to run straight on the ball and make turns, we trained mice on stage 3, where we began to familiarize the mice with running into the choice arm that matches the color presented in the maze stem (sample cue). The walls of the stem were either black or white (sample cue; randomly selected from trial to trial). The left and right choice T-arms were black and white, respectively, or vice versa (test cue). At this stage of training, the correct choice arm was signaled both by a tall tower in the correct arm and by the T-arm that matched the color of the sample cue. Note that in this stage, the mouse can still perform accurately even if it ignores the sample and test cues, as it can just run to the choice arm that has the tower. In the fourth stage, the maze was the same, except that we added a tower to the unrewarded choice arm as well. In this stage, the mouse cannot simply run to whichever arm has a tower (since both arms have towers) and must run into the arm that matches the color of the sample cue. In stage four, the maze was exactly like the previous one, except the choice arms and towers (test cue) appeared grey as the mouse ran down the maze. The colors of the choice arms and towers (test cue) were not revealed until the mouse reached 3/4 of the way down the stem. When the black and white choice arms were revealed, the mouse can begin to plan and execute a turn left or right. In the final stage, we begin training towards the final implementation of the delayed match-to-sample task. We introduced a delay segment by making the walls of the T-stem grey at the end of the stem and gradually increased the length of the stem that is grey (delay segment). The mouse received its first trials of the delayed match-to-sample task when at least the entire last 1/4 of the stem walls were grey. In such trials, when the mouse’s position reaches 3/4 of the way down the stem, there was a short moment in which both the stem walls and the choice arms were grey. At this point, the mouse must rely on its memory of the initial sample cue walls to make the correct turn. We made this grey segment longer and longer until the mouse performed with over 85% accuracy for a delay averaging at least two seconds in duration. The entire training program was completed in 12-18 weeks.

### Surgery

When mice reliably performed the delayed match-to-sample task, the cranial window implant surgery was performed. Mice were given free access to water for two days prior to surgery. During the surgery, mice were anesthetized with 1.5% isoflurane and the headplate was removed. Craniotomies were performed over PPC centered at 2 mm posterior and 1.75 mm lateral to bregma. GCaMP6 was injected at 3 locations spaced 200 μm apart at the center of the PPC. A micromanipulator (Sutter, MP285) moved a glass pipette to approximately 250 μm below the dura, and a volume of approximately 50 nL was pressure-injected over 5-10 minutes. Dental cemented sealed a glass coverslip on the craniotomy, and a new head plate was implanted, along with a ring, to interface with a black rubber objective collar to block light from the VR system during imaging. Craniotomies were also performed over the contralateral PPC (2 mm posterior and 1.75 mm lateral to bregma), anterior cingulate cortex (1.34 mm anterior and 0.38 mm lateral to bregma, and 1.3 mm ventral to the surface of the dura). Over 200 nL of retrograde tracer (CTB-Alexa647, CTB-Alexa405, or red retrobeads) was injected at each site. The retrograde tracer injection into RSC was made through the craniotomy for PPC imaging, targeting the most medial portion of the craniotomy (∼0.5 mm lateral from the midline). Craniotomies for the retrograde tracers were sealed with KwikSil (World Precision Instruments). Mice recovered for 2-3 days after surgery before resuming the water schedule. Imaging was performed nearly daily in each mouse, starting 2 weeks after surgery, and continued for 1-2 weeks.

### Obtaining anatomical stacks

Anatomical stacks of cells double-labeled with GCaMP and retrograde tracers were acquired using a two-photon microscope taken every 2 μm from the surface of the dura to about 300 μm below the surface. For images of cells double-labeled with GCaMP and CTB-Alexa647, we used a dichroic mirror (562 long pass, Semrock) and bandpass filters (525/50 and 675/67 nm, Semrock) and delivered the excitation light at 820 nm to visualize both GCaMP and the Alexa fluorophore simultaneously. We also took images of GCaMP and CTB-Alexa405 using a dichroic mirror (484 long pass) and bandpass filters (525/50 and 435/40) and delivered excitation light at 800 nm. Finally, we took images of GCaMP and red retrobeads with dichroic mirror (562 long pass) and bandpass filters (525/50 and 609/57) and delivered excitation light at 820 nm. These anatomical stacks allowed us to identify cells later visually as either double-labeled with GCaMP and retrograde tracer or GCaMP-expressing and unlabeled with retrograde tracer. We then matched these GCaMP-expressing neurons from the anatomical stacks with the same GCaMP-expressing neurons during functional imaging, which was performed at 920 nm and during behavior. In some sessions, we took z-stacks immediately before functional imaging to ensure some labeled cells were present in a field of view.

### GCaMP imaging during behavior

For functional imaging, 5 mice were imaged using a Sutter MOM at 15.6 Hz at 256 x 64-pixel resolution (∼250 x 100 um) through a 40x magnification water immersion lens (Olympus, NA 0.8). Five other mice were imaged using a custom-built two-photon microscope with a resonant scanning mirror (at 30 Hz frame rate) with a 16x objective. ScanImage was used to control the microscope. Imaging sessions lasted 45-60 minutes. Each session was imaged over multiple 10-minute acquisitions separated by 1 minute, allowing any amount of GCaMP bleaching to recover. The imaging frame clock and an iteration counter in VirMEn were recorded to synchronize imaging and behavioral data.

### Data processing

Custom-written MATLAB software was designed for motion correction, definition of putative cell bodies, and extraction of fluorescence traces (dF/F). Fluorescence traces were deconvolved to estimate the relative spike rate in each imaging frame and all analyses were performed on the estimated relative spike rate to reduce effects of GCaMP signal decay kinetics.

To estimate the neural activity of individual cells from the calcium imaging data, we processed the data by the following steps. (1) Motion correction: Motion artifacts in imaging data were corrected in each imaging frame. First, “line-shift correction” was performed to align for line-by-line alternating offsets in images due to bidirectional scanning. Then, “sample movement correction” was performed to remove between-frame rigid movement artifacts by FFT-based 2d cross-correlation ^69^ and within-frame non-rigid movement artefacts by Lucas-Kanade method ^70^. (2) Cell selection: The spatial footprint of a putative cell was identified based on the correlation of fluorescent time series between nearby pixels. The correlation of fluorescence time series was calculated for each pair of pixels within a ∼60 x 60 μm square neighborhood. Then putative cells were identified by applying a continuous-valued, eigenvector-based approximation of the normalized cuts objective to the correlation matrix, followed by discrete segmentation with k-means clustering, which generated binary masks for all putative cells. (3) dF/F calculation: The magnitude of calcium transients was estimated by subtracting the background fluorescence from the raw fluorescence of a putative cell. For each putative cell, the background fluorescence of local neuropil was estimated by the average fluorescence of pixels that did not contain putative cells. Then neuropil fluorescence time series was scaled to fit the raw fluorescence of a putative cell by iteratively reweighted least-squares (*robustfit*.*m* in MATLAB) and was subtracted from the raw fluorescence to yield neuropil-subtracted fluorescence (Fsub). Then, dF/F was calculated as (*F*_sub_–*F*_basline._)/*F*_basline_, where *F*_baseline_ was a linear fit of *F*_sub_ using iteratively reweighted least-squares (robustfit.m in MATLAB). The codes used in steps (1-3) are available online: (https://github.com/HarveyLab/Acquisition2P_class.git). (4) Deconvolution: The timing of spike events that led to calcium transients was estimated by deconvolution of fluorescent transients. dF/F was deconvolved by OASIS AR1 ^71^, which models the fluorescence of each calcium transient as a spike increase followed by an exponential decay, whose decay constant was fitted to each cell. The deconvolved fluorescence resulted in spikes that were sparse in time and varied in magnitude. The deconvolved fluorescence was used as neural activity for the majority of the analyses.

### Non-parametric vine copulas (NPvC) models of single neuron activity

To quantify the information carried by a neuron’s activity about task variables, while discounting possible contributions from movement variables, we built a multivariate probabilistic model of the activity of each neuron, time, behavioral variables (running velocity and acceleration), and task variables (schematized in Figure 2C). We built this model using non-parametric vine copulas (NPvC). We chose vine copulas because they allow constructing arbitrarily complex multivariate relationships (including relationships between non-neural variables, e.g. relationships between trial types and behavioral variables) by combining bivariate relationships, which can be sampled accurately and robustly from finite number of trials regardless of the details of the marginal distributions. We chose non-parametric estimators because the form of these relationships is not known *a priori* and we wanted to avoid biases originating from inaccurate assumptions. Here, we describe the details of the computation of our non-parametric vine-copula model, specifically for computing the probability density function of neural population activity given behavioral and task variables at each time point in the trial (Figure 2D) and for computing its goodness of fit (Figure 2E).

We used vine copulas to estimate the conditional joint probability density function 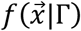 for a set of variables 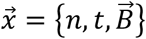, which in our case consists of the activity of a neuron *x*_1_ ≡ *n*, time *x*_2_ ≡ *t* and the 5-dimensional vector 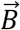 of the five behavioral variables that we measured (virtual heading direction, lateral and forward running velocities, and lateral and forward accelerations), for each trial type Γ. Each trial type Γ (Γ = 1 … 8) is defined by the sample cue, the test cue, and trial outcome (correct or incorrect). Thus, there are four trial types for trials with correct choices and four trial types for incorrect trials. Using the copula decomposition, the probability density function for each trial type Γ is represented as a product of the single variable marginal probability density functions 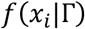, and the copula 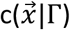, which captures the dependencies between all the variables, as follows:

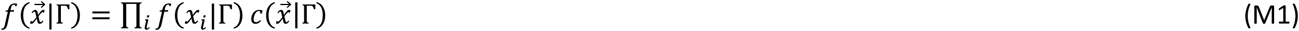

We used a kernel density estimator ^72^ to compute the single-variable marginal probability densities and a non-parametric c-vine vine copula to estimate the copula, representing the correlation structure between variables. We used a c-vine graphical model with neuron activity as the central variable in the model. The probability density function of a c-vine can be expressed as the product of a sequence of non-parametric bivariate copulas ^31,73^ (Supplemental Information: Vine copula modelling of neural responses). We used similar order for the variables in the c-vine graphical model as 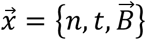 introduced above. Furthermore, we simplified the vine copula structure by considering the decomposition of the copula as a product of a time-dependent and a time-independent component 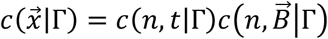 meaning that we considered that the tuning of neurons to movement variables is time-independent. The sequence of bivariate copulas shaping the vine-copula were then fitted to the data using a sequential kernel-based local likelihood process (Supplemental Information: Nonparametric pairwise copula estimation)^34^. For each of the bivariate copulas, the kernel bandwidth was fitted so as to maximize the local likelihood obtained in a 5-fold cross-validation method ^34^. Using the estimated bandwidths for each copula in the vine sequence, we computed the multivariate copula density function of data points using a 5-fold cross-validation process. We first used the training set to estimate the copula density on a 50 by 50 grid (Supplemental Information: Nonparametric pairwise copula estimation) and then used the copula estimated on the grid point to interpolate the copula density on the test set. A similar procedure was followed to estimate cross-validated single variable marginal density functions. The conditional probability of neural responses in each trial condition and for each value of behavioral variables is computed using 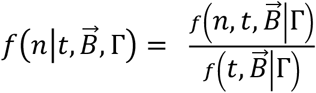 where to compute the density function 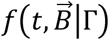, we marginalized by integrating over the neuron response

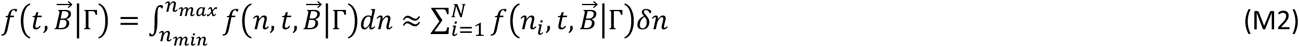

In Eq M2, we approximated the integral as a sum over a set of *N* = 200 points {*n*_*i*_} ranging linearly between the minimum (*n*_*min*_) and maximum (*n*_*max*_) value of the neural response within the session and 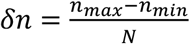. By computing the density function 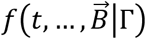 by marginalization of 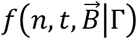, instead of fitting a new vine model between 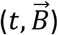 variables, any difference between these two density functions is only related to the dependency of the neural activity to other variables and not to a difference between how dependencies between 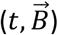 variables are being quantified in the two models because of fitting differences.

For each time point, we computed a copula fit of the activity of each neuron (indicated as NPvC fit in Figure 2D) as a point *n*_*cop*_ ∈ {*n*_*i*_} with largest log likelihood given the considered trial type and the values of behavioral variables at the considered time point:

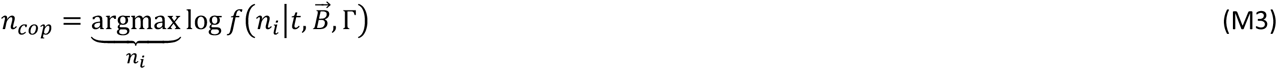

### Generalized Linear Models (GLMs) of single neuron activity

We also fit a GLM model to compare it with the copula. We used a GLM with Poisson noise, a logarithmic link function, and an elastic-net regularization^23,26,35^. We used task variables defining the trial type consisting of sample cue, test cue, choice, and all their interactions (each binary task variable was coded as -1 or +1) together with the same behavioral variables we used in copula modelling as the predictors. Time-dependency during the task for single neurons was expanded using a raised cosine basis ^26^ as follows:

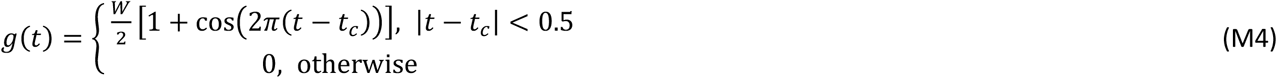

For task variables, the cosine basis function has a width of 1 s, and the value at the center peak was either positive (*W* = +1) or negative (*W* = −1) depending on the identity of each task variable. The center peaks *t*_*c*_ were spaced with a 0.5 s interval to tile the epoch with a half-width overlap. For selectivity to movements of the mouse, we first z-scored each movement variable as *Z*_*i*_ and used similar cosine basis as follows:

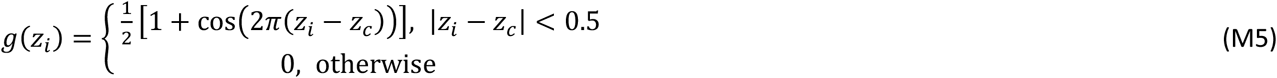

We considered 13 center peaks *Z*_*c*_ ranging from -3 to 3 with spacing of 0.5 for the cosine basis of movement variables.

### Computation of neural data fitting performance of NPvC and GLMs

The performance of both the NPvC and the GLM in fitting single-trial single-neuron activity was evaluated by computing the fraction of the deviance explained (FDE) on the test data, defined as follows ^74^:

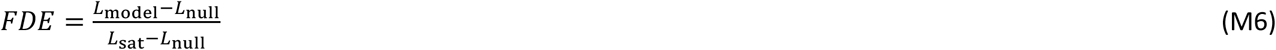

where *L*_model_, *L*_null_, and *L*_sat_ are the likelihoods of observing the test data for the considered model (copula or GLM), the null model, or saturated model, respectively and are always computed for all the time points in all trials in a cross-validated fashion. For the GLM, the null model is a model that does not have any predictors, and its prediction of activity at any time point is the time-averaged rate of the neuron. For the copula null model, we defined a model of neural response without including any of the predictors in the trial, which quantifies how much of the neural activity can be explained without knowing the time and movement variables used in the vine copula model. The likelihood is then computed using the marginal distribution of neural activity *L*_null_ = log [*f*(*n*_data_)]. The saturated model is the generative model in which the prediction exactly matches the observed activity at each time point in the test data. For the GLM, these values were computed using the analytical form of the GLM output at each time point using the logarithmic link function. For the copula model, which is nonparametric, the vine-copula model is used to compute the likelihood for each neural activity 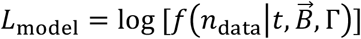 as the model likelihood where *n*_data_ is the real single trial neural activity at time t from data. To compute the saturated likelihood, we considered the fact that the model prediction is derived from Eq. M3 and the saturated likelihood was considered to be the peak likelihood obtained 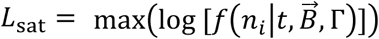 where {*n*_*i*_} is the set of *N* = 200 grid points on the neural activity described before ranging from the minimum to maximum possible neural activity of each neuron. Each of the likelihood values on the data were computed using a 5-fold cross-validation. The density function is estimated over a grid using the training set, and the grid is used to compute the density function over the test set (Supplementary Method 2). We considered five different task epochs and fitted the model to each of them. The periods were from 0.5 s before to 2 s after the sample cue onset, delay onset, test cue onset, start of the turn, and reward onset. We used only sessions that had at least 5 trials of each of the 8 trial types.

### Estimation of Mutual information for single neurons

We used probabilities estimated from the vine-copula approach, explained in the previous section, to estimate single neuron mutual information values (presented in Figure 3). To compute the mutual information between a group of variables (including neural activity and/or behavioral variables) and a task variable, we used a decoding approach and then computed information in the confusion matrix^75–77^. A task variable (*c*) here was a binary variable taking a value *c* of +1 or -1 to indicate one of the two possible values of either sample cue, test cue, reward location, or choice direction. For each task variable, we decoded its value in each trial using the copula model to compute (through Bayes’ rule) the posterior probability of the task variable given the observation in the same trial of a set of variables 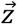, which could be the full set 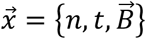. or a subset of it. This posterior probability of *c* can be obtained from the copula model as follows:

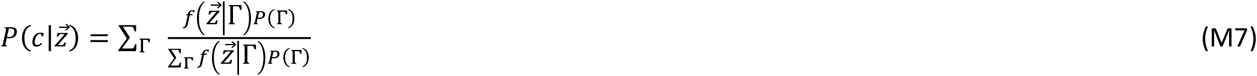

where the sum is over all the trial types Γ (corresponding to the combinations of two sample cues, two test cues, and two trial outcomes (correct or incorrect) and consists of eight possible values when using all trials and four values when using either correct or incorrect trials) which have value *c* for the considered task variable. For example, considering analyses limited to correct or incorrect trials the sum will be over the 4 correct or incorrect trial types. Considering analyses using all the trials the sum will be over all 8 trial types. Considering sample cue correct trials only, the sum in Eq. M6 will be over the two trial types with correct outcome and two sample cues, and so on. *P*(Γ) is the probability of occurrence of trial type Γ across all considered trials. In the above equation, the probability density of the variable 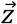 given the trial type Γ,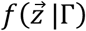, is computed using the copula model by using Eq. M1 when 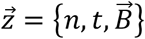 or using Eq. M2 when 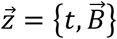.

We then used the posterior probabilities to decode the most likely task variable given the variables 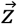 observed in the considered trial:

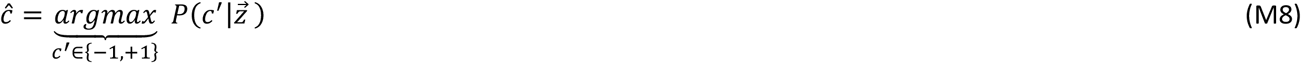

The information about the task variable decoded from neural activity was then computed as the mutual information between the real value *C* of the variable and the one *Ĉ* decoded from neural activity, as follows:

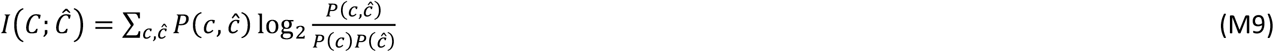

where *P*(*C,Ĉ*) is the confusion matrix, that is the probability that the true value of the task variable is *c* and the value of the decoded one is *Ĉ*, and *P*(*c*) and *P*(*Ĉ*) are the marginal probabilities.

To compute the information 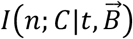 between the task variable and the neural activity at a given time point, conditioned over the behavioral variables, we used the following equation for conditional mutual information:

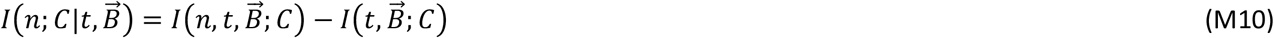

In the above equation,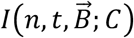 was computed after decoding *Ĉ* using the copula derived from Eq. M1.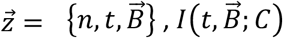 was computed after decoding *Ĉ* using the copula constricted with Eq. M2 a 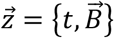.

We computed the mutual information 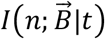 between the neural activity and the behavioral variables 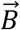 at time *t* (used in Figure 3A-B) using Shannon’s mutual information formula with probabilities defined by the copula 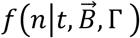, the marginalized copula *f*(*n*|*t*, Γ ), and the probability *P*(Γ) of each trial type Γ as follows:

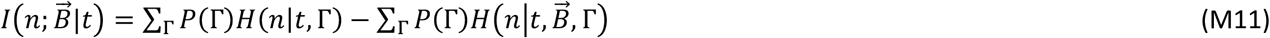

The entropies 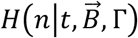 and *H*(*n*|*t*, Γ) were computed by averaging the corresponding NPvC estimated log-likelihoods at the neural activity and movement values sampled from the trial type Γ at each time point.

For the information about the task variables, we chose a computation of information from the decoding matrix, similar to e.g.^23^, because when computing neural population information about categorical values this computation is robust and has a lower variance and bias than a direct computation from the probabilities. The reason is because the decoding step compresses the dimensionality of the neural responses from which information is computed, especially when considering neuron pair responses and interaction information. This computational robustness was useful because we wanted to use the information values for each neuron pair about task variables at the single neuron and the single pair levels. For the information values about behavioral variables, it was not useful to do the decoding and confusion matrix information computation because the space of the behavioral variables was larger than that of the neuronal activity, and thus we used a direct calculation of information through Shannon formula from the response probabilities. To correct for the limited sampling bias in the information estimation, we computed a shuffled information distribution by repeating the same process after shuffling the label for the trial type (for trial type information Eq. M9) or the neural activity (for the Eq. M10 information) 1000 times and subtracted the mean of the shuffled distribution from the estimated mutual information to correct for the bias^78^.

### Calculation of noise correlations from pairs of neurons

We computed the noise correlations between each pair of neurons (Figures 4, 5, 6) as rank correlations using the estimated cumulative distributive function (CDF) computed with the copula. We first computed the rank of each activity at each time *t* by evaluating its CDF computed through the copula (CDF is a measure of rank percentile^79^, because it measures the fraction of points with a value lower than the considered one). To compute the noise correlation, we pooled the CDFs computed for the activities in single trials, from the same trial condition, over the first two seconds after the test cue onset and computed the Spearman correlation coefficient. To compute the CDFs at fixed trial type, we first computed the conditioned marginal density functions *f*(*n*|*t*, Γ) for each neuron and to compute noise correlations at fixed trial type conditioned on the behavioral variables, we used the NPvC to compute the conditional marginal density functions 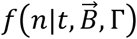 which allows us to specify the rank specific to the observed value of the behavioral variables at the considered time point for each neuron. To compute the CDFs for each of these density functions, we first computed the CDF over a grid with 200 points on the *n* axis ranging from the minimum to the maximum possible neural activity values by numerical integration of the density function (using the integral definition of CDF). We then used the CDF values on the grid to compute the CDF at each value of neural activity in each trial and time point by interpolation over the grid. Using CDFs and copulas to compute noise correlations has been shown to be robust and accurate in the estimation of noise correlations in neurons^80^.

We used Spearman correlation coefficients of the neural activity ranks computed from the copula rather than from the real data because the calculation of the ranks conditioned on the behavioral variables could not be performed directly from the data (because the behavioral variables span a continuum that cannot be sampled with a finite number of data points).

### Estimation of interaction information and of NPvC models for pairs of neurons

Following previous work^81^, for any pair of neuron with activity *n*_1_, *n*_2_, the interaction information about a task variable for the pair of neurons was defined as the difference between the pairwise information *I*^pairwise^(*n*_1_, *n*_2_; *C*) about the task variables carried by the joint observation of the pair of neuron (which reflects both single neuron properties and the effect of interactions between neurons) and the independent information *I*^ind^(*n*_1_, *n*_2_; *C*), which reflects only single cell properties and is defined the information for the pair of neurons when they are conditionally independent.

Importantly, in computing interaction information, we also conditioned over the movement variables (as we did for as for single neuron information) in order to isolate the part of the interaction that is related to the task variable selectivity. Conditioning removes the interaction emerging from shared movement selectivity and correlations between the task and movement variables. Thus, interaction information was computed as follows:

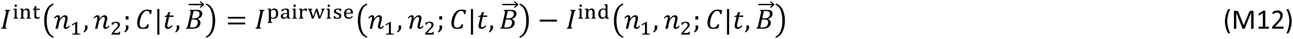

To compute the full and independent information components, we first estimated the pairwise and independent probability density functions 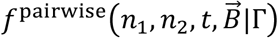 and 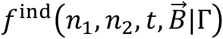 for each trial type respectively. The full probability density function was estimated using the following breakdown of the density function

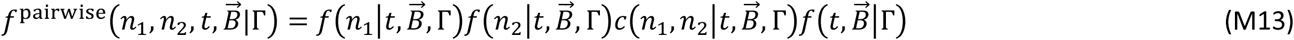

The independent probability density function was estimated using a similar breakdown

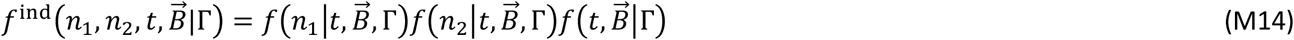

The extra component in the full density function, Eq. M13,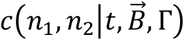, is the neuron-pairwise copula term, which represents the interaction between pairs of neurons for the fixed trial condition Γ and conditioned over the movement variables 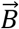 at any time *t*. This term was computed (conceptually similarly to the computation of conditional noise correlation explained above) by building a bivariate copula between the values of conditioned probability functions of the pair of neurons we use the definition of copula as the density function between the cumulative density functions of data,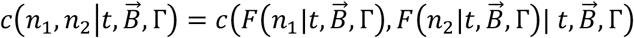, where *F*(·) denotes the cumulative density function computed from integrating over the density function. To compute this copula, we assumed that the conditional copula density is independent from the value of the movement variables at each time point, i.e.,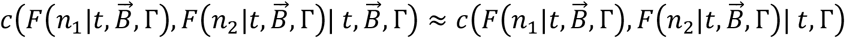, also note as simplifying assumption. We note, however, that this quantity still depends on the average shared co-tuning of the two neurons to movement variables through the single neuron cumulative density functions. This copula is estimated by pooling the values of conditioned probability functions at each time point for a given trial type.

To compute each of the conditional marginal densities 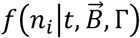, where *i* indicates the neuron index, we used the single neuron NPvC models we fitted for single neurons and the following identity

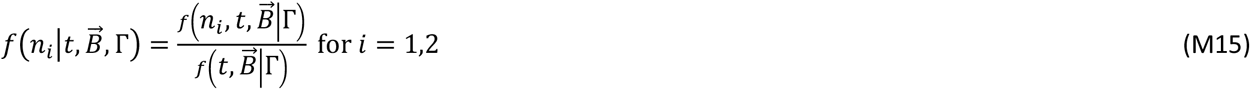

The density functions 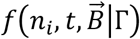 and 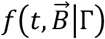 were numerically estimated as explained in Section Methods: Probabilistic models of neural activity. All these densities functions were estimated using the same 5-fold cross-validation method explained in Section Methods: Probabilistic models of neural activity.

After computing the density functions of Eqs. M7-M9, we used the same decoding approach as for single neuron information values (see Section Methods: Mutual information estimation) to compute the information values in the right-hand side of Eq. M12,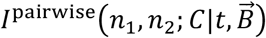 and 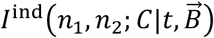 at any time point. The only difference compared to the single neuron information computation is that to compute the decoded task variable *Ĉ* (which is used later to compute the confusion matrix), we used the pairwise density functions (Eqs. M14, M15) instead of single neuron density functions.

We labeled each pair of neurons as information-limiting, independent, or information-enhancing if the mean of the interaction information for the pair in the first two second after the test cue onset was significantly negative, not different from zero, or positive, respectively. We computed the statistical significance of the sign of pairwise interaction information by building a bootstrap distribution of the mean interaction values (1000 bootstraps) and computed a p-value using a signed t-test for the probability that the mean of this distribution is either positive (p < 0.01, signed t-test), negative (p < 0.01, signed t-test) or indistinguishable from zero (p > 0.01, t-test) after applying a Holm-Bonferroni multiple comparison correction over all pairs ^82^. In total, we had 145,439 non-labeled pairs and 1355 same-target projection pairs.

### Triplet statistics and global clustering coefficient

To compute the triplet probabilities presented in Figure 5, we counted the number of each triplet type in each session and divided this count by the total triplet count (in total n = 1,080,382 triplets of non-labeled neurons and n = 30,204 triplets of same-target projections neurons). We estimated the statistical standard errors of the mean for these probabilities with bootstrapping (100 bootstrap subsamples). Since we were interested in deviations of triple probabilities from the values expected in networks with randomly structured interactions, for each bootstrap subsample, we computed an estimate of the probability of each triplet type in a random network by shuffling the pairwise interactions across neurons within each session. Importantly, this shuffling did not change the total number of information-liming, information-enhancing, and independent pairs within the network. The relative triplet probability was computed as the difference between the real triplet probabilities and the shuffled probability, averaged over 1000 shuffles.

We also computed global clustering coefficients ^40,41^ to detect the presence of subgraph clusters of information-limiting or information-enhancing pairwise interactions within the network. We computed the global clustering coefficient separately for subgraphs of only information-limiting and only information-enhancing pairwise interactions. To compute the information-limiting (information-enhancing) global clustering coefficient, we first defined a binary graph in which neuron pairs have an edge if the pair has a significant value of information-limiting (information-enhancing) interaction information. If the pair does not have significant interaction information, this pair does not have an edge. For each of these two graphs separately, we then counted the number of closed triplets (triplets with edges between all three neurons) and open triplets (with only two edges between the three neurons) and computed a clustering coefficient (*CC*) as follows^40,41^:

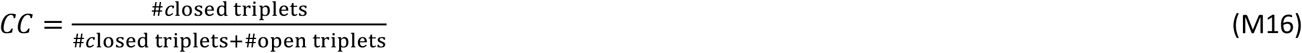

The *CC* ranges between 0 and 1, with larger values corresponding to a graph with larger numbers of closed triplets, which means the existence of clusters of nodes that are more connected within themselves compared to other nodes. A graph with a cluster of nodes connected (similar to the middle panel of Figure 5A) will have few open triplets because any selection of three nodes will result in either a closed triplet (when the three nodes are selected from the cluster) or will have only one edge, and thus using Eq. M16, the clustering coefficient will be close to or equal to one. We also computed a relative clustering coefficient as the difference between the clustering coefficient computed on data and the one computed on a randomly organized network, using the shuffling procedure described above. Statistics were assessed by bootstrapping as explained above for the triplet probabilities.

### Parametric two-pool model of network structure

To better understand the relation between network structure and triplet probabilities, we considered a simple parametric model for the network structure within the framework of well-studied approach of network clustering modeling known as planted partition model or stochastic block model^83^. We considered a network with *N* nodes (corresponding to single neurons) that are connected with either a positive interaction with probability *P*(+) or with a negative interaction with probability *P*(−) = 1 − *P*(+). We considered a simple structure consisting of two pools of size *N*/2, with each pool having different positive and negative probabilities of (*P*_1_(+), *P*_1_(−)*P* = (*P*(+) + δ*P*_1_, *P*— δ*P*_1_) and (*P*_2_(+), *P*_2_(−)*P* = (*P*(+) + δ*P*_2_, *P*— δ*P*_2_) where δ*P*_1_ and δ*P*_2_ are the two parameters that express the difference of information-enhancing probabilities within each pool relative to the average information-enhancing probability in the full population. Each value of (δ*P*_1_, δ*P*_2_) defines a network structure. A network with a uniform distribution will have (δ*P*_1_ = 0, δ*P*_2_ = 0) while other combinations of (δ*P*_1_, δ*P*_2_) generate two-pool networks with extra information-limiting or information-enhancing interactions within the pools (as is shown in Figure 5F). Considering the network is fully connected, we computed analytically the probabilities of the edges connecting the two pools in terms of (δ*P*_1_, δ*P*_2_) for fixed values of (*P*(+), *P*(−)*P*, as follows:

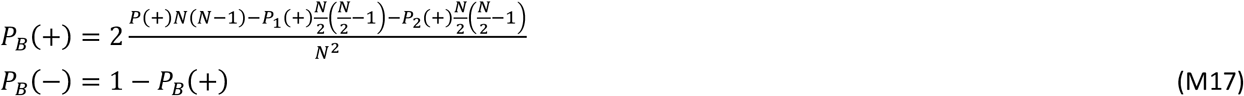

Because in the model we specified the full probability of connections between the two pools, we could also analytically compute the probabilities of each triplet type both for the structured and shuffled networks as described in Supplemental Information: Two-pool model of network structure.

### Motif expansion of population information to identify the contributions of structured and unstructured interactions to population coding

To understand how the structure of interactions among different neurons contributes to population coding, we computed an analytical expansion of the information carried by the activity of a population of neurons about a task variable as a function of the network motifs in which interaction information between pairs of neurons is structured. As is often done in models of population coding^46,84^, we assumed that neural activity for a fixed trial type follows a multivariate Gaussian distribution. Further, as we found in our data, when computing the expansion, we assumed that the population had a finite size, that single neuron information values were small, and that pairwise noise correlations between the neurons were small, which (as we will show) in turn implies that pairwise interaction information values are smaller than single neuron information values (again, as found in our data). With these assumptions, we approximated the information carried by the population as a sum of terms that can be directly related to the structure of interaction information within the network. The details of this expansion and assumptions are presented in detail in Section Supplemental Information: Analytical expansion of the population information.

We first expressed, as is often done in studies of population codes and it follows from extending to arbitrary population size the concept of pairwise interaction information in Eq. M12, the full population information as the sum of two components: the independent information *I*^ind^, which is the information of a population with the same single neuron information values as found in the data, but with no interaction between them (where interaction is defined as noise correlation, that is a statistical relationship between the neurons at fixed trial type), and the interaction information *I*^ind^, defined as the difference between the real and the independent population information:

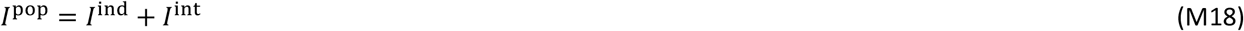

The independent information *I*^ind^ is a function of only the single neuron information values (see Eq. SM21). The interaction information quantifies the total effect of noise correlations on the stimulus information carried by the population. We decomposed the population interaction information into a sum of different components expressing the contribution of different graph motifs of the interaction network (a graph in which the edges correspond to interaction information values). As a result, the interaction information is a function of the single neuron information values, the interaction information values, and the number (probability) of triplet motifs (see Eq. SM31). To assess the total contribution to information representation of network structure (triplet-wise arrangements of pairwise links of interaction information) with respect to an unstructured, random arrangements of the same links within the network, we then rearranged this expansion to break down interaction information into two components:

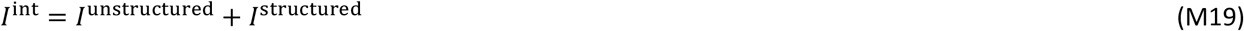

These two terms are calculated in (Supplemental Information: The population information component corresponding to the network organization of pairwise interactions), and their equations are reported in Eqs. SM32 and SM33. The unstructured interaction information component is the information in a population with the same distribution of pairwise interactions as the data, but randomly rearranged across neurons by shuffling (thus no network structure), minus the independent information. This component quantifies the contribution of the distribution of pairwise interactions in the population information. The structured interaction information component is defined as the difference between the total population information of the original network and the population information in an unstructured network. Thus, this component isolates the contribution of the structure of pairwise interactions in the network.

To compute the unstructured and structured triplet interaction information, we used Eq. SM31 after computing the number and weights of different motifs using the real and the shuffled networks (see Supplemental Information: The population information component corresponding to the network organization of pairwise interactions). For calculating information, we used a complete and general expansion of population information that is valid also for non-symmetric networks (Supplemental Information: Simplifying the structured interaction information, and Figure S7). However, in Figures 6B-D, we illustrate this expansion for the simple case of a symmetric network, which describes well our empirical data. In fact, we prove in the Supplemental Material that, in the case of a symmetric two-pool network, these computations can be greatly simplified because of symmetries present in the network. In the symmetric case, the only motif contributing to the structure component of the interaction information is the closed triplet (Supplemental Information: Analytical expansion of the population information) and its associated triplet probabilities and weights. As shown in Figure 6C, we can define four different types for triplets with different combinations of information-enhancing and information-limiting interaction links. Two of these motifs with even number of information-limiting (‘+,+,+’, ‘+,-,-’) contribute positively to the structured interaction information and the other two with odd number of information-limiting interaction links (‘+,+,-’, ‘-,-,-’) contribute negatively. The strength of contribution of the triplets to the structured interaction information is given by the deviations from the shuffled-network-level of the probability of each triplet type (computed as described in Methods: Triplet statistics and clustering coefficient), multiplied by a non-negative numerical coefficient that is function of the single-neuron and pairwise interaction information values that shape the triplets. Equations for these terms are reported in Eqs. SM46, SM47. As it is shown through analytical calculations in supplementary Information (Eq. SM31), one important property of these three terms is the difference in how they scale with population size. While the independent information scales linearly, the unstructured and structured interaction information terms scale faster with second and third orders of population size respectively.

## Supplemental Information

### Analytical expansion of the population information

Here we present in detail an analytical expansion of how the information carried by a population of *N* neurons depends on the values of single neuron information, on the values of pairwise interaction information, and on how pairwise information values are organized at the network level. We also present how this population information grows with the population size N. This expansion is based on a small set of assumptions. First, following established population coding models, we assume that neural population activity for a fixed trial type follows an N-dimensional multivariate Gaussian distribution. Second, we assume that single neuron information values and noise correlation values are small, assumptions well supported by our data. Third, we consider arbitrary but finite population sizes N (the largest population size for which the expansion is still accurate depends on the precise values of single neurons information and correlation values set by simple equations reported below).

To estimate the information carried by a population of *N* interacting neurons in an analytically tractable, yet sufficiently general way, we model the probability density of the activity of a population of *N* neurons, ***r***, in a trial condition, *c* (which, in our data, would be sample cue, test cue, reward location or choice), as a multivariate Gaussian, *p*(***r***|*c*) = 𝒩(***μ***_c_, **Σ**_c_) with ***μ***_c_ and **Σ**_c_ indicating the mean vector and covariance matrix for each trial condition. We assume two trial conditions, *c* = 1,2 having equal prior probability. Thus, the marginal distribution of population activity is given by a mixture of two Gaussian distributions, 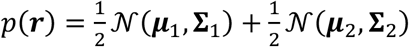. For simplicity, we consider a constant covariance matrix across trial conditions, **Σ**_1_ = **Σ**_2_ = **Σ** (trial-independent noise correlations). We express the covariance matrix as a function of a diagonal matrix, **σ**, with elements 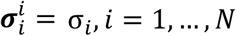 corresponding to the standard deviations of single neuron responses, and a correlation matrix, ***R***, as **Σ** = **σ**^*T*^**Rσ**. The correlation matrix consists of a unit diagonal and symmetric off-diagonal elements, 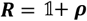, where 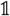 is the unit matrix and 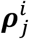 are the pairwise noise correlations, with 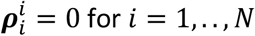.

The mutual information between the population response, ***r***, and the stimulus, *c*, is computed as *I*_pop_(***r***|*c*) = *H*(***r***) − *H*(***r***|*c*), where *H*(·) denotes the entropy of the response probabilities *p*(***r***) and *p*(***r***|*c*). The entropy of the conditional response probability, *H*(***r***|*c*), is computed using the analytic form of the entropy of a Gaussian distribution (Appendix 1). The entropy of a Gaussian mixture, *H*(***r***), does not permit an analytic closed-form expression. We computed the entropy by approximating the mixture distribution, *p*(***r***), with a Gaussian distribution, which is the maximum entropy distribution given its first two moments. Using this approximation, the mutual information can be written as (see Section Supplemental Information: Additional mathematical details of the calculation of the information expansion for details):

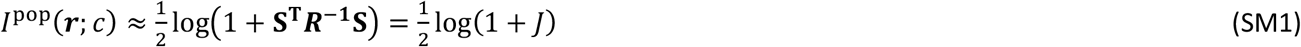

where 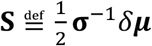 and δμ = μ_c=2_ – μ_c=1_. In order to obtain a compact form, we introduced the quantity 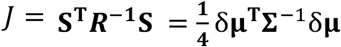, which corresponds, up to a multiplicative factor, to the discrete formulation of the linear Fisher information ^85^. We used the maximum-entropy approximation because it is accurate under our conditions and leads to more compact equations. However, we also took an alternative approach that consists of a Taylor expansion of the information, *I*^pop^, up to the second order in J (the order needed to take into account triplets). We verified that, by using this approach, the terms of the structured and unstructured interaction information were the same as those found with the maximum-entropy approach at the leading order, and exhibit only minor differences in numerical coefficients, but not in sign, at higher orders. (see Section Supplemental Information: Additional mathematical details of the calculation of the information expansion). Therefore, the results found with this more compact derivation are quantitatively and qualitatively comparable to those obtained with other approaches.

We define the independent information as the information obtained when the noise correlations are set to zero,

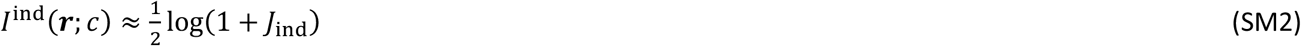

where *J*_ind_ = S^T^S.

We then define the interaction information as the difference between the population information (Eq. SM1) and the independent information (Eq. SM2),

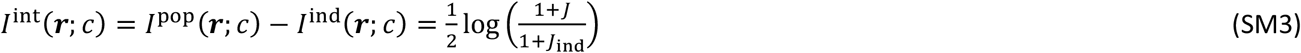

We simplify this expression by expressing the inverse of the correlation matrix in powers of ***ρ***, 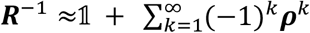 (representing a Neumann series). This is valid assuming that the noise correlations are small and λ_1_, the largest eigenvalue of ***ρ***, is smaller than 1. The largest eigenvalue depends on the distribution of the noise correlations. By assuming that the entries of the correlation matrix are drawn from a Gaussian distribution with a mean of zero and standard deviation equal to *s*, the largest eigenvalue will be of order 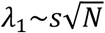. If instead we assume that all the correlations are equal to *ρ*, the largest eigenvalue scales faster with matrix size as λ_1_∼*ρ*(*N* − 1).

The Fisher information term, J, can be written as *J* = *J*_*ind*_ + *J*_*cor*_, with a ‘correlational’ information term *J*_*cor*_ corresponding to the sum of different powers of the correlation matrix, with increasing powers expressing increasing orders of interaction. The term *J*_*cor*_ takes the form:

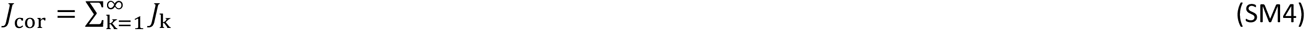

with

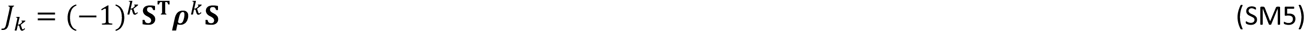

Each of the terms *J*_*k*_ is bounded by the corresponding power of the largest eigenvalue of the correlation matrix 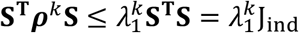, from which it follows that:

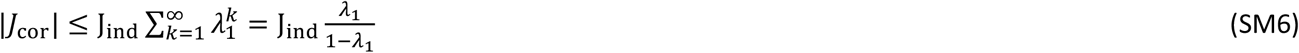

By defining the scaled quantities 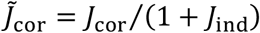 and 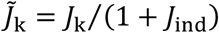,Eq. SM3 can be written as

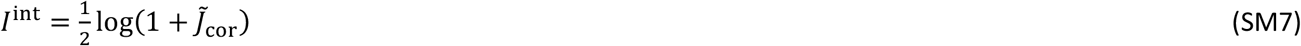

We then expand Eq. SM7 in orders of 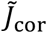 as follows:

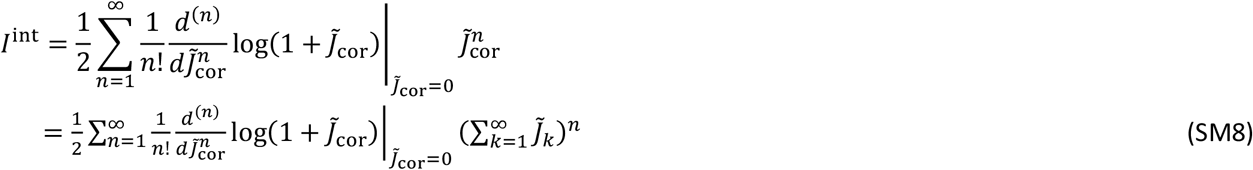

We rearrange the terms in Eq. SM8 in orders of interaction,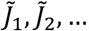,as follows:

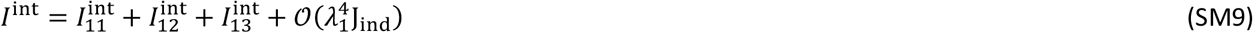

where each term is extracted from expanding Eq. SM8 as,

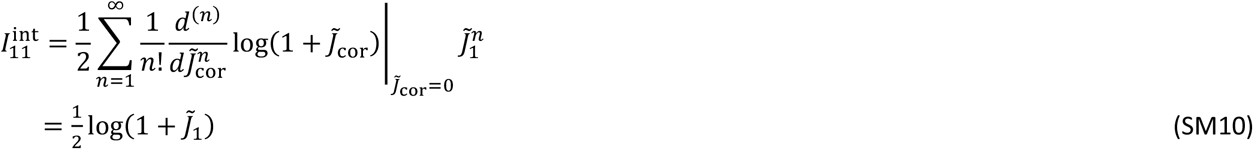

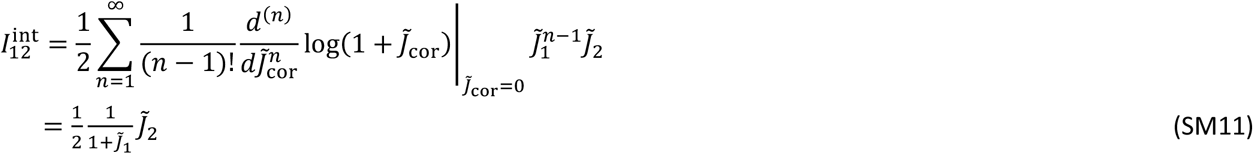

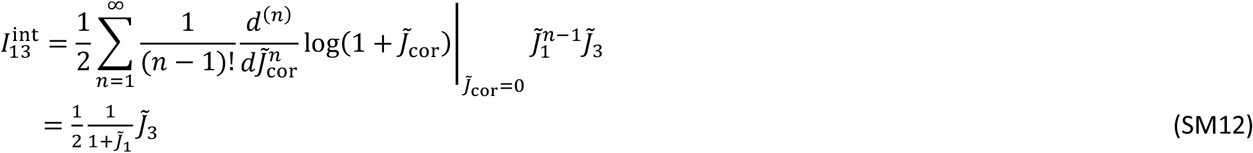

We ignored the linear term of 4th order as it is proportional to 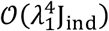. The term proportional to 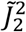

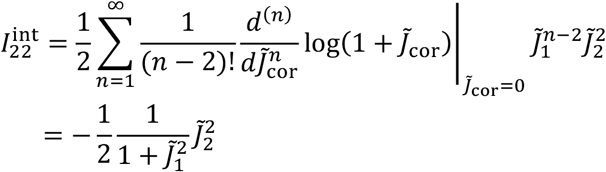

is of order 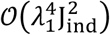 and can be ignored by assuming λ_1_*J*_ind_ < 1 (further, we note that the approximation in Eq. SM2 breaks down when *J*_ind_ > 3, since the mutual information is bounded to *I*^ind^ < 1bit, implying 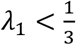).

We then express the different orders of interactions in Eq. SM5 in terms of single neuron Fisher information, *J*_ind,*i*_ and first-order pairwise Fisher information elements, *J*_1,*ij*_, as

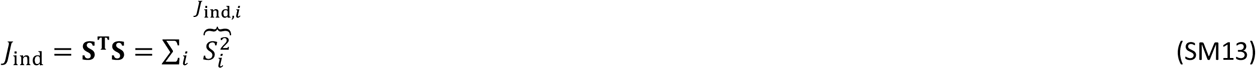

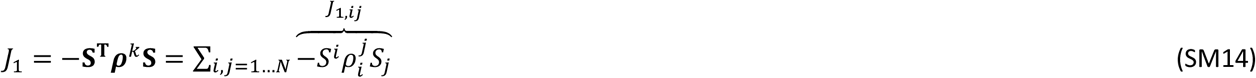

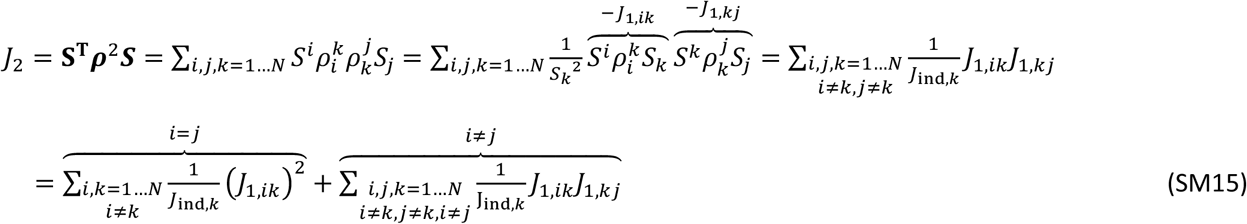

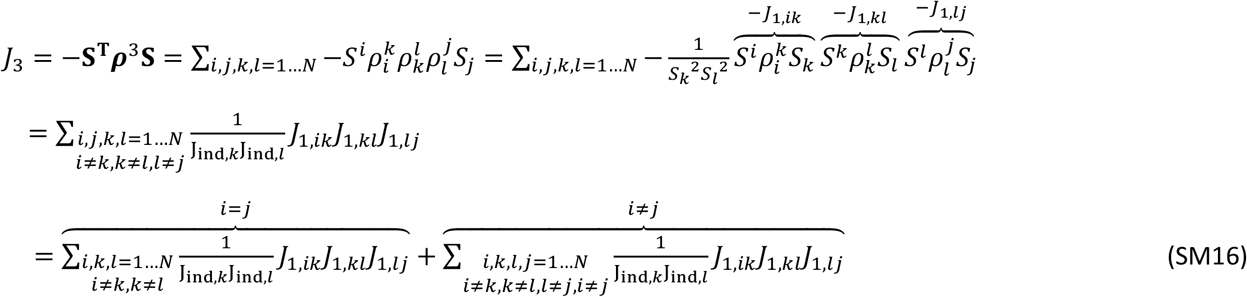

Using Eqs. SM9-16, the interaction information can be computed as a sum over different components, which are computed using *J*_1,*ij*_ and *J*_ind,*i*_. The summations over the pairwise components are structured such that we can associate them to weighted functions over graph motifs. We can then express Eq. SM9 as a sum over different graph motifs, which appear in the summations of SM13-16 and are shown in Figure S8A:

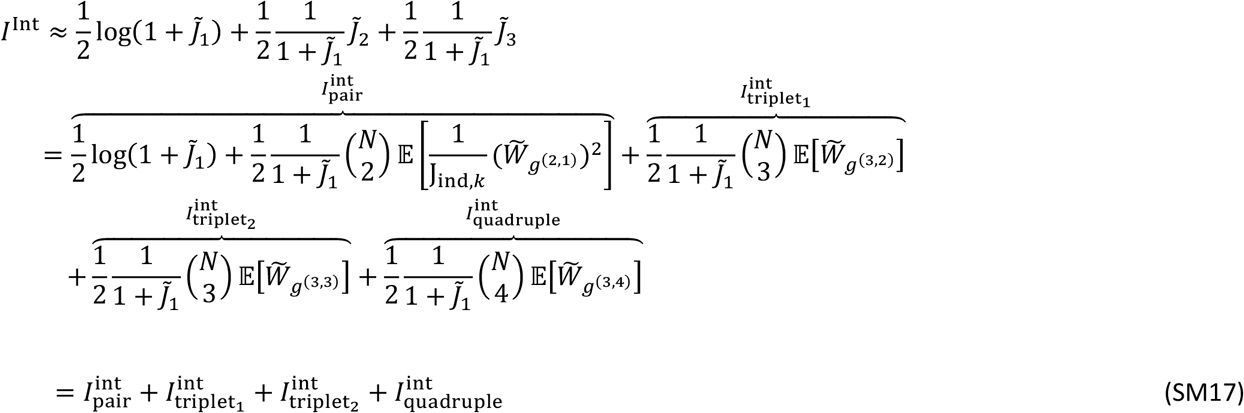

The result is a sum over terms corresponding to different graph motifs *g*^(*n,e*)^ that are connected subgraphs with *n* nodes (or single neurons) and *e* edges (or pairwise interactions). Each term is an expectation of weights 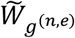 (or interaction weight per motif) computed by averaging over all the subgraphs belonging to a motif *g*^(*n,e*)^ (Figure S8A), multiplied by the number of each motif in the graph, which is a binomial coefficient of the number of nodes and defines the scaling of the corresponding information component as a function of the population size. For any graph with a set of nodes and links, the weights 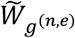 can be expressed in terms of pairwise Fisher information J_1,*jk*_ and single neuron Fisher information *J*_ind,*k*_ using SM13-16 as follows

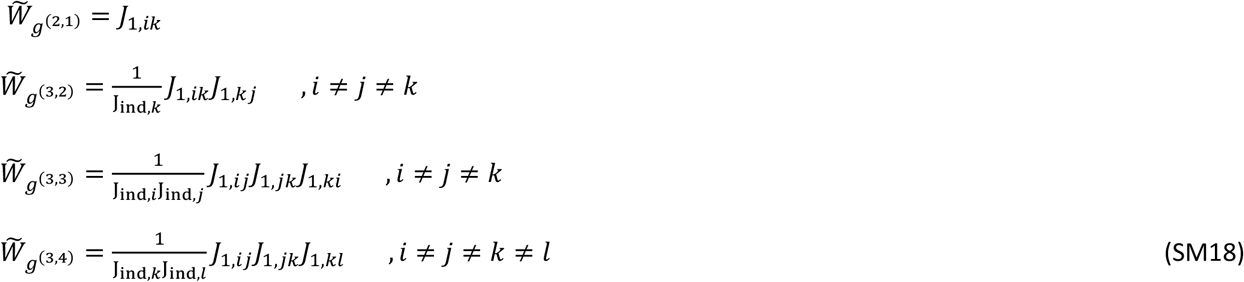

### Population information expressed in terms of single-neuron and pairwise interactions information values

Here, we rewrite the information breakdown, Eq. SM17, as a function of single neuron information and single pair interaction information. This makes it easy to compute the population information from the values of single neuron and pairwise information in the population computed from the data and reported in Figures 3-4 of the paper.

The single neuron information, *I*_*j*_, is the independent information conveyed by a single neuron. Using Eq. SM2 for *N* = 1, we have that

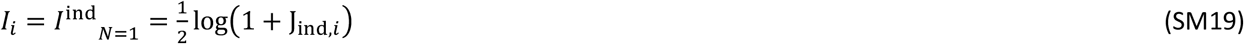

By inverting this equation, we obtain the single neuron Fisher information as a function of the single neuron information,

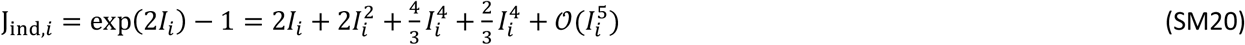

By inserting Eq. SM20 into Eq. SM2, we get the independent information of a population of size *N*

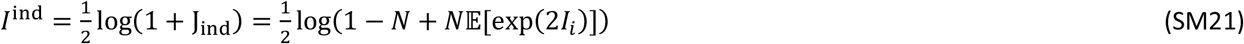

*Wh*ere 𝔼[exp(2*I*_*i*_)] is the average of exp(2*I*_*i*_) over population.

The pairwise interaction information corresponds to the interaction information of a population of size *N* = 2

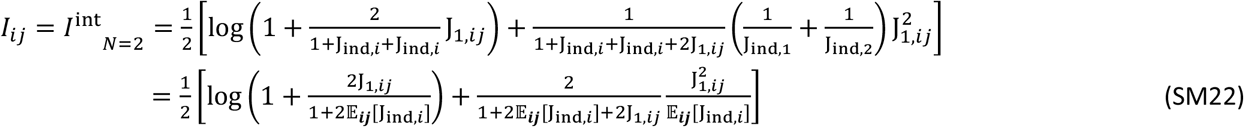

Where 𝔼_*ij*_[·] indicates the average over the pair of neurons. Here the second term comes from the elements of J_2,*ij*_ in Eq. SM11 for one pair. In the last approximation, we assumed that J_ind,*i*_∼J_ind,*j*_∼𝔼_*ij*_[J_ind,*i*_]. In general, the ratio between pairwise and single-neuron Fisher information scales as

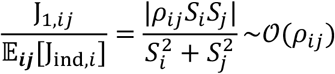

In order to simplify Eq. SM22, we assume that this ratio, which is of the order of the noise correlation, is of the same order as the single neuron information, i.e. *ρ*_*ij*_∼𝒪(𝔼[J_ind,*i*_])∼𝒪(𝔼[I_*i*_]). This assumption allows us to expand everything to the order of the single neuron information. In what follows, we use the short notation 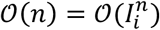 to denote the expansion order.

By expanding Eq. SM22 up to the 4^th^ order to match the order of the information corresponding to the closed triplets (the main graph motifs that are of interest in our data given the results shown in Figure 5), we obtain the following equation

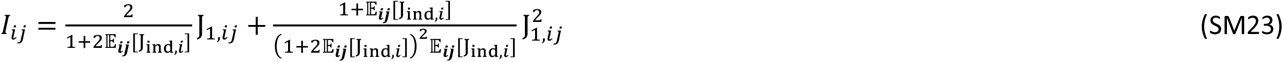

Among the two solutions to Eq. SM23, we pick the one consistent with the linear approximation (i.e., the equation obtained when the term of order 2 is ignored), which yields

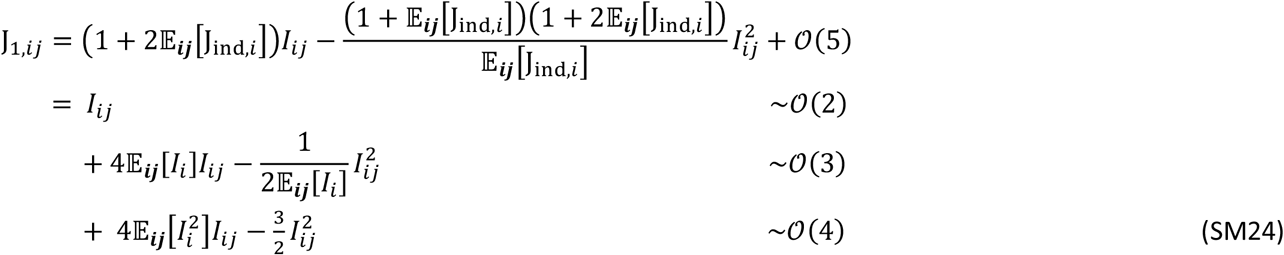

where we substituted the Taylor expansion of J_ind,*i*_ Eq. SM20, and we truncated the expression up to the 4^th^ order.

Given the approximations for J_1,*ij*_ and J_ind,*i*_ in Eqs. SM19-24, we can compute the terms in Eq. SM9.

The first term can be written as

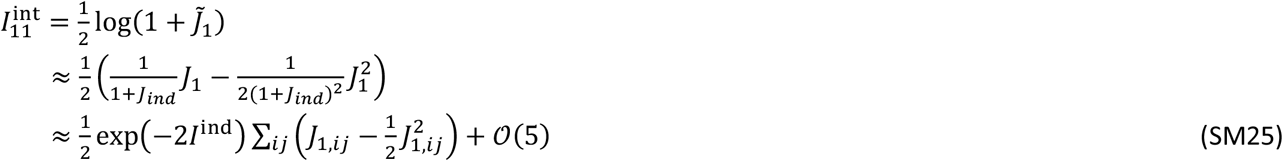

where we kept only terms up to the 4^th^ order and in the last line we substituted Eq. SM21. In performing the sum over the pair indices, we have *J*_1_ = ∑_*ij*;_ *J*_1,*ij*_ where *J*_1,*ij*_ is computed from Eq. SM24. We further make the approximation that the average of the single neuron information and the interaction information are uncorrelated across pairs, i.e. 𝔼[𝔼_*ij*_[*I*_*i*_]*I*_*ij*_] ≈ 𝔼[*I*_*i*_]𝔼[*I*_*ij*_]. This approximation is useful to simplify the equation, and it is accurate under the assumption that the correlation between the independent information of a pair of neurons and their interaction information is small, which we verified in our data. Further, we approximate the inverse of the average with the average of the inverse, 𝔼[1/𝔼_*ij*_[*I*_*i*_]] ≈ 1/𝔼[*I*_*i*_]. We finally obtain an expression that depends only on the statistics of the single neuron and pairwise interaction information across the population as follows,

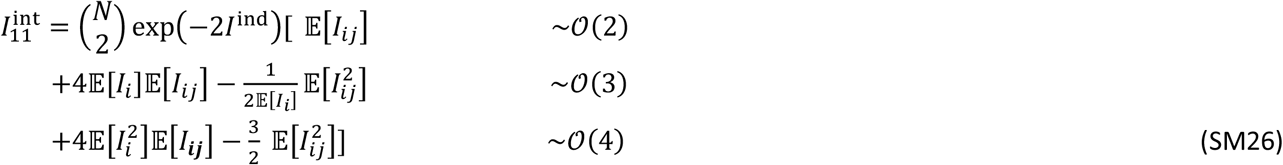

In order to maintain a compact notation, in this expression and in what follows, we preserve terms which are 𝒪(5) coming from the expansion of exp(−2*I*^ind^).

The second term of the interaction information can be rewritten as

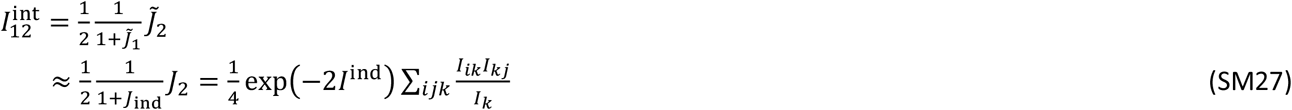

where in the first approximation we ignored terms proportional to 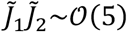, and in the last line we used the definition of *I*^ind^, Eq. SM2.

The contribution of the 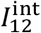 can be decomposed into two terms, one related to the pairs and the other to the open triplet graph motifs

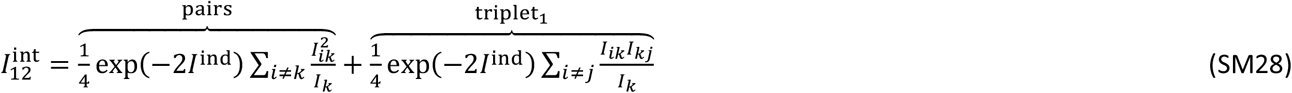

Similarly, the third order term can be expressed as follows,

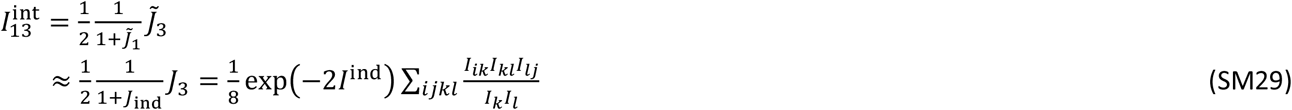

which can be decomposed into the closed triplet and quadruplet components

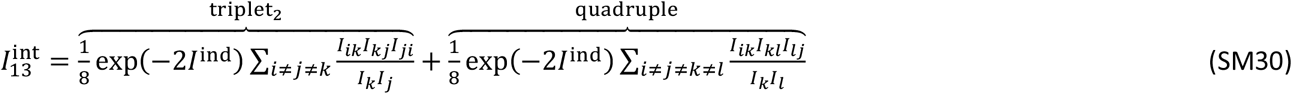

Putting back Eqs. SM26, SM28, and SM30 into Eq. SM9, we can write, similarly to Eq. SM17, the interaction information as a weighted sum over different graph motifs

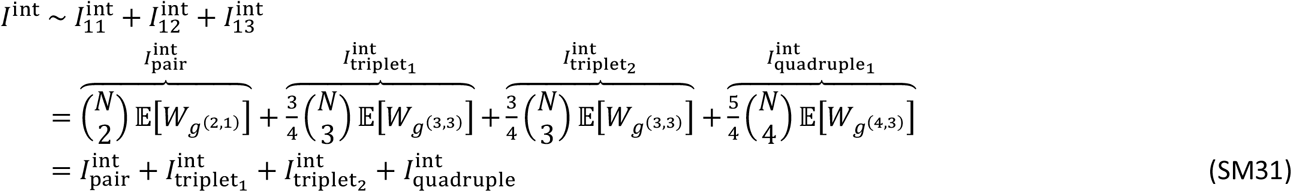

where the interaction weight per motif, *W*_*g*_(*n,e*), for each network motif is computed in terms of pairwise interaction information and single neuron information values present in the motif using Eqs. SM26, SM28, and SM30. These values are shown in Table. S1.

### The population information component corresponding to the network organization of pairwise interactions

The population interaction information presented in Eq. SM31 consists of two types of terms. Some of the terms only depend on the distribution of the pairwise interaction information values and are independent of the way pairwise interactions are structured in the network. These components include (but are not limited to) the terms related to the first and second order motifs *g*^(1,0)^ and *g*^(2,1)^ since these terms do not depend on the way the pairwise interaction values are structured in the network and depend only on the single neuron information and pairwise interactions in the network. On the other hand, there are components of the interaction information that depend not only on the distribution of the values of pairwise interaction values but on the way these interactions are structured in the network. For example, if we consider a *g*^(3,2)^ graph, the interaction weight *W*_*g*_(3,2) is proportional to the product of the interaction information values *I*_*ij*_ and *I*_*jk*_ of the two edges and will be positive for the cases in which both edges of the *g*^(3,2)^ graph are information-enhancing or information-limiting and is negative when one edge is information-enhancing and the other edge is information-limiting. So, changing the structure of interactions values in the network (without changing the distribution of values) can change 𝔼 [*W*_*g*_(3,2)] since we can generate networks with different numbers of positive and negative interaction weights *W*_*g*_(3,2) from the same distribution of pairwise interactions. We can separate these two components of population information by removing the structure of pairwise interactions while keeping the distribution. This can be done by a shuffling transformation *I*^*sh*^ → Φ^*sh*^(*I*), in which the values of the elements of the interaction matrix *I* are redistributed into a randomly organized matrix while keeping the interaction matrix symmetric and without changing the distribution of values of the matrices 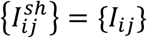, where 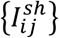 and 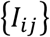 are the sets of pairwise interaction information values in the network. This transformation can be used to find the component of the population interaction information that originates from the structure of pairwise interactions in the network. The component of the population information related to the network structure is then defined as the difference of the population information using the real interaction matrix and the population information computed using the shuffled interaction network as it is shown in Figure 6C.

As defined above, the population interaction information is the difference between the population information and the independent population information. Given this definition, the population interaction information can be decomposed into a sum of the structured interaction information (defined above) and a term that is the difference of the population information computed from the shuffled interaction network and the independent information and is called here the unstructured interaction information (Figure 6C).

The unstructured (*I*^unstructured^) component of the interaction information can be expressed in terms of the original interaction matrix *I* and the shuffled interaction matrix *I*_*sh*_ as follows:

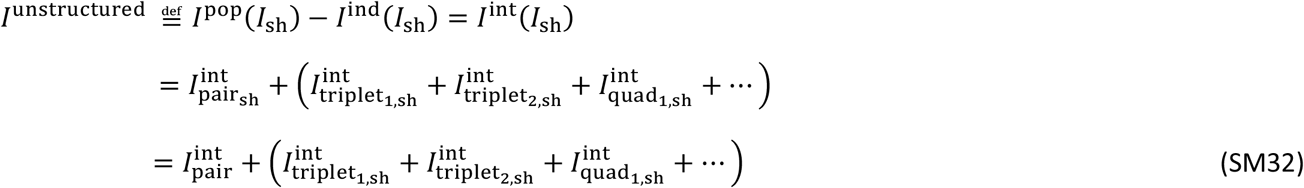

where 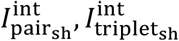 and 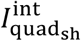 are computed the same way as the non-shuffled ones, using Eq. SM31, but for a network with shuffled edges or shuffled interaction information matrix instead of the true interaction information matrix. In the last line, we used the fact that shuffling does not affect the expectations of the single pair motif and so 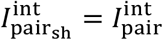 by definition of the shuffling procedure.

The structured (*I*^structured^) component of the interaction information is defined as the difference of the population information computed using the real and the shuffled interaction networks as:

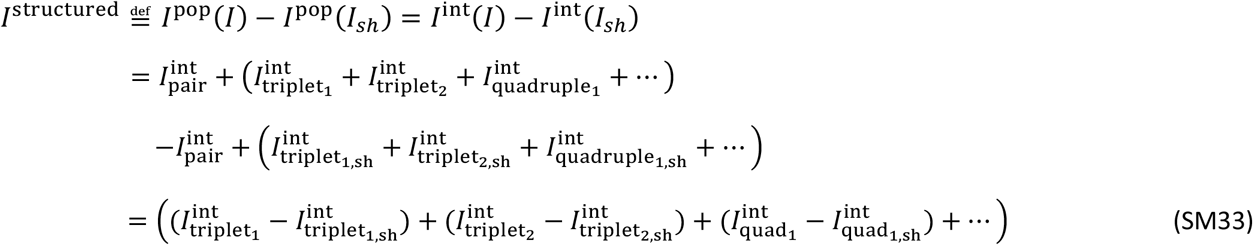

Where in the first line we used the fact that the original and shuffled interaction networks have equal independent information. The interaction information *I*^ind^ and the full population information *I*^pop^ can then be represented as the sum of the unstructured, structured and independent components:

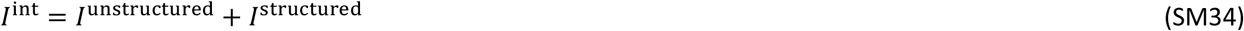

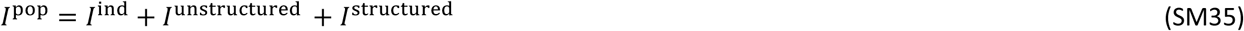

This breakdown makes it possible to understand the contribution of basic factors underlying information processing in a network including distribution of single neuron information values, distribution of pairwise interactions, and the network structure of the pairwise interactions.

### Relating structured interaction information and triplet probabilities

The interaction information of a network can be estimated by sampling different subgraphs of each motif *g*^(*n,e*)^ and computing the interaction weights per motif *W*_*g*_(*n,e*) for each subgraph. This can be achieved by using analytic forms presented in Table S1 to estimate the expectation of the interaction weights for each graph motif and then using Eq. SM31 to scale it to a population of size *N*. Although this is a general process and can be applied to any network with a general distribution of pairwise interactions, this formulation does not provide insight into how interaction information is related to the structure of pairwise interactions in the network.

Every motif is defined only by the number of nodes (neurons) and edges (pairwise interactions) of the subgraph. Simple graphs usually have binary edges, meaning that the connection between any two nodes is either present or absent. This is different for the interaction graph, which is not a binary graph. Each edge can get either a positive or negative value. For the interaction graph, each motif consists of a few different motif types (or sub-motifs) that have the same number of nodes and edges but differ in the number of positive (information-enhancing) and negative (information-limiting) pairs. For example, an open triplet motif is a graph consisting of three nodes and two edges, but it can take on three different sub-motifs corresponding to either ‘-,-’, ‘+,+’, or ‘-,+’ combinations for the two edges of the graph (as shown in Figure S9).

One way to understand better the structured interaction information is to break down the contribution of each motif into a sum over sub-motifs. To start, we represent the interaction weight per motif as a product of its absolute value 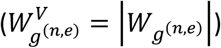 and its sign 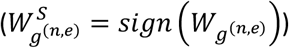

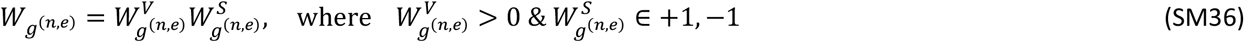

The interaction weight sign 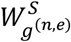 defines whether each subgraph contributes positively (information-enhancing) or negatively (information-limiting) to the population information based on its sign. For each motif *g*^(*n,e*)^, we call the subset of sub-motifs that generates information-enhancing interaction as 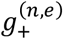 and the sub-motif that generates a information-limiting interaction as 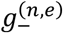, as shown in Figure S8B. For example, a *g*^(3,2)^ graph can contribute positively if both of its edges (interactions) have similar signs (either positive or negative), and it will contribute negatively if one edge is information-enhancing and the other is information-limiting. Similarly for triplets of *g*^(3,3)^ motif, there are two information-limiting and two information-enhancing types of graphs with different combinations of information-enhancing or information-limiting pairwise interactions, as it is shown in Figure S8B. The interaction information corresponding to each network motif was computed in Eq. SM31 in terms of the expectations of interaction weights.

Using Eq. SM36, we can rewrite the expectation value of the interaction terms into a sum over the interaction terms corresponding to each of the subgraphs 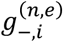 and 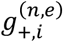 presented in Figure S8B (*i* is the index of the sub-motif with positive or negative sign)

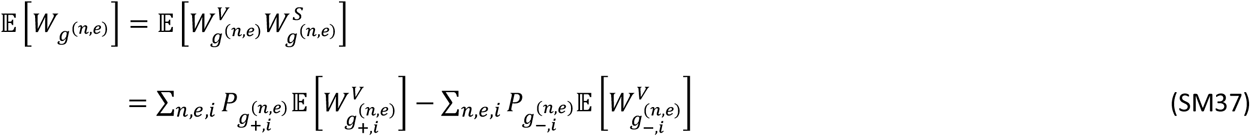

Where 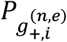 and 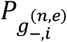 are the probability of the presence of each of the subgraphs shown in Figure S8B with positive and negative interaction weights in the network. These probabilities can be estimated by sampling enough subgraphs of each motif from the network. For the open triplet term, this can be written as follows,

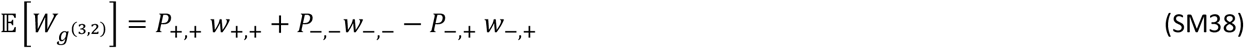

Where *w*^Ȳ^*s* are the expectations of the interaction weight values over the subgraphs with the given number of information-limiting or information-enhancing nodes, for example 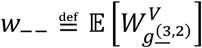. Similarly, *P*_−,−_,*P*_+,+_ and *P*_−_,_+_ are the probabilities of the three subgraphs of *g*^(3,2)^ shown in Figure S8B with different combinations of information-limiting (‘-’) or information-enhancing (‘+’) pairwise interactions.

The interaction weight expectation corresponding to the closed triplet subgraph *g*^(3,3)^ can be expanded as follows,

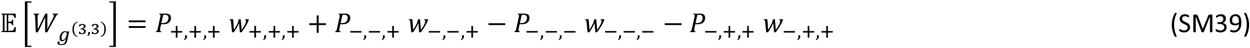

Where *P*_+,+,+_, *P*_−,−,+_,*P*_−,−,−_ and *P*_−,+,+_ are the probability distribution of the four types of triplets present in the network, each of which contributes with the corresponding weight value (*w*’s).

Given Eqs. SM38 and SM39, the triplet component of the interaction information can be estimated by first computing probabilities of the open triplets (*P*_+,+_, *P*_−,−_, and *P*_−,+_) and closed triplets *P*_+,+,+_, *P*_−,−,+_, *P*_−,−,−_, and *P*_−,+,+_) in the network and then computing the corresponding interaction weight expectations *w* over the network. Given basic properties of a network with every edge being either information-enhancing or information-limiting, the probabilities of open and closed triplets are related as follows,

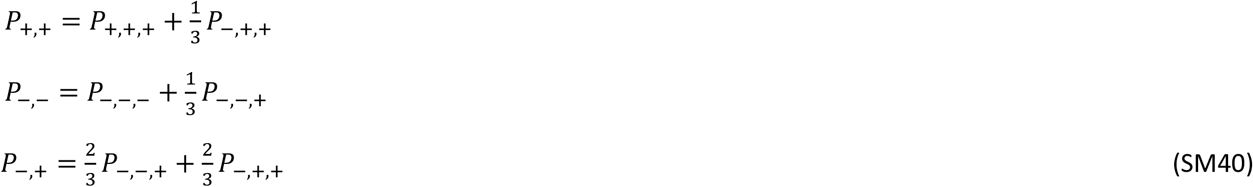

using the fact that each closed triplet consists of three open triplets. Note that we can easily include independent pairs in the calculations but given that the interaction weight per motif for a subgraph with an independent edge is zero, independent pairs do not contribute to the network components of the information. We therefore considered every pair to be either information-enhancing or information-limiting for simplicity. Using Eqs. SM39,40, the triplet components of the population information can be computed from the probabilities of four different closed triplet sub-motifs ‘-,-,-’, ‘+,+,+’, ‘-,-,+’, and ‘-,+,+’. Similarly, we can calculate higher order terms such as the quadruplet term by calculating the probabilities of each sub-motif and their corresponding interaction expected weights.

In general, each of the different sub-motifs shown in Figure S8B can have different interaction expectation weight values (*w*), but we observed in our data that the distribution of single neuron and pairwise interaction signs are approximately uniform, meaning that pairs of neurons with information-enhancing and information-limiting interactions have similar distributions of single neuron information and absolute interaction information values. Furthermore, we showed similar uniform distributions of single neuron and pairwise interaction values for different triplet sub-motifs and similar distributions of interaction motif values (Figure S6).

In the regime in which the interaction weight values are the same for different sub-motifs (for example when *w*_+,+,+_ = *w*_−,−,−_ =*w*_−,+,+_ =*w*_−,−,+_ the case of closed triplets), we can simplify the interaction weight expectations of Eqs. SM38-40 as follows,

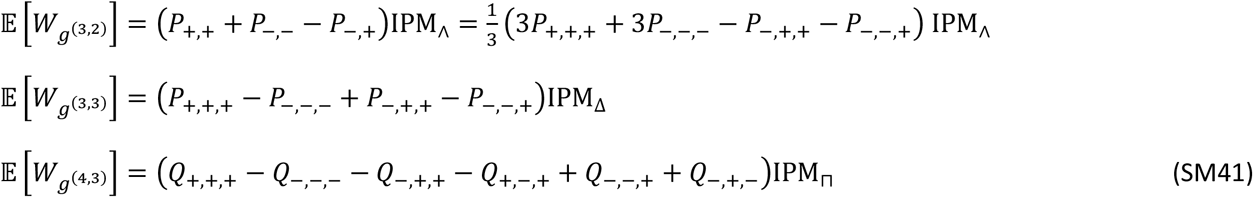

Where (*P*_+,+,+_, …) are the probabilities of different triplets and (*Q*_+,+,+,_ … ) are the quadruplet probabilities. We also used the notation, similar to what is used in Figure 6, IPM_∧_, IPM_Δ_, and IPM_⊓_, denoting the expected interaction weight values for open and closed triplets and quadruplets respectively. They are computed as an expectation over all the sub-motifs belonging to a motif (contrary to *w*_−−_,… etc which are expectations over sub-motifs with specific interactions)

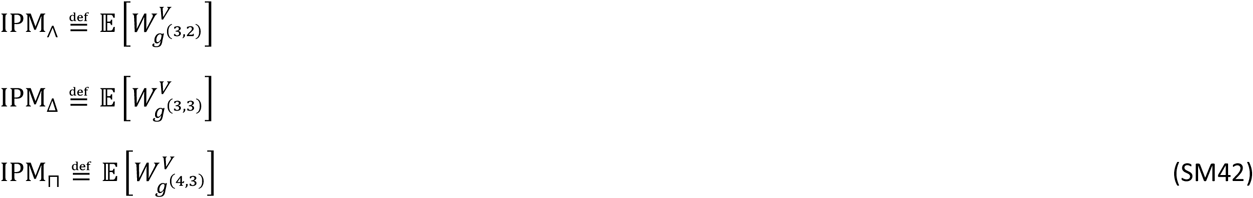

Where *W*^*V*^s are the absolute values of the interaction weights (Table. S1) and can be computed by sampling enough of triplets or quadruplets from the network (meaning that IPM is positive definite by construction). The other components are functions of triplets and quadruplet probabilities, which define whether the overall interaction term is information-enhancing or information-limiting. These other components can be estimated either by sampling from the network or by an analytic formulation after considering a particular model for the structured interaction information, as will be shown in the next section.

### Analytical calculation of triplet probabilities in the two-pool model of network structure

Here we report the analytical calculation of the triplet probabilities in the two-pool model fully described in Methods, section Parametric two-pool model of network structure. Using the probabilities of information-enhancing and information-limiting interactions within and between pools computed in Eq M17, we can compute the probabilities of different network motifs by counting how many of them exist in the network. For example, if we want to compute the probability of the ‘+,+,+’ triplets, we can see that there are four different possible such triplets, as it is shown in Figure S8C, and each of them can be counted and divided by the total number of triplets to compute the probability of *P*_+++_ which can be expressed as follows

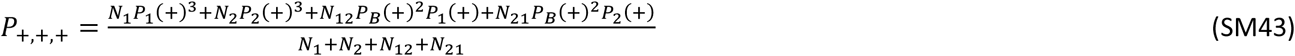

Where 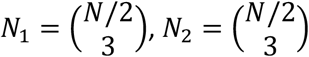 are the number of triplets in each of the pools and 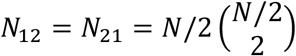 are the number of triplets with two edges between the two pools. Similarly, we can compute the probabilities of the other triplets as follows

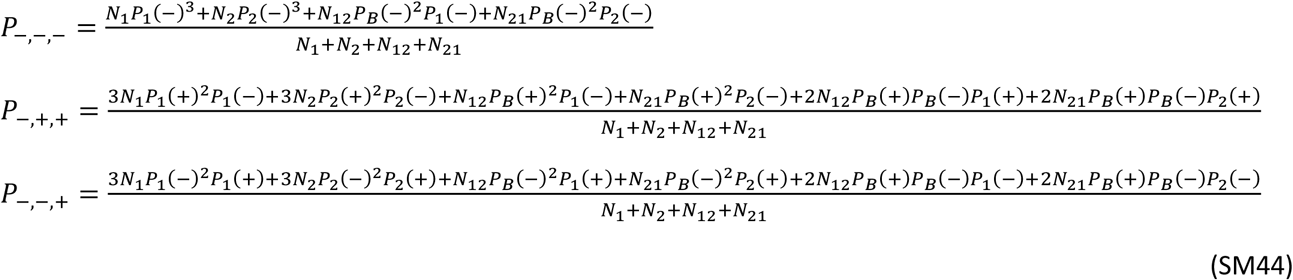

Similar calculations can be performed for quadruplets. Next, we need to compute similar probabilities but for a network with random structure which corresponds to a network with δ*P*_1_ = δ*P*_2_ = 0

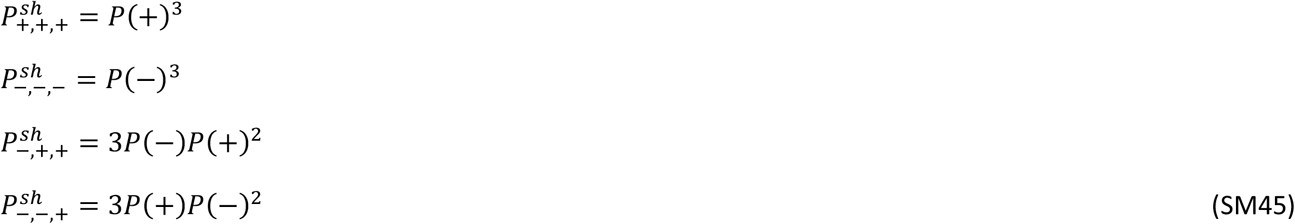

There are four free parameters in these equations. The first parameter is, *N*, the number of neurons in the network (we consider pools of equal size here). The second parameter is δ*P* = *P*(+) − *P*(−) which defines whether the pairwise interactions are more information-enhancing or information-limiting. The third and fourth parameters are the two-pool parametrizations δ*P*_1_ = *P*_1_(+) − *P*(+) and δ*P*_2_ = *P*_2_(+) − *P*(+), which define the existence of structure in the network. For any value of *N* and δ*P*, we can consider a grid 2D grid over δ*P*_1_ and δ*P*_2_ and compute the triplet probabilities on the two-pool model space defined by (δ*P*_1_, δ*P*_2_), as was done in the triplet probabilities presented in Figure 5H, defined as the difference between the real and shuffled probabilities computed using Eq. SM44 and SM45.

### Simplifying the structured interaction information

The structured interaction information can be computed using Eq. SM33 after estimating the real and shuffled interaction terms using Eq. SM32. We can further simplify this calculation to better understand the relationship between the structure of pairwise interactions (which in the case of two-pool network corresponds to being far away from the origin) and the information by assuming that the shuffling the network does not change the expected interaction per motif or IPM, meaning that IPM = IPM^*sh*^. This assumption, on our data, is supported by the fact that single neuron and pairwise interactions do not depend on the type of interaction or triplet. Using this assumption, the structured interaction information, Eq. SM33, can be rewritten as follows

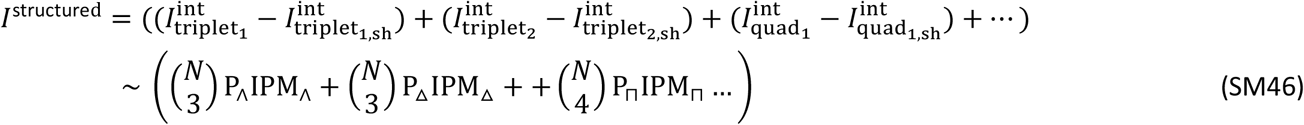

Where the motif probability weights, *P*, are functions of different motif types subtracted from the values corresponding to the random network

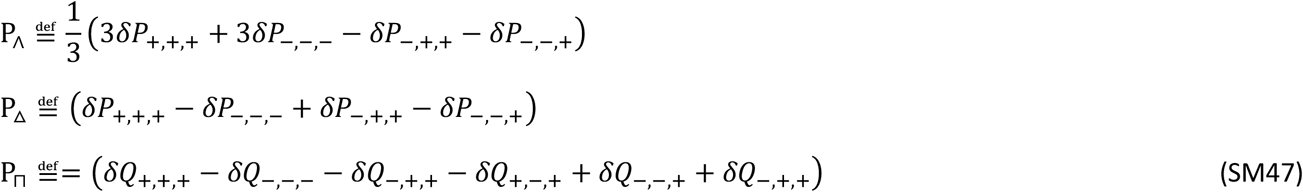

Where residual probabilities δ*P*’s are the difference between the real and shuffled probabilities of each motif type for example 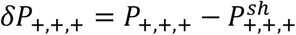.

To compute different components of structured interaction information, we need to first estimate IPM values, using the distributions of single neuron and pairwise interactions and then estimate the probability weight function *P* for each of the open and closed triplets and quadruplets.

Using the assumption of uniformity of IPM’s, the only component that is related to network structure is the residual motif probabilities with respect to the shuffled network which changes by changing the network structure. In fact, we can use the two-pool model to compute the residual probabilities of Eq. SM47, using Eqs. SM44 and SM45 for different values of network size *N* and δ*P* = *P*(+) − *P*(−) as shown in Figure S8D. Interestingly, the results presented in Figure S8D show that for a two-pool network, the quadruplet component is an order of magnitude smaller than the open and closed triplet component and can be ignored in our calculations if we assume the network has a structure similar to a two-pool network. Also, closed, and open triplets have different probability distributions for different networks. The open triplet probability changes more across network sizes, especially for symmetric networks (points close to the diagonal), while the closed triplet probability weight is more sensitive to the change in δ*P* = *P*(+) − *P*(−), especially for asymmetric networks. As can be seen from the left panel of Figure S8D, the open triplet component for larger size networks is much smaller than the closed triplet component on the diagonal, which corresponds to symmetric networks. Given this relationship, and since based on other observations in Figure 6E the structure of the network of neurons projecting to the same target is similar to a symmetric network, we only considered the closed triplet component to make the calculations and visualizations easier. To check the effect of this assumption, we computed the structured interaction information after including the open triplet component. The results are presented in Figure S7D and are very similar to Figure 6E. On the other hand, we also computed the structured interaction information using the IPM uniformity assumption of Eq. SM46, and the results are shown in Figure S7F. Again, as expected, computing the structured interaction information under the assumption that IPM values are uniform between different triplet types shows similar trends to what were computed without using this assumption in Figure 6E because in fact there are small differences between IPM values between different triplet types in our data and uniformity assumption is a good approximation for our networks.

### Additional mathematical details of the calculation of the information expansion

Here we derive the population information Eq. SM1 by approximating the mixture distribution 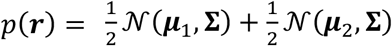 with a Gaussian distribution with mean 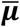 and covariance 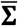. Given the mixture distribution *p*(***r***) we can compute its first and second moment as follows

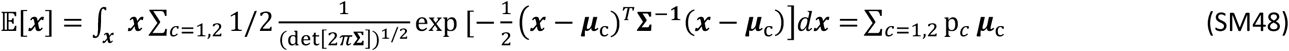

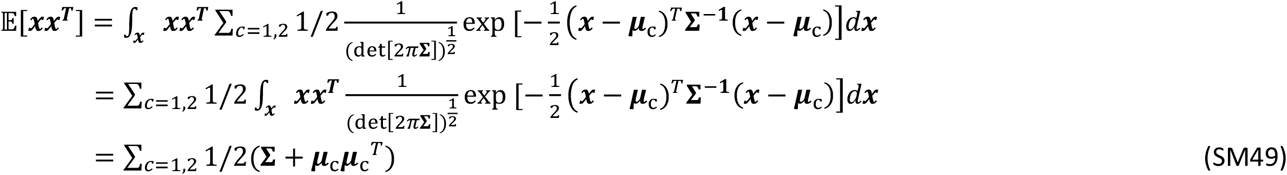

Which means 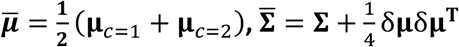.

The population information can then be computed as the difference of the entropies of Gaussian distributions

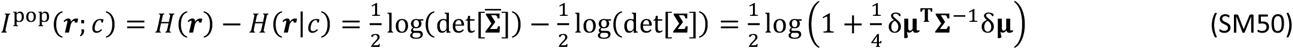

Where we used analytical form of the entropy of a Gaussian distribution, 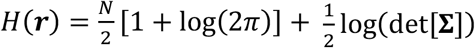. Also, we used the following identity

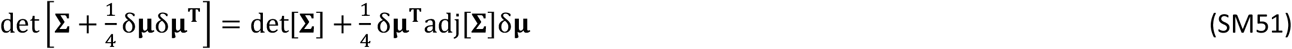

Next, we show that this population information matches up to the second order of *Z* = δμ^*T*^**Σ**^−1^δμ with the population information that we get without the approximation that the mixture distribution of responses is a Gaussian.

We compute the mutual information between population activity and the trial condition as *I*_pop_(***r***; *c*) = *H*(***r***) − *H*(***r***|*c*). Since mutual information is invariant with respect to the shift in responses (*I*_pop_(***r***; *c*) = *I*_pop_(***r*** + ***a***; *c*) for any ***a***), and to simplify the computations, we shift the responses such that the mean activity over two trial conditions become zero 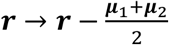. After this shift, the conditional response distributions for *c* = 1,2 will be 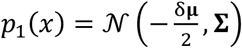 and 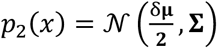 where δμ = μ_2_–μ_1_ is the difference between the means of the distributions corresponding to each trial conditions.

The entropy of the mixture distribution can be computed as follows,

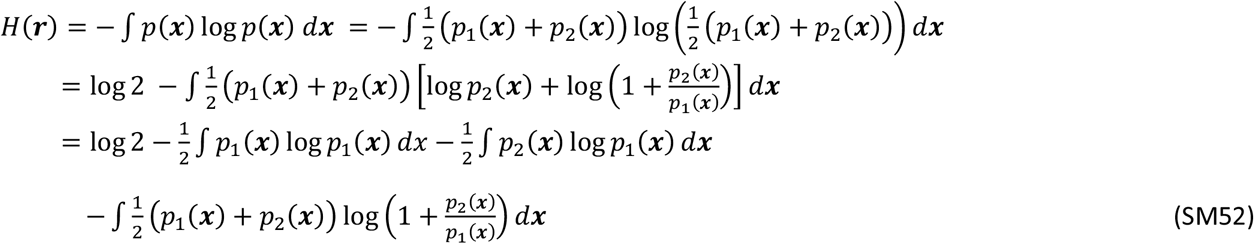

The first two integrals can be analytically computed on the single trial condition Gaussian distributions and the results are given as follows,

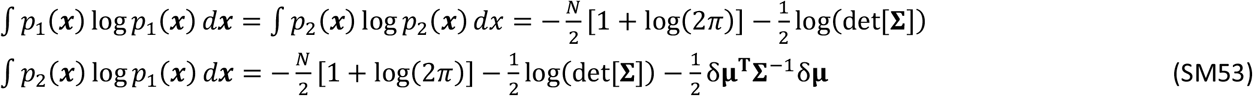

By inserting Eq. SM53 in Eq. SM52, we get

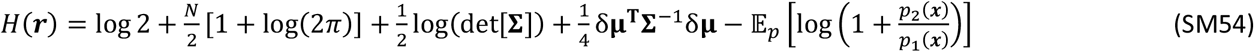

The last term, which is an expectation over the distribution 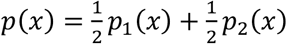, can be written as follows,

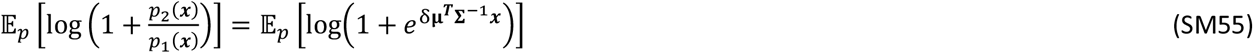

We approximate the argument of the expectation using the Taylor expansion log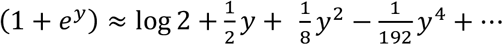 where *y* = δμ^*T*^**Σ**^− 1^***x***,

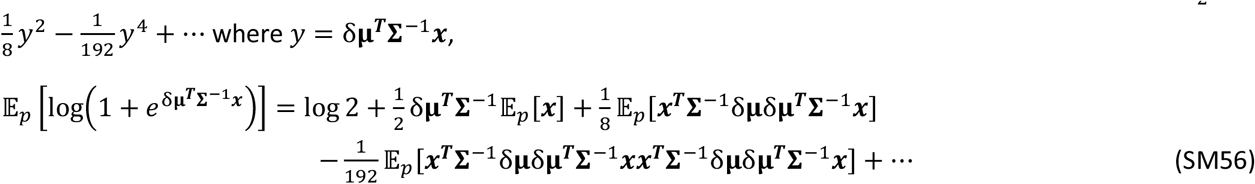

Equation Eq. SM56 transforms the expected value of Eq. SM52 into a sum of expectations of quadratic forms of the vector ***x*** distributed according to *p*(***x***). We computed these quantities as a function of the moments of the single components of the Gaussian mixture distribution. The first moment is zero by definition 𝔼_*p*_[***x***] = 0, and the third and fourth terms of Eq. SM56 have the following expressions:

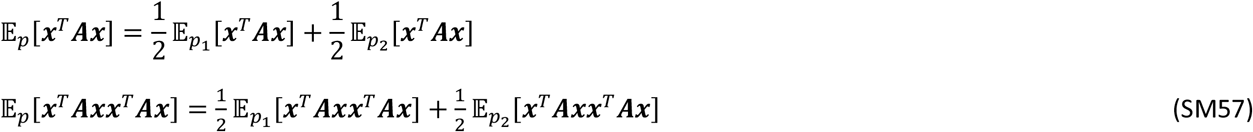

The expectations over *p*_1_(***x***), and *p*_2_(***x***), which are Gaussian distributions have the following expressions,

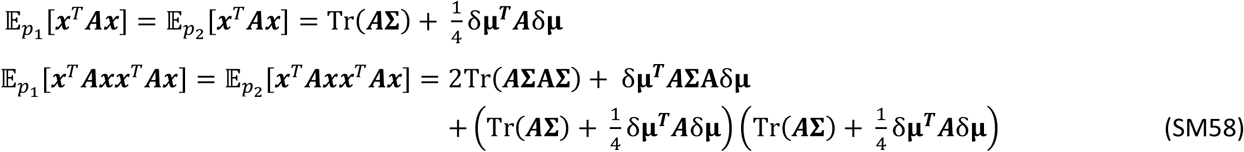

Where, after replacing *A* = **Σ**^−1^δμδμ^*T*^**Σ**^−1^, we get,

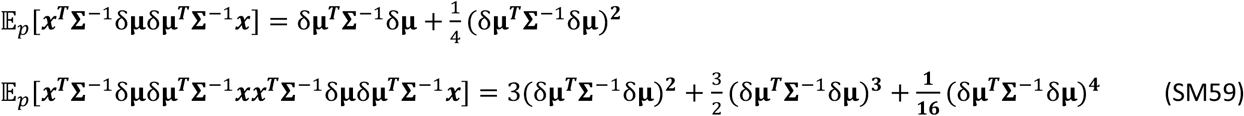

In principle, it is possible to continue to compute these terms up to arbitrary higher order moments in the expansion of Eq. SM53 and SM54, and they will lead to terms with higher powers of δμ and **Σ**^−1^.

Using these results together with the analytical form of the entropy of Gaussian distribution for the response distributions in each trial condition 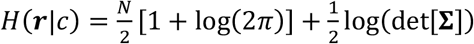, the mutual information of the population can be written as an expansion over increasing orders of the variable *Z* = δμ^*T*^**Σ**^−1^δμ as follows:

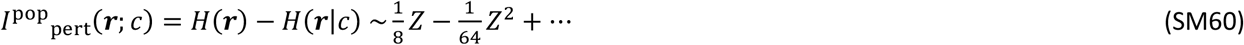

Now if we expand the population information of Eq. SM50 (note that J = *Z*/4), we get:

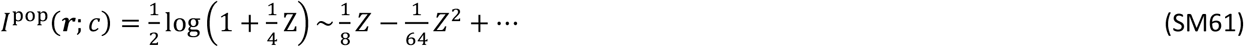

The above equation shows that up to the second order in *Z*, the population information computed after approximating the mixture distribution of responses with a Gaussian matches the population information of the true mixture model.

### Vine copula modelling of neural responses

Here we present the basics of estimating the multivariate density function between a set of variables using the nonparametric vine copula method, which was used to fit neural activity in terms of task and movement variables. The advantage of using the copula is that it makes it possible to estimate a general density function independent from the properties of the single variable or marginal distributions. Moreover, the vine-copula graphical model, which is the multivariate extension of copula, makes it possible to estimate multivariate density functions by breaking the estimation into a sequence of bivariate density estimations, thus avoiding problems related to the curse of dimensionality.

The goal here is to estimate the response probability of neuronal activity *x*_1_ ≡ *n* at any time *x*_2_ ≡ *t* and for any value of other variables possibly modulating the neural activity such as movement variables 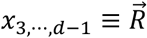, virtual heading direction *x*_*d*_ ≡ θ and trial type Γ = 1 … 8 corresponding to two types of sample cues, two types of test cues and correct and incorrect trials (four trial types for correct trials and four trial types for incorrect trials). The probability of neural response in each trial condition and for each value of behavioral variables can be obtained as follows:

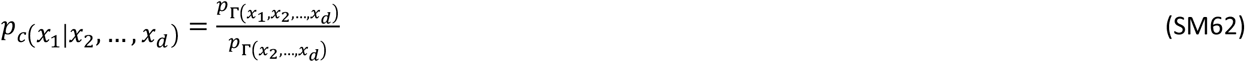

Where 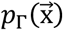 is the joint probability density function of the multivariate variable 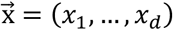 and *p*_Γ_ is a short notation for the conditional density function 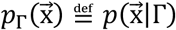, which can be computed simply by using only data from the trial type Γ = 1 … 8.

Estimating high dimensional multivariate probability density functions is a challenging problem because of the curse of dimensionality, and it requires making strong approximations and assumptions about the dependencies between variables. A powerful approach in improving high-dimensional probability density function estimation has been recently developed using the breakdown of the probability density function into a product of marginal probabilities and a component which captures only the dependencies between variables, called the copula

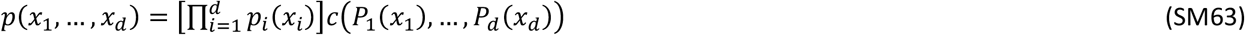

Where *p*_*i*_(X_I_) is the marginal distribution probability of single variable *x*_*i*_ which can be estimated using any parametric or non-parametric method. The copula component of the joint probability density function *c*(*P*_1_(*x*_1_), …, *P*_*d*_(*x*_*d*_)) is the joint probability density of the cumulative density function (CDF) values of the original variables, *u*_*i*_ = P_i_(*x*_*i*_)∼𝒰(0,1) and captures all the high dimensional dependency structure between variables. Since the copula is defined on the d-dimensional unit cube 𝒰(0,1)^*d*^ with uniform marginal distributions, it provides a more robust and reliable numerical density estimation even in two dimensions^34^. Despite its advantages for the original joint density estimation because of the factorization of marginal distributions, in large dimensions estimating the copula suffers from the same curse of dimensionality problems and can be intractable. However, a powerful approach to circumvent this problem has been recently developed combining the advantages of copula and graphical models by decomposing the multivariate copula into a product of bivariate conditional copula densities in a sequential graphical model called *vine-copula*^31,33^. Thus, the complexity of the problem is reduced from a d-dimensional density estimation into a series of bivariate density estimations that are numerically less sensitive to the sample sizes and therefore scalable to large dimensions.

A *d*-dimensional vine is an acyclic graph consisting of a set of (*d* − 1)-dimensional linked trees *T*_*k*_ = (*N*_*k*_, *E*_*k*_),*k* = 1, ⋯, *d* − 1 where *N*_*k*_ and *E*_*k*_ are sets of nodes and edges representing variables and their pairwise dependencies^86^. The organization of the links and edges are such that the edges in *k*^*th*^ tree are nodes in the (*k* − 1)^*th*^ tree and the nodes in *k*^*th*^ tree are connected only if their corresponding edge in the (*k* − 1)^*th*^ tree share a node. A full vine consists of *d*(*d* − 1)/2 edges each of them are bivariate copulas.

The vine graphical models can be used to build multivariate copulas by assigning uniform samples from conditional copulas to the nodes and pairwise copula density functions to the edges. Stated another way, starting from the first tree, each node represents a uniform random variable *u*_*i*_ = *P*_*i*_(*x*_*i*_) sampled from the marginal CDF of *x*_*i*_, which together with another node *u*_*j*_ = *P*_*j*_(*x*_*j*_) shares an edge ε_*ij*_ ∈ *E*_1_ representing the copula density between the two nodes *c*_*ij*_ ≡ *c*(*u*_*i*_, *u*_*j*_). The edge ε_*ij*_ generates a new node in the next tree representing the conditional variable *u*_*j*|*i*_ defined as follows:

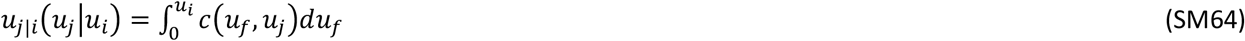

which *u*_*j*|*i*_(*u*_*j*_|*u*_*i*_) is a sample from the conditional copula density function *C*_*ij*_∼𝒰(0,1) corresponding to the probability density function *c*_*ij*_ and is represented by a new node in the second tree. In each step, by going into higher order trees, the set of conditioning variables increases and each node in the *j*^*th*^ tree *u*_*k*|𝒟_ will be conditioned over a set of *j* − 1 dimensional variables 𝒟^87^.

It has been shown^31,33^ that for each vine graph, using the Hammersley–Clifford theorem^88^, the multivariate copula will be equal to the product of the bivariate copulas corresponding to the edges of the vine graph as defined before, as follows:

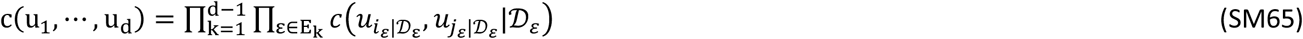

An example of a four-dimensional vine graph, which is called a c-vine, is shown in Figure S9. In principle, the choice of the graphical model for the vine defines the way the full dependency structure is decomposed into a set of bivariate conditional copulas, but the multivariate copula density in Eq. SM65 is invariant to this choice. Practically, computing the bivariate dependencies conditioned over a large dimensional set is still challenging. However, this decomposition is nevertheless useful as an approximation for the dependency structure known as a simplified assumption by considering the bivariate copula in each edge to be constant with respect to the conditioning set of variables 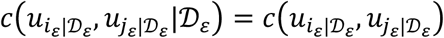. It has been shown, using simulations, that even in case in which the data do not satisfy this assumption, this approximation generates a close estimate of the true dependency and density function^89^. This assumption makes it possible to transform a highly complex multivariate density estimation problem into a set of bivariate density estimations that are simpler and less biased and can be computed even with limited sample sizes. It should be noted that, even though the simplified approximation provides a power estimate for the probability density estimation, there are various ways to extend the computations beyond this approximation. Recent methods on going beyond the simplified assumption have been proposed and shown to increase the power of vine copula method^89^.

### Nonparametric pairwise copula estimation

Using the vine construction, the computation of multivariate copula reduces to the computation of a set of *d*(*d* − 1)/2 bivariate copulas. For each bivariate copula, the probability density function of a set of points on the unit square 𝒰(0,1)^2^ must be computed over a grid covering the whole square. Using these density maps on the grid, we can then compute the values corresponding to each variable in the nodes within the next tree (Figure S9), using Eq. SM65. We can then continue the sequential computation through the higher-order trees until reaching the last tree that has only two variables remaining. The bivariate copula density computation in each stage can be done using either the assumption that the copula belongs to a parametric family of bivariate copulas or using nonparametric approaches that make it possible to achieve the computation without assumptions about the dependency structure of the data. Here, we used a nonparametric copula method (NPC) developed in previous studies^34,73,90–92^ based on local likelihood approach. This method has been shown to provide accurate density and entropy estimations with low biases even in data with small sample sizes, as is often the case for experimental data.

The general kernel estimation of the copula is challenging because we must deal with density estimation in the bounded unit square 𝒰(0,1)^2^, and it is known that kernel computations can be sensitive to boundary effects and asymptotic tail dependencies. To circumvent this problem, we used a trick to transform the data to a new space using a probit function ((u_1_, u_2_) → (Φ^−1^(u_1_), Φ^−1^(u_2_))), where Φ(u) is the CDF of the standard normal distribution, with unbounded support. The probit transformation changes the support from unit square to a space with asymptotically smooth boundaries and the marginal distribution of data transform from uniform (which is the density function of the CDF values) to a standard normal distribution. We then used the local likelihood method to compute the density function of the transformed data in the new space and then transformed back the values to the original space, as follows:

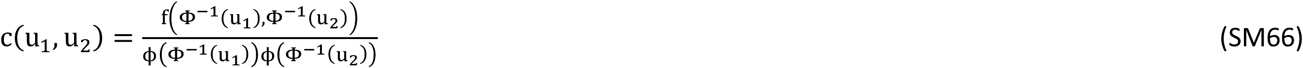

In the above *ϕ* is the standard normal density function and *f* is the copula density function computed in the probit transformed space computed using the local likelihood kernel density estimation, which provides better behavior compared to the normal kernel using the same approach as presented in prior works^34,90,93^. The local likelihood kernel estimation was defined using a local polynomial function 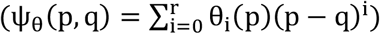 of order *r*. The copula density will be the sum of the weighted naive kernel density and a normalization term ensuring that the density function integrates to one, as follows:

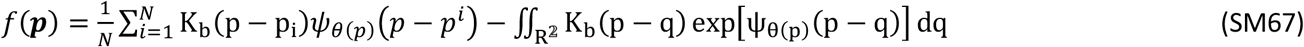

In the above *p* ≡ (*u*_1_, *u*_2_) is a point on the bivariate CDF space, K_b_ is a Gaussian kernel with bandwidth b and θ(*p*) is a parameter. The parameters and the bandwidth of the maximum likelihood estimate of this model were estimated for a training set after expressing the maximum likelihood problem as a cross-validated convex optimization problem using the method explained in^34^. After optimizing the bandwidths, we computed the density function for any test point, using the integrals computed from the training set. The variables that are transformed to the next tree will be the conditional *u*_*j*|*i*_(*u*_*j*_|*u*_*i*_), as computed with Eq. SM64 using the training set, and these variables are computed for the test set. This process continues until the last tree that results in having the full vine-copula density function for any test point.

